# Multifactorial Methods Integrating Haplotype and Epistasis Effects for Genomic Estimation and Prediction of Quantitative Traits

**DOI:** 10.1101/2022.08.06.503033

**Authors:** Yang Da, Zuoxiang Liang, Dzianis Prakapenka

## Abstract

The rapid growth in genomic selection data provides unprecedented opportunities to discover and utilize complex genetic effects for improving phenotypes but methodology is lacking. Epistasis effects are interaction effects and haplotype effects may contain local high-order epistasis effects. Multifactorial methods with SNP, haplotype and epistasis effects up to the third-order are developed to investigate the contributions of global low-order and local high-order epistasis effects to the phenotypic variance and the accuracy of gnomic prediction of quantitative traits. These methods include genomic best linear unbiased prediction (GBLUP) with associated reliability for individuals with and without phenotypic observations including a computationally efficient GBLUP method for large validation populations, and genomic restricted maximum estimation (GREML) of the variance and associated heritability using a combination of EM-REML and AI-REML iterative algorithms. These methods were developed for two models, Model-I with 10 effect types, and Model-II with 13 effect types including intra- and inter-chromosome pairwise epistasis effects that replace the pairwise epistasis effects of Model-I. GREML heritability estimate and GBLUP effect estimate for each effect of an effect type are derived except for third-order epistasis effects. The multifactorial models evaluate each effect type based on the phenotypic values adjusted for the remaining effect types and can use more effect types than separate models of SNP, haplotype and epistasis effects; and provide a methodology capability to evaluate the contributions of complex genetic effects to the phenotypic variance and prediction accuracy, and to discover and utilize complex genetic effects for improving the phenotypes of quantitative traits.

## INTRODUCTION

Genomic estimation and prediction of quantitative traits using single nucleotide polymorphism (SNP) markers and mixed models have become a widely approach for genetic improvement in livestock and crop species. The rapid growth in genomic selection data provides unprecedented opportunities to discover and utilize complex genetic mechanism but methodology and computing tools are lacking for investigating complex genetic mechanisms using the approach of genomic estimation and prediction. The integration of global low-order epistasis effects and local high-order epistasis effects contained in haplotypes for genomic estimation and prediction is a step forward for the discovery and application of complex genetic mechanisms to improve the phenotypes of quantitative traits. The integrated model with multiple types of genetic effects can use more effect types than separate models SNP, haplotype and epistasis effects, and may provide more accurate understanding of each effect type than the separate models due to the use of phenotypic values adjusted for the genetic values of the remaining effect types in the model.

The theory of genetic partition of two-locus genotypic values defines four types of epistasis values, additive × additive (A×A), additive × dominance (A×D), dominance × additive (D×A), and dominance × dominance (D×D) epistasis values by Cockerham and Kempthorne [1, 2]. The Cockerham method defines each epistasis coefficient as the product of the coefficients of the two interacting effects that each can be additive or dominance [1]. This definition of epistasis coefficient is the basis for defining epistasis model matrices in terms of the model matrices of additive and dominance effects. Cockerham also defines a pedigree epistasis relationship as the product between the pedigree additive and dominance relationships [1], and this definition is the theoretical basis for Henderson’s approach to express epistasis relationship matrices as the Hadamard products of the additive and dominance relationship matrices [3].

The Henderson approach of Hadamard products for epistasis relationship matrices was suggested for genomic prediction using epistasis effects by replacing the pedigree additive and dominance relationship matrices with the genomic additive and dominance relationship matrices calculated from SNP markers [4-6]. This genomic version of the Henderson’s Hadamard products calculates genomic epistasis relationship matrices based on the model matrices of SNP additive and dominance effects without creating large epistasis model matrices that can be difficult or impossible to compute. For m SNPs, each pairwise (second-order) epistasis model matrix is (m −1)/2 times as large as the SNP additive or dominance model matrix. For 50,000 SNPs, an epistasis model matrix is nearly 25,000 times as large as the SNP additive or dominance matrix. The calculation of such a large model matrix for pairwise epistasis effects is difficult and the calculation of the model matrices for epistasis effects higher than the second-order is practically impossible for 50,000 or more SNPs. Since genomic additive and dominance relationship matrices are calculated from the SNP additive and dominance model matrices [7-10], the Hadamard products between SNP additive and dominance relationship matrices removes the computing difficulty associated with the large epistasis model matrices for calculating genomic epistasis relationship matrices. However, this genomic version of the pedigree-based epistasis relationship matrices contains intra-locus epistasis effects that is not present in the epistasis model [11]. For this reason, the genomic version of Henderson’s Hadamard product could be described as approximate genomic epistasis relationship matrices (AGERM). Formulations have been developed to obtain the exact genomic epistasis relationship matrices (EGERM) by removing the intra-locus epistasis effects contained in AGERM by modifying Henderson’s Hadamard products without creating the epistasis model matrices [11-14]. The difference between AGERM and EGERM tends to diminish as the number of SNPs increases [13]. A Holstein dataset with 60,671 SNPs showed AGERM and EGERM had the same heritability estimates and the same accuracy of predicting the phenotypic values [15], and the swine dataset with 52,842 SNPs in this manuscript showed the two methods had similar results. However, EGERM required many times of computing time as required by AGERM. The methods in this article allow the use of either AGERM or EGERM, and our computing package of EPIHAP implements both AGERM and EGERM. Henderson’s Hadamard products [3] and hence AGERM are applicable to any order of epistasis effects, and EGERM also has a general formula for any order of epistasis effects [13]. However, limited tests showed that fourth-order global epistasis virtually contributed nothing to the phenotypic variance but generated considerable computing difficulty [16], raising question about the value for global epistasis effects beyond the third-order. Methods of genomic estimation and prediction of global epistasis effects up to the third-order should have a wide-range applications, given that the number of reported epistasis effects lag far behind the number of single-point effects [17-19] even though epistasis effects are important genetic effects [20-22]. In contrast to the computing difficulty and uncertain impact of global high-order epistasis effects beyond the third-order, local high-order epistasis effects in haplotypes with many SNPs were responsible for the increased accuracy of predicting phenotypic values of certain traits. For examples, a haplotype model with 12 SNPs per haplotype block had the best prediction accuracy for low density lipoproteins in a human population [23], a haplotype model with 500 Kb haplotype blocks that on average had 105 SNPs per block had the best prediction accuracy for average daily gain in a swine population [24], and a haplotype model with 15 SNPs per haplotype block had the best prediction accuracy in a wheat study [25]. The integration of haplotype and epistasis effects provides an approach to investigate the contributions of global low-order epistasis effects and local high-order epistasis effects to the phenotypic variance and the accuracy of genomic prediction under the same model.

An epistasis GWAS in Holstein cattle showed that intra- and inter-chromosome epistasis effects affected different traits differently, e.g., daughter pregnancy rate was mostly affected by inter-chromosome epistasis effects whereas milk production traits were mostly affected by intra-chromosome epistasis effects [26]. Genomic heritability estimates of intra- and inter-chromosome heritabilities for daughter pregnancy rate using methods in this article showed that intra-chromosome A×A heritability was 0.031, and inter-chromosome A×A heritability 0.178 [15], consistent with the GWAS results of 21% intra-chromosome and 79% inter-chromosome A×A effects among the top 33,552 pairs of A×A effects in the GWAS study [26]. Therefore, dividing pairwise epistasis effects into intra- and inter-chromosome epistasis effects for genomic prediction and estimation allows the investigation of the contributions of intra- and inter-chromosome pairwise epistasis effects to the phenotypic variance and prediction accuracy.

The purpose of the multifactorial model in this article is to integrate haplotype effects and epistasis effects up to the third-order for genomic estimation and prediction of quantitative traits, to provide a general and flexible methodology framework for genomic prediction and estimation using complex genetic mechanisms, and to provide methodology details of the EPIHAP computer package that implements the integration of haplotype and epistasis effects [15, 16]. The multifactorial model has the advantage of using more effect types and assessing each effect type based on the phenotypic values adjusted for all remaining effect types over separate SNP, haplotype and epistasis models. We hypothesize that some traits involve only a small number of the effect types, some traits are more complex and involve more effect types, global low-order epistasis are more important than local high-order epistasis effects of haplotypes for some traits whereas the reverse is true for some other traits, and some traits may be affected by both global low-order and local high-order epistasis effects. The methodology in this article will provide an approach to evaluate these hypotheses, facilitate the discovery and utilization of global low-order and local high-order epistasis effects relevant to the phenotypic variance and prediction accuracy of each trait, and obtain new knowledge of complex genetic mechanisms underlying quantitative traits.

## METHODS

### Quantitative genetics (QG) model with SNP, haplotype and epistasis effects and values

The mixed model with single-SNP additive and dominance effects, haplotype additive effects and pairwise SNP epistasis effects in this article is based on the quantitative genetics (QG) model resulting from the genetic partition of single-SNP genotypic values [9, 10], haplotype genotypic values [27], and pairwise genotypic values [1]. An advantage of this QG model is the readily available quantitative genetics interpretations of SNP additive and dominance effects, values and variances; haplotype additive effects, values and variances; epistasis effects, values and variances; and the corresponding SNP, haplotype and epistasis heritability estimates. Two QG models are developed: Model-I with 10 effect types including SNP additive and dominance effects, haplotype additive effects, and epistasis effects up to the third-order; and Model-II with 13 effect types resulting from replacing the pairwise epistasis effects of Model-I with intra- and inter-chromosome epistasis effects. Detailed descriptions of the effects, values, model matrices and the coding of the model matrices as well as the precise definition of each term in the two QG models are provided in **Supplementary Text S1 and Table S1**. With these precise definitions of genetic effects, values and model matrices in the QG models, a concise multifactorial QG model covering both Model-I and Model-II is established, i.e.:

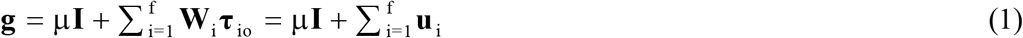

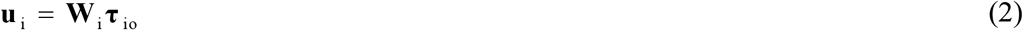

where **τ**_io_ = genetic effects of the i^th^ effect type from the original QG model based on genetic partition, **W**_i_ = model matrix of **τ**_io_, **u**_i_ = genetic values of the i^th^ effect type from the original QG model, and f = number of effect types. For Model-I, subscripts i=1,&,10 represent SNP additive (A), SNP dominance (D), haplotype additive, A×A, A×D, D×D, A×A×A, A×A×D, A×D×D and D×D×D effects sequentially. For Model-II, subscripts i=1,&,13 represent SNP additive, SNP dominance, haplotype additive, intra-chromosome A×A, intra-chromosome A×D, intra-chromosome D×D, inter-chromosome A×A, inter-chromosome A×D, inter-chromosome D×D, A×A×A, A×A×D, A×D×D and D×D×D effects sequentially. The variance-covariance matrix of the genetic values of Equations 1 and 2 is:

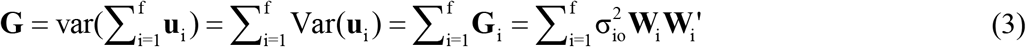

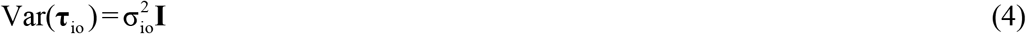

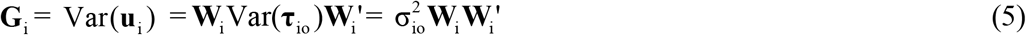

where 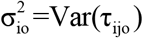 = genetic variance of the i^th^ effect type under the original QG model common to all individuals (all j values). Note that **W**_i_ **W**_i_′ is not a genomic relationship matrix but is the primary information for calculating each genomic relationship matrix. The structure of the **G** matrix of Equation 3 assumes independence between the genetic values of different effect types. However, the GBLUP values of different effect types using the **G** matrix of Equation 3 could be correlated. Under the Hardy-Weinberg equilibrium (HWE) and LE assumptions, additive, dominance and epistasis effects are independent of each other [1, 2]. For genome-wide SNPs, the LE assumption generally does not hold for closely linked loci, and nonzero Hardy-Weinberg disequilibrium (HWD) may exist numerically. These and other unknown factors in real data may result in the existence of correlations between different effect types. Haplotype additive values are correlated with SNP additive effects because a haplotype additive value is the sum of all SNP additive values and an epistasis value within the haplotype block plus a potential haplotype loss [28]. In two recent haplotype studies for genomic prediction, the integration of SNP and haplotype effects increased the prediction accuracy for four of the seven traits in the human study [23] and for three of the eight traits in the swine study [24], showing that SNP and haplotype additive values compensated each other for prediction accuracy and that the correlation between SNP and haplotype additive values were incomplete for those traits. The correlation between haplotype and epistasis values can be complex: the correlation should be nonexistent if the A×A values are inter-chromosome A×A values or intra-chromosome A×A values involving distal SNPs not covered by the haplotypes, but the correlation could be strong if the A×A values are intra-chromosome A×A values involving proximal SNPs covered by the haplotypes.

### The reparametrized and equivalent QG model for genomic estimation and prediction

Genomic relationship matrices will be used for genomic estimation and prediction, and the use of genomic relationship matrices results in a reparametrized and equivalent model of the original QG model for genetic values, to be referred to as the RE-QG model, where “reparametrized” refers to the reparameterization of the genetic effects, model matrix and genetic variance of each effect type; and “equivalent” refers to the requirement of the same first and second moments for the original QG model (Equations 1-5) and the RE-QG model described below. This RE-QG model of genetic values can be expressed as:

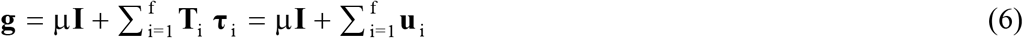

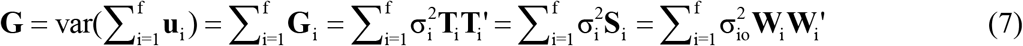

where

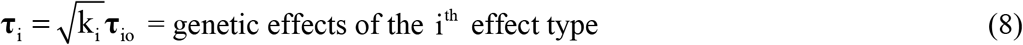

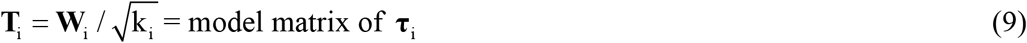

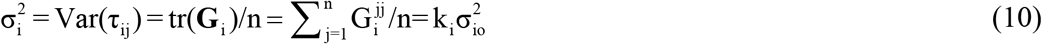

= variance of the genetic effects of the i^th^ effect type common to all individuals

= average variance of all individuals for the genetic values of the i^th^ effect type

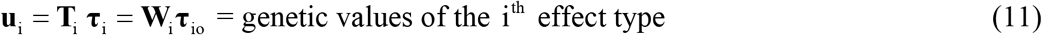

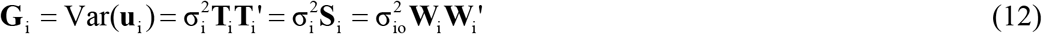

= variance-covariance matrix of the genetic values of the i^th^ effect type

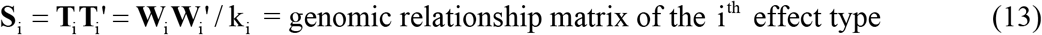

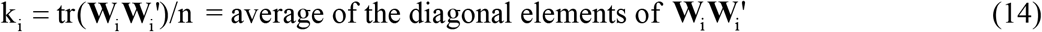

Equations 8-10 are the reparametrization of the genetic effects, model matrices and genetic variances of the original QG model, whereas Equations 11 and 12 show the genetic values and the variance-covariance matrix of the genetic values are the same under the RE-QG and QG models. In Equation 10, 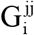 = the genetic variance of the j^th^ individual for the i^th^ effect type = the j^th^ diagonal element of the **G**_i_ matrix defined by Equation 12. The k_i_ formula of Equation 14 as the average of the diagonal elements of **W**_i_ **W**_i_’ was originally proposed for genomic additive relationships [8], and was used for genomic dominance relationships [9, 10], haplotype additive genomic relationships [27], and pairwise epistasis genomic relationships [6]. The need of this RE-QG model is due to the use of the genomic relationship matrices (e.g., Equation 13), because the QG model does not contain genomic relationship matrices (Equation 3). Detailed notations of the QG model of Equations 1-5 in reference to the RE-QG model described by Equations 6-14 are summarized in **Supplementary Table S1**.

The formula of genomic relationship matrix (**S**_i_ of Equation 13) is based on the model matrix of each effect type and can be difficult or impossible to compute if epistasis model matrices are used. This computing difficulty of epistasis model matrices is removed by calculating **S**_i_ based on the model matrices of SNP additive and dominance effects without creating the epistasis model matrices using either AGERM or EGERM. AGERM refers the genomic version of Henderson’s Hadamard products between pedigree additive and dominance relationship matrices [3] with the pedigree additive and dominance relationship matrices replaced by the genomic additive and dominance relationship matrices [4-6]. AGERM contains intra-locus epistasis that should not exist [11] and EGERM removes intra-locus epistasis from AGERM based on products between genomic additive and dominance relationship matrices [11, 13].

The QG and RE-QG models have the same prediction accuracy due to the equivalence between these two models. The genetic values (**u**_i_, Equations 1 and 6) and the variance-covariance matrix of the genetic values (**G**_i_, Equations 5 and 12) under the QG and RE-QG models are identical, although these two equations have different expressions for the genetic effects and model matrices. Consequently, the QG model without using genomic relationship matrices and the RE-QG model using genomic relationship matrices have identical accuracy of genomic prediction. The choice of the k_i_ formula for defining the genomic relationship matrix does not affect the accuracy of genomic prediction but affects the interpretation and application of the genetic variance and genomic relationships for each effect type. Since the interpretation of each genetic variance is a focus whereas the interpretation of the genomic relationships is not a focus in this study, the interpretation of the genetic variance and associated heritability is the consideration in choosing the k_i_ formula of Equation 14.

The RE-QG model using genomic relationships (Equations 6-14) has two major advantages over the QG model without using genomic relationship matrices (Equations 1-5) although the two models have the same prediction accuracy. First, the use of genomic relationships, originally proposed for genomic additive relationships [7], provides a genomic version of the traditional theory and methods of best linear unbiased prediction (BLUP) that uses pedigree relationships, and this genomic version can utilize a wealth of BLUP-based theory, methods and computing strategies. Second, the genetic variance of the genetic effects of each effect type under the RE-QG model can be used for estimating genomic heritability whereas the genetic variance of the genetic effects under the QG model cannot be used for estimating genomic heritability. With the k_i_ value defined by Equation 14, The variance of the genetic effects of the i^th^ effect type, 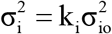 (Equation 10), has the unique interpretation as the average variance of the genotypic values of all individuals and is a common variance to all individuals. Moreover, 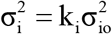 is unaffected by the number of levels for each effect type, unless the number of levels such as the number of SNPs is too small to provide sufficient coverage of the genome [9, 23, 29]. In contrast, the QG model does not have a method to estimate genetic variance components for calculating genomic heritabilities, because 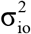 is an inverse function of the number of effect levels. As the number of effect levels such as the number of SNPs increases or decreases, the value of each element in **W**_i_ **W**_i_’ changes in the same direction and the 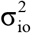 estimate changes in the opposite direction, i.e., as the number of effect levels increases or decreases, 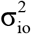 decreases or increases. Consequently, the 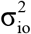 estimate does not have a unique interpretation and cannot be used for estimating genomic heritability [9]. Moreover, the variance of the genetic value of an individual 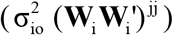 cannot be used for calculating genomic heritability because of the individual specificity of the (**WW**_i_’) ^jj^ values, as shown as follows.

The exact relationship between the genetic variance for the i^th^ effect type of the j^th^ individual under the RE-QG model and the QG model can be described based on the **G**_i_ matrix defined by Equation 12, i.e.:

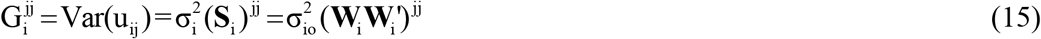

where 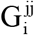 = the j^th^ diagonal element of the **G**_i_ matrix defined by Equation 12 = the genetic variance of the j^th^ individual for the genotypic value of the i^th^ effect type, and u_ij_ = the j^th^ element of **u**_i_ defined by Equation 11. Equation 15 shows that different individuals do not have a common variance of the genetic values 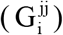 unless all diagonal elements of **S** or **W**_i_**W**_i_’ are identical, which could not happen with genome-wide SNP data in the absence of identical twins because genome-wide SNPs have a high degree of individual specificity. Consequently, 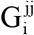 is not a common variance to all individuals and cannot be used for calculating the genomic heritability of the i^th^ effect type. In contrast, 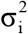 of Equation 10 under the RE-QG model as the average variance of the genotypic values of all individuals is common to all individuals and can be used for calculating the heritability of each effect type. For the example of Model-I, the exact genetic interpretation of 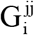 is: 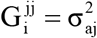 = the variance of the genomic additive (breeding) value of the j^th^ individual for 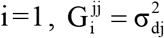 = the variance of the genomic dominance value of the j^th^ individual for 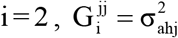 = the variance of the genomic haplotype additive value of the j^th^ individual for 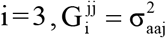 = the variance of the A×A value of the j^th^ individual for 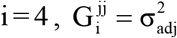 = the variance of the A×D value of the j^th^ individual for 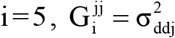 = the variance of the D×D value of the j^th^ individual for 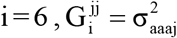 = the variance of the A×A×A value of the j^th^ individual for 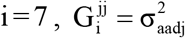 = the variance of the A×A×D value of the j^th^ individual for 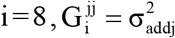 = the variance of the A×D×D value of the j^th^ individual for i= 9, and 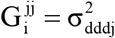 = the variance of the D×D×D value of the j^th^ individual for i =10. These genetic interpretations along with those for intra- and inter-chromosome pairwise epistasis effects of Model-II under the QG and RE-QG models are summarized in **Supplementary Table S1**.

## RESULTS AND DISCUSSION

### The multifactorial model of phenotypic values

Based on the RE-QG model of Equations 6-14, the multifactorial model for phenotypic values is:

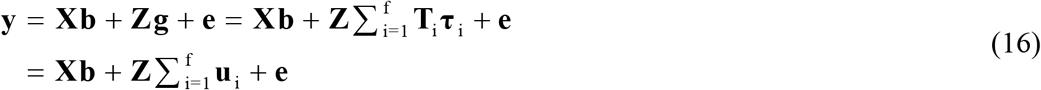

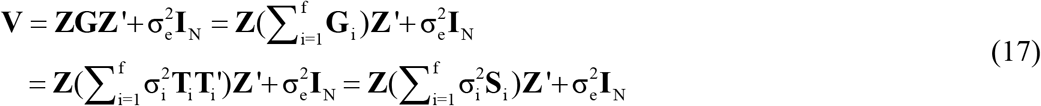

where **y** = N×1 column vector of phenotypic observations, **Z** = N×n incidence matrix allocating phenotypic observations to each individual = identity matrix for one observation per individual (N = n), N = number of observations, n = number of individuals, **b** = c×1 column vector of fixed effects such as heard-year-season in dairy cattle, c = number of fixed effects, **X** = N×c model matrix of **b, e** = N×1 column vector of random residuals, 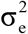 = residual variance, and 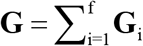 (Equation 7). The phenotypic values (**y**) are assumed to follow a normal distribution with mean **Xb** and variance-covariance matrix of **V**. The methods described below for genomic estimation and prediction are based on the conditional expectation (CE) method, which is more efficient computationally than the methods based on mixed model equations (MME) when the number of genetic effects is greater than the number of individuals [9, 27].

For Model-I with 10 effect types, the genomic epistasis relationship matrices can be calculated using either EGERM or AGERM. However, EGERM or AGERM did not consider intra- and inter-chromosome genomic epistasis relationship matrices that are required by Model-II with 13 effect types. This research derives intra- and inter-chromosome genomic epistasis relationship matrices for both EGERM and AGERM.

### Intra- and inter-chromosome genomic epistasis relationship matrices

The main derivation of the intra- and inter-chromosome genomic epistasis relationship matrices is the partition of the numerator of a genomic epistasis relationship matrix into intra- and inter-chromosome numerators. The first step is to derive the intra-chromosome numerator, and the second step is to derive the inter-chromosome numerator as the difference between whole-genome numerator and the intra-chromosome numerator. The last step is to divide the intra-chromosome numerator by the average of the diagonal elements of the intra-chromosome numerator, and to divide the inter-chromosome numerator by the average of the diagonal elements of the inter-chromosome numerator. Using this procedure, intra- and inter-chromosome epistasis relationship matrices were derived for both EGERM and AGERM (**Supplementary Text 1**).

### Genomic best linear unbiased prediction (GBLUP) and reliability

Based on the multifactorial genetic model of Equations 16 and 17, the GBLUP of the genetic values of the i^th^ effect type (**û**_i_) and the best linear unbiased estimator (BLUE) or generalized least squares (GLS) estimator of fixed effect 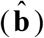 are:

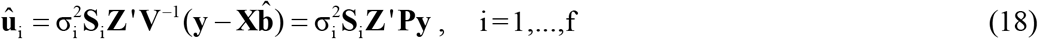

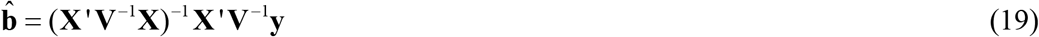

where **P** = **V**^−1^ − **V**^−1^**X** (**X**′ **V**^−1^**X**)^−^ **X**′ **V**^−1^. The GBLUP of total genetic values of the n individuals is the summation of all types of genetic values:

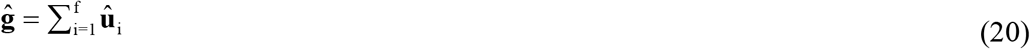

Reliability of GBLUP is the squared correlation between the GBLUP of a type of genetic values and the unobservable true genetic values being predicted by the GBLUP. The expected accuracy of predicting the genetic values by the GBLUP is the square root of reliability, or the correlation between the GBLUP of a type of genetic effects and the unobservable true genetic effects being predicted by the GBLUP. In the absence of validation studies for observed prediction accuracy, reliability or the expected prediction accuracy is the measure of prediction accuracy of the GBLUP.

The reliability of the GBLUP of the total genetic value (Equation 20) of the

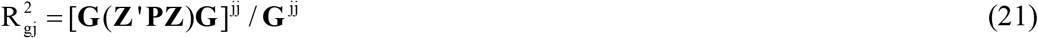

where 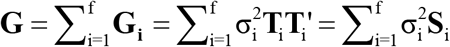 (Equation (7), 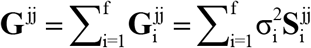, and subscript or superscript jj denotes the j^th^ diagonal element. The reliability formula for any or a combination of genetic values can be readily derived from Equation 21, e.g., the reliability of **û**_3_ (GBLUP of haplotype additive values) is obtained from Equation 21 by deleting all terms except **G**_3_ (**Z**′ **PZ**)**G**_3_ in the numerator and 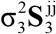 in the denominator, with changes in the **V** and **P** matrices accordingly.

### Calculation of GBLUP and reliability for individuals with and without phenotypic observations separately

Two strategies are available for calculating GBLUP and reliability of Equations 20 and 21. Strategy-1 is a one-step strategy that include all individuals with and without phenotypic observations in the same system of equations so that GBLUP and reliability are calculated simultaneously for all individuals. This strategy essentially augments the mixed model for individuals with phenotypic observations with a set of null equations consists of ‘0’s but uses each genomic relationship matrix for all individuals, and these null equations and the use of the relationship matrix for all individuals do not affect the GBLUP, reliability and heritability of individuals with phenotypic observations. The advantage of this one-step strategy is the simplicity of data preparation. For example, for a k-fold cross validation study, the phenotypic input file only needs to have k columns of the trait observations, with one column for each validation where the phenotypic observations for the validation individuals are set as ‘missing’ and the **X** and **Z** model matrices for the ‘missing’ observations are set to zero. With this strategy, the genotypic data needs to be processed only once. As the number of traits increases for validation studies, this one-step strategy becomes more appealing due to the savings in data preparation work. This strategy has been implemented in our computing tools of GVCBLUP [30], GVCHAP [31] and EPIHAP [15, 16]. However, when the number of validation individuals or individuals without phenotypic values is large, each genomic relationship matrix (**S**_i_ matrix) is large and the one-step strategy becomes more difficult as the number of individuals increases.

For large numbers of individuals without phenotypic observations, calculating GBLUP for individuals with and without phenotypic values separately is more efficient computationally than calculating GBLUP for all individuals in the same system of equations by applying Henderson’s BLUP for animals without phenotypic observations [32] to GBLUP. Let n_1_ = number of individuals with phenotypic observations, n_0_ = number of individuals without phenotypic observations, n = n_1_ + n_0_, and let the **S**_i_ matrix be partitioned as:

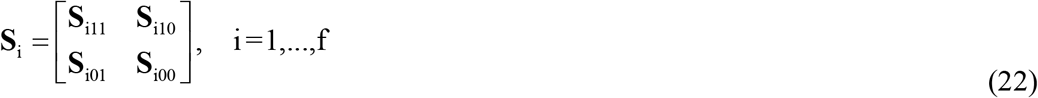

where **S**_i11_ = n_1_ × n_1_ genomic relationship matrix of the genetic values of the i^th^ effect type for individuals with phenotypic observations, **S**_i01_ = n _0_ × n_1_ = genomic relationship matrix of the genetic values of the i^th^ effect type between individuals without phenotypic observations and individuals with phenotypic observations, **S**_i10_ = **S**_i01_′ = n_1_ × n _0_ = genomic relationship matrix between individuals with phenotypic observations and individuals without phenotypic observations, and **S**_i00_ = n_0_ × n_0_ genomic relationship matrix of the genetic values of the i^th^ effect type for individuals without phenotypic observations. In Equations 16 and 17, **y** = **y**_1_, and the **Z** matrix needs to be changed to **Z** = [**Z**_1_ **0**], the **u**_i_ vector partitioned as **u**_i_ = [**u**_i1_′ **u**_i0_′]′, and the **g** vector partitioned as **g** = [**g**_1_′ **g**_0_′], where **Z**_1_ = N × n_1_ incidence matrix allocating phenotypic observations to individuals with phenotypic observations, **0** = N × n _0_ incidence matrix with elements ‘0’ connecting phenotypic observations to individuals without phenotypic observations. With these changes and Equation 22, the **V** matrix of Equation 17 can be re-written as:

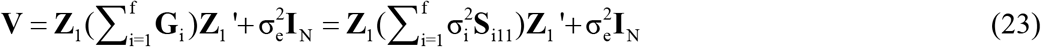

and the GBLUP and reliability for individuals with and without phenotypic observations can be calculated as:

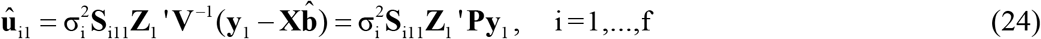

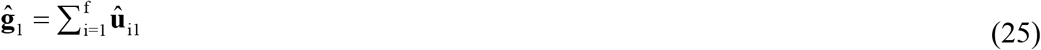

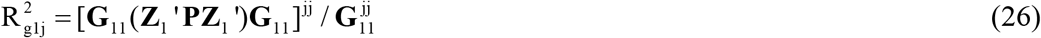

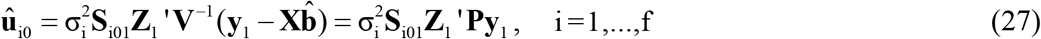

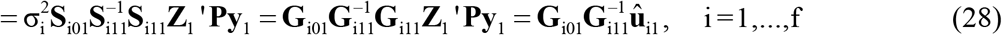

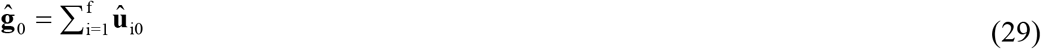

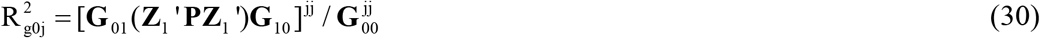

where **û**_i1_ = n ×1 column vector of the GBLUP of the genetic values of the i^th^ effect type for individuals with phenotypic observations, **ĝ**_1_ = n_1_ ×1 column vector of the GBLUP of the total genetic values for individuals with phenotypic observations,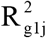 = reliability for the j^th^ individuals with phenotypic observations, **û**_i0_ = n _0_ ×1 column vector of the GBLUP of the genetic values of the i^th^ effect type for individuals without phenotypic observations **ĝ**_0_ = n _0_ ×1 column vector of the GBLUP of the total genetic values for individuals without phenotypic observations, 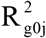 = reliability for the j^th^ individuals without phenotypic observations, 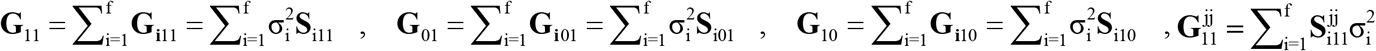, and 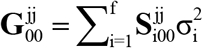.

Equations 27 and 28 yield identical results if 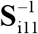 exists. However, when the number of individuals is greater than the number of effect levels such as the number of SNPs, 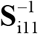 in Equation 28 does not exist and Equation 27 still can calculate the GBLUP. The usefulness of Equation 28 is showing the GBLUP of individuals without phenotypic observations is the regression of the genetic values of individuals without phenotypic observations on the genetic values of individuals with phenotypic observations. The advantage of Equation 27 is that it does not calculate 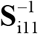 and hence is unaffected by the singularity of **S**_i11_. Therefore, Equation 27 is recommended for calculating GBLUP for individuals without phenotypic observation when the number of such individuals is large. The GBLUP calculations of Equations 24, 27 and 28 do not involve the genomic relationship matrix among individuals without phenotypic observations **S**_i00_, which is much larger than **S**_i11_ when n_1_ is much larger than n _0_. The reliability calculation for individuals without phenotypic observations (Equation 30) only uses the diagonal elements of **S**_i00_, not the entire **S**_i00_.

### Advantage of integrated model over separate models

The multifactorial model of Equations 16 and 17 integrating SNP, haplotype and epistasis effects have the advantage of using more effect types and assessing each effect type based on the phenotypic values adjusted for all remaining effect types over separate models for SNP, haplotype and epistasis effects that do not have a mechanism to adjust for effect types not in the model and each uses a smaller number of genetic effects in the model.

This advantage of the multifactorial model assessing each effect type based on the phenotypic values adjusted for all remaining effect types can be shown using the MME version of the GBLUP for the i^th^ effect type, i.e.,

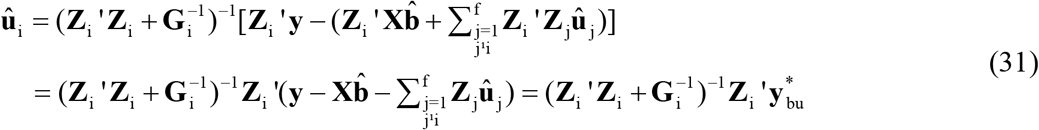

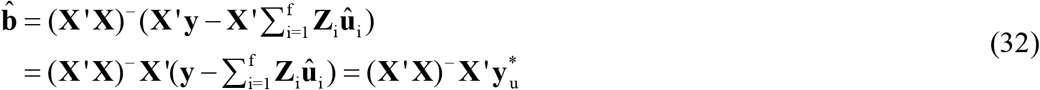

where 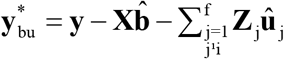 = phenotypic observations adjusted for the fixed effects and all random genetic values except those of **û**1_i_, 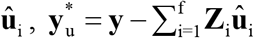 = phenotypic observations adjusted for all random genetic values, and (**X**′ **X**)^−^ is a generalized inverse of **X**′ **X**. Equation 31 shows the MME version of **û**_i_ uses the phenotypic values adjusted for the GBLUP of all other effect types in the model. Since the MME version of **û**_i_ (Equation 31) and 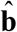 (Equation 32) are identical to the CE version of **û**_i_ (Equation 18) and 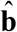(Equation 19), the CE version of **û**_i_ (Equation 18) uses the phenotypic values adjusted for the GBLUP of all other effect types in the model even though the CE version does not do such adjustments explicitly.

### Genomic restricted maximum estimation (GREML) of variances and heritabilities

The estimation of variance components uses GREML and a combination of EM-REML and AI-REML algorithms of iterative solutions. EM-REML is slow but converges whereas AI-REML is fast but fails for zero heritability estimates. In our GVCBLUP and GVCHAP computing packages that implement these two algorithms [30, 31], EM-REML is used automatically when AI-REML fails. The EM-REML iterative algorithm for the multifactorial model of Equations 16 and 17 is:

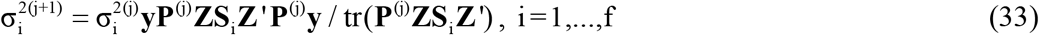

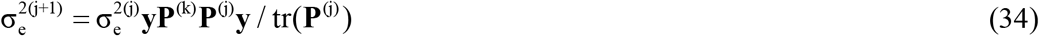

where j = iteration number. The AI-REML iterative algorithm is an extension of the early formulations [33, 34] to the multifactorial model of Equations 16 and 17:

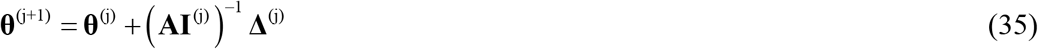

where 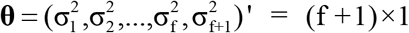 column vector of variance-covariance components, 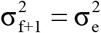 = residual variance, **Δ** = (Δ_1_, Δ_2_,&, Δ_f_, Δ_f+1_)′ = (f +1)×1 column vector of the partial derivatives of the log residual likelihood function with respect to each variance component, and j = iteration number. A typical term in **Δ** (Δ_i_) and a typical term in **AI** (AI_ik_) are:

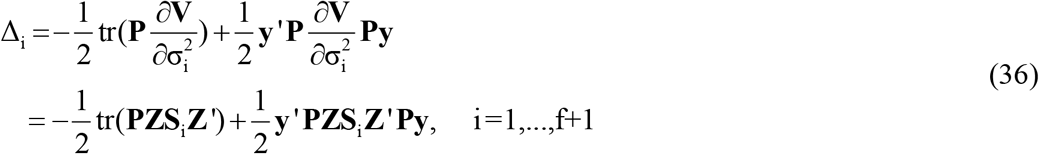

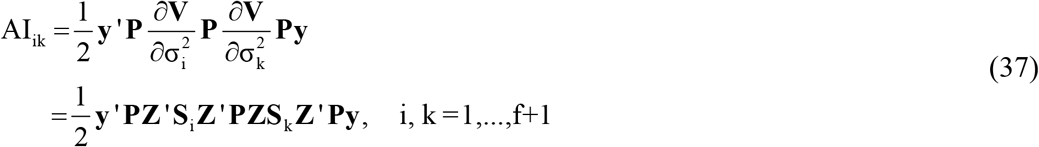

where **S**_f+1_ = **I** _N_. For the full Model-I or Model-II, some effect types inevitably may have zero variances. In those cases, AI-REML (Equations 35-37) fails, and EM-REML (Equations 33 and 34) still converges although slow convergence rate can be expected for the full Model-I or Model-II. Once the effect types with zero variances are removed from the model, AI-REML converges, and fast convergence rate can be expected. The estimate of the genomic heritability for each type of genetic effects 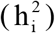 and the total heritability of all types of genetic effects (H^2^) are:

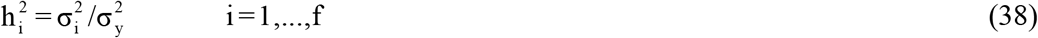

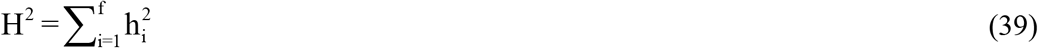

where 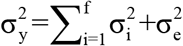 = phenotypic variance.

The heritability estimates of Equation 38 can be used for model selection by removing effect types with heritability estimates below a user determined threshold value from the prediction model. Since different traits may have different genetic architectures, we hypothesize that some traits may involve only a small number of the effect types and some traits are more complex and involve more effect types, global epistasis may be more important than local high-order epistasis effects of haplotypes for some traits whereas the reverse may be true for other traits, and some traits may be affected by both global high-order and local high-order epistasis effects. The heritability estimates from Equation 37 provide an approach to evaluate these hypotheses and identify effect types relevant to the phenotypic variance whereas the total heritability of Equation 38 provides an estimate of the total genetic contribution to the phenotypic variance. In addition to the use of heritability estimates, prediction accuracy based on GBLUP can be used for model selection by requiring a threshold accuracy level for the effect type to be included in the prediction model, e.g., we identified the A + A×A model to have the same accuracy of predicting the phenotypic values of daughter pregnancy rate as the full Model-I in U.S. Holstein cows [15].

### Estimation of pairwise epistasis effect and heritability

The heritability of a SNP, haplotype block or pairwise epistasis effect is the contribution of the genetic effect to the phenotypic variance and is also the contribution to the heritability of the effect type, and is estimated through the GBLUP of the corresponding genetic effects. These heritability estimates can be used to identify genome locations with large contributions to the phenotypic variance. The estimation of pairwise epistasis effects and heritability is the most demanding computing because the pairwise epistasis model matrices must be creased and are no longer avoidable. Estimating the effects and heritabilities for third-order epistasis effects is computationally unfeasible and is not considered. GBLUP of SNP, haplotype and pairwise epistasis effects of Model-I (**Supplementary Table S1**) are calculated as:

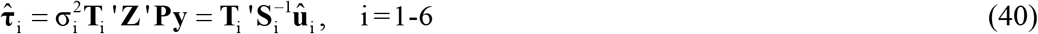

where 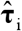 is the m×1 column vector of SNP additive effects for i= 1, or SNP dominance effects for i= 2; or b×1 column vector of haplotype additive effects for i= 3; or 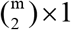 column vector of A×A epistasis effects for i= 4, or 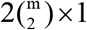 column vector of A×D epistasis effects for i= 5, or 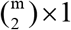 column vector of D×D epistasis effects for i= 6. For i= 5, the order of A×D and D×A effects is determined by the order of the model matrices of those effects, i.e., 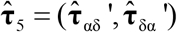 if **T**_5_ = (**T**_αδ_, **T**_δα_), or 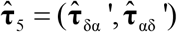 if **T**_5_ = (**T**_δα_, **T**_αδ_). The heritability of the j^th^ effect of the i^th^ effect type 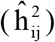 is estimated as a faction of the genomic heritability of the i^th^ effect type 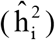:

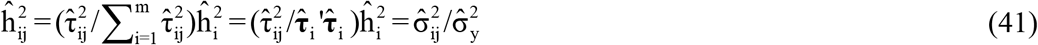

where 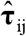 = the j^th^ effect of 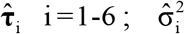 = estimated variance of the i^th^ effect type; 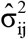 = estimated variance of the j^th^ effect of the i^th^ effect type; 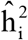 = the genomic heritability of the i^th^ effect type defined by Equation (52). For proving Equation 57, 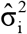 and 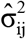 can be formulated based on the method of mixed model equations (MME), i.e.,

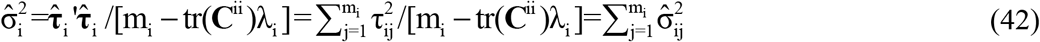

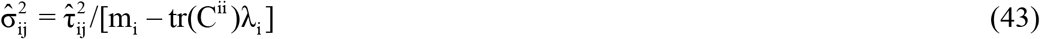

where **C**^ii^ is the submatrix in the inverse or generalized inverse of the coefficient matrix of the MME corresponding to the effect type, m_i_ = number of effects of the i^th^ effect type, and 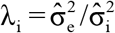. Dividing Equation 43 by 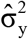 and multiplying by 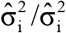 yield Equation 41, i.e.,

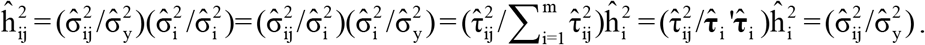

It is readily seen that the sum of all heritability estimates of the i^th^ effect type is the genomic heritability of the i^th^ effect type, i.e., 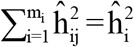. Note that Equations 42 and 43 using MME are only for proving Equation 41. The MME method is computationally prohibitive for estimating genetic effects and their variances under the multifactorial model although the MME method yield identical results as the CE method, which is computationally feasible for genomic estimation and prediction under the multifactorial model.

### Comparison between exact and approximate genomic epistasis relationship matrices

We evaluated the differences between AGERM and EGERM in genomic heritability estimates and prediction accuracies using a publicly available swine genomics data set that had 3534 animals from a single PIC nucleus pig line with five anonymous traits and 52,842 genotyped and imputed autosome SNPs after filtering by requiring minor allele frequency (MAF) > 0.001 and proportion of missing SNP genotypes < 0.100 [35]. The EGERM used the method of Jiang and Reif [13] and the AGERM methods were described in **Supplementary Text 1**. The heritability results showed that EGERM had slightly higher heritability estimates than AGERM except the A×A heritability of T3 where AGERM had slightly high estimate than EGERM (0.280 vs. 0.278, **Table 1**). From **Table 1**, effect type with nonzero heritability estimates was included in the prediction model for evaluating the observed prediction accuracy as the correlation between the GBLUP of genotypic values and the phenotypic values in each validation population and then averaged over all 10 validation populations. The results showed that AGERM and EGERM had the same prediction accuracy for this swine sample (**Table 2**). A disadvantage of EGERM is the computing time for the construction of EGERM, about 9.51 times as much time for pairwise relationship matrices, 8.29 as much time for third-order and 9.44 times as much time for fourth-order as required for AGERM (**Table 3**). However, computing time is not the deciding factor for choosing between the exact and approximate methods, because the multi-node approach that calculate each genomic relationship matrix in pieces and adds those pieces together can reduce the computing time to an acceptable level when multiple threads/cores are available and the two-step strategy can be used so that each genomic relationship is calculated only once for different traits and validation populations [31]. Prediction accuracy is the ultimate deciding factor for choosing between different methods. We reported results of comparing AGERM and EGERM using 60,671 SNPs and 22,022 first-lactation Holstein cows with phenotypic observations of daughter pregnancy rate, showing that AGERM and EGERM had the same heritability estimates and prediction accuracy, but EGERM required 21 times as much computing time as required by AGERM, which required 1.32 times as much time for the genomic additive relationship matrix [15]. The combined results of the swine and Holstein samples indicated that EGERM and AGERM had similar results and that the computing difficulty of EGERM over AGERM increased rapidly as the sample size increased. Given the computing difficulty of EGERM and the negligible differences between EGERM and AGERM in prediction accuracy, AGERM should be favored for its mathematical simplicity and computing efficiency at least for samples with 50,000 SNPs or more.

**Table 1.** Genomic heritability estimates of additive, dominance and epistasis effects up to the third-order for five traits in a swine population.

|  | Trait |  |  |  |  |
| --- | --- | --- | --- | --- | --- |
|  | T1 | T2 | T3 | T4 | T5 |
| Effect | Exact genomic epistasis relationship matrices (EGERM) |  |  |  |  |
| A | 0.023 | 0.217 | 0.131 | 0.336 | 0.366 |
| D | 0.000 | 0.013 | 0.000 | 0.000 | 0.052 |
| A×A | 0.046 | 0.186 | 0.278 | 0.017 | 0.054 |
| A×D | 0.000 | 0.000 | 0.091 | 0.000 | 0.000 |
| D×D | 0.000 | 0.000 | 0.091 | 0.000 | 0.000 |
| A×A×A | 0.000 | 0.000 | 0.000 | 0.000 | 0.000 |
| A×A×D | 0.000 | 0.000 | 0.079 | 0.000 | 0.000 |
| A×D×D | 0.000 | 0.000 | 0.102 | 0.000 | 0.000 |
| D×D×D | 0.000 | 0.000 | 0.117 | 0.000 | 0.000 |
| Total heritability | 0.069 | 0.416 | 0.889 | 0.354 | 0.471 |
| Effect | Approximate genomic epistasis relationship matrices (AGERM) |  |  |  |  |
| A | 0.022 | 0.215 | 0.139 | 0.329 | 0.360 |
| D | 0.000 | 0.013 | 0.000 | 0.000 | 0.051 |
| A×A | 0.043 | 0.176 | 0.280 | 0.016 | 0.050 |
| A×D | 0.000 | 0.000 | 0.091 | 0.000 | 0.000 |
| D×D | 0.000 | 0.000 | 0.090 | 0.000 | 0.000 |
| A×A×A | 0.000 | 0.000 | 0.000 | 0.000 | 0.000 |
| A×A×D | 0.000 | 0.000 | 0.075 | 0.000 | 0.000 |
| A×D×D | 0.000 | 0.000 | 0.095 | 0.000 | 0.000 |
| D×D×D | 0.000 | 0.000 | 0.109 | 0.000 | 0.000 |
| Total heritability | 0.065 | 0.404 | 0.879 | 0.346 | 0.461 |

**Table 2.** Observed prediction accuracy of epistasis models relative to the additive model for five traits in a swine population.

|  | Trait |  |  |  |  |
| --- | --- | --- | --- | --- | --- |
|  | T1 | T2 | T3 | T4 | T5 |
| Prediction accuracy of SNP model |  |  |  |  |  |
| A | 0.066 | 0.495 | 0.326 | 0.468 | 0.493 |
| A+D | 0.056 | 0.495 | 0.326 | 0.468 | 0.496 |
| Epistasis model | A+AA | A+D+AA | A+AA+AD+DD+<br>AAD+ADD+DDD | A+AA | A+D+AA |
| EGERM |  |  |  |  |  |
| Prediction accuracy | 0.063 | 0.498 | 0.336 | 0.468 | 0.497 |
| Accuracy increase (%) | -4.545 | 0.606 | 3.067 | 0.000 | 0.202 |
| AGERM |  |  |  |  |  |
| Prediction accuracy | 0.063 | 0.498 | 0.336 | 0.468 | 0.497 |
| Accuracy increase (%) | -4.545 | 0.606 | 3.067 | 0.000 | 0.202 |

**Table 3.**
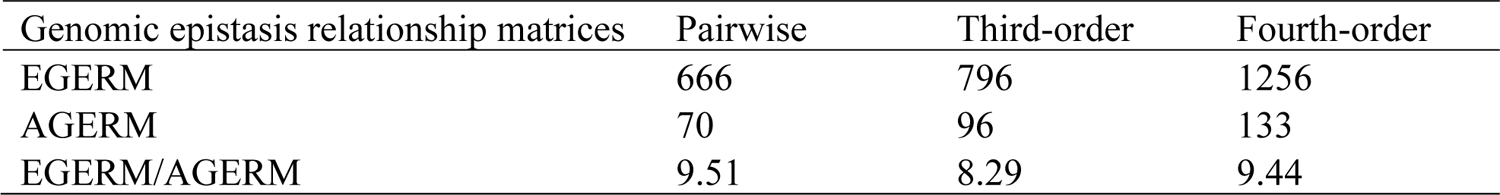
Computing time (in seconds) for the construction of exact and approximate genomic epistasis relationship matrices for a swine population with 3534 pigs and 52,843 SNPs using 20 threads of the Mangi supercomputer of the Minnesota Supercomputer Institute at the University of Minnesota.

| Genomic epistasis relationship matrices | Pairwise | Third-order | Fourth-order |
| --- | --- | --- | --- |
| EGERM | 666 | 796 | 1256 |
| AGERM | 70 | 96 | 133 |
| EGERM/AGERM | 9.51 | 8.29 | 9.44 |

### Numerical demonstration

The methods of genomic epistasis relationship matrices based on the additive and dominance model matrices, GREML, GBLUP and reliability, and estimation of effect heritability are demonstrated using a R program (DEMO.R) and a small artificial sample for the convenience of reading the numerical results (**Supplementary Text S2 and R program**). Because of the artificial nature and the extremely small sample size, this numerical demonstration does not have any genetic and methodology implications and is for showing calculations of the methods only. This R program is an extension of the R demo program of GVCHAP that integrates SNP and haplotype effects and has a computing pipeline for producing the input haplotype data from the SNP data [31].

## CONCLUSION

The multifactorial methods with SNP, haplotype and epistasis effects up to the third-order provide an approach to investigate the contributions of global low-order and local high-order epistasis effects to the phenotypic variance and the accuracy of gnomic prediction. Genomic heritability of each effect type from GREML and prediction accuracy from validation studies using GBLUP can be used jointly to identify effect types contributing to the phenotypic variance and the accuracy of genomic prediction, and the GBLUP for the multifactorial model with selected effect type can be used for genomic evaluation. With many capabilities including the use of intra- and inter-chromosome separately, the multifactorial methods offer a significant methodology capability to investigate and utilize complex genetic mechanisms for genomic prediction and for understanding the complex genome-phenome relationships.

## Supporting information

Text S1, Text S2, R program, Table S1

## AUTHOR CONTRIBUTIONS

YD conceived this study and derived the formulations. ZL contributed to formulations of the epistasis genomic relationships, implemented the epistasis methods in EPIHAP, validated and evaluated the methods. DP contributed to the data processing for methodology evaluation. YD and ZL prepared the manuscript.

## FUNDING

This research was supported by the National Institutes of Health’s National Human Genome Research Institute, grant R01HG012425 as part of the NSF/NIH Enabling Discovery through GEnomics (EDGE) Program; grant 2020-67015-31133 from the USDA National Institute of Food and Agriculture; and project MIN-16-124 of the Agricultural Experiment Station at the University of Minnesota. The funders had no role in study design, data collection and analysis, decision to publish, or preparation of the manuscript.

## CONFLICT OF INTEREST STATEMENT

The authors declare that the research was conducted in the absence of any commercial or financial relationships that could be construed as a potential conflict of interest.

## ACKNOWLEDGMENTS

The supercomputer of Minnesota Supercomputer Institute at the University of Minnesota and the Ceres and Atlas high performance computing system of USDA-ARS were used for the evaluation and testing of the methods and EPIHAP computing package. The use of the Ceres and Atlas computers by this research was supported by USDA-ARS projects 8042-31000-002-00-D and 8042-31000-001-00-D. The authors thank Paul VanRaden, Steven Schroeder and Ransom Baldwin for help with the use of the USDA-ARS computing facilities.

## SUPPLEMENTARY MATERIAL

**Text S1**. Quantitative Genetics Models and Genomic Epistasis Relationship Matrices

**Text S2**. Numerical Demonstration

**R program**. DEMO.R for Numerical Demonstration

**TABLE S1** | Notations of the quantitative genetics (QG) model, reparameterized and equivalent QG (RE-QG) model, and multifactorial (MF) model.

## Notes

### Competing Interest Statement

The authors have declared no competing interest.

### Summary of Updates

Line 323, (36) is changed to 7.

