## Supplementary material for "Multifactorial Methods Integrating Haplotype and Epistasis Effects for Genomic Estimation and Prediction of Quantitative Traits": Text S1, Text S2, R program, Table S1

Yang Da, Zuoxiang Liang, Dzianis Prakapenka
Department of Animal Science, University of Minnesota, Saint Paul, MN 55108, USA

#### Text S1. Quantitative Genetics Models and Genomic Epistasis Relationship Matrices

**Quantitative genetics (QG) model with additive and dominance effects and values**
The mixed model with single-SNP additive and dominance effects, haplotype additive effects and pairwise SNP epistasis effects described below is based on the quantitative genetics (QG) model resulting from the genetic partition of single-SNP genotypic values [1, 2], haplotype genotypic values [3], and pairwise genotypic values [4]. An advantage of this QG model is the readily available quantitative genetics interpretations of SNP additive and dominance effects, values and variances; haplotype additive effects, values and variances; epistasis effects, values and variances; and the corresponding SNP, haplotype and epistasis heritability estimates.

This section describes the QG-model of SNP additive and dominance effects of two loci that will be used for defining the epitasis models. Assuming two SNPs with alleles *A* and *a* at SNP 1 and *B* and *b* at SNP 2, the two-locus model with SNP additive and dominance effects for the nine two-locus genotypic values can be expressed as:

$$\mathbf{g} = \mathbf{1}\mu + \mathbf{a}_1 + \mathbf{a}_2 + \mathbf{d}_1 + \mathbf{d}_2 = \mathbf{1}\mu + \mathbf{w}_{\alpha 1}\alpha_{1o} + \mathbf{w}_{\alpha 2}\alpha_{2o} + \mathbf{w}_{\delta 1}\delta_{1o} + \mathbf{w}_{\delta 2}\delta_{2o} \quad \text{(S1.1)}$$

$$= \begin{bmatrix} g_{AABB} \\ g_{AABb} \\ g_{AAbb} \\ g_{AaBB} \\ g_{AaBb} \\ g_{Aabb} \\ g_{aaBB} \\ g_{aaBb} \\ g_{aabb} \end{bmatrix} = \begin{bmatrix} 1 \\ 1 \\ 1 \\ 1 \\ 1 \\ 1 \\ 1 \\ 1 \\ 1 \end{bmatrix} \mu + \begin{bmatrix} w_{\alpha 1}^{11} \\ w_{\alpha 1}^{11} \\ w_{\alpha 1}^{11} \\ w_{\alpha 1}^{12} \\ w_{\alpha 1}^{12} \\ w_{\alpha 1}^{12} \\ w_{\alpha 1}^{22} \\ w_{\alpha 1}^{22} \\ w_{\alpha 1}^{22} \end{bmatrix} \alpha_{1o} + \begin{bmatrix} w_{\alpha 2}^{11} \\ w_{\alpha 2}^{12} \\ w_{\alpha 2}^{22} \\ w_{\alpha 2}^{11} \\ w_{\alpha 2}^{12} \\ w_{\alpha 2}^{22} \\ w_{\alpha 2}^{11} \\ w_{\alpha 2}^{12} \\ w_{\alpha 2}^{22} \end{bmatrix} \alpha_{2o} + \begin{bmatrix} w_{\delta 1}^{11} \\ w_{\delta 1}^{11} \\ w_{\delta 1}^{11} \\ w_{\delta 1}^{12} \\ w_{\delta 1}^{12} \\ w_{\delta 1}^{12} \\ w_{\delta 1}^{22} \\ w_{\delta 1}^{22} \\ w_{\delta 1}^{22} \end{bmatrix} \delta_{1o} + \begin{bmatrix} w_{\delta 2}^{11} \\ w_{\delta 2}^{12} \\ w_{\delta 2}^{22} \\ w_{\delta 2}^{11} \\ w_{\delta 2}^{12} \\ w_{\delta 2}^{22} \\ w_{\delta 2}^{11} \\ w_{\delta 2}^{12} \\ w_{\delta 2}^{22} \end{bmatrix} \delta_{2o}$$

where $\mathbf{g}$ = $9\times 1$ column vector of genotypic values; $\mu$ = common mean; $\alpha_{io}$ = additive effect of SNP i ($i = 1, 2$); $\mathbf{w}_{\alpha i}$ = $9\times 1$ model matrix of $\alpha_{io}$; $\delta_{io}$ = dominance effect of SNP i ($i = 1, 2$); $\mathbf{w}_{\delta 1}$ = $9\times 1$ model matrix of $\delta_{io}$, $\mathbf{a}_i = \mathbf{w}_{\alpha i}\alpha_{io}$ = $9\times 1$ column vector of additive values of SNP i ($i = 1, 2$); and $\mathbf{d}_i = \mathbf{w}_{\delta 1}\delta_{1o}$ = $9\times 1$ column vector of dominance values of SNP i ($i = 1, 2$). The additive codings of the genotypes of the i[th] SNP in $\mathbf{w}_{\alpha i}$ are $w_{\alpha i}^{11} = 2q_i$, $w_{\alpha i}^{12} = q_i - p_i$, $w_{\alpha i}^{22} = -2p_i$, and the dominance codings in $\mathbf{w}_{\delta 1}$ are $w_{\delta i}^{11} = -2q_i^2$, $w_{\delta i}^{12} = 2p_i q_i$, and $w_{\delta i}^{22} = -2p_i^2$ for $A_1A_1$, $A_1A_2$ and $A_2A_2$ genotypes of the i[th] SNP respectively, with $p_i$ = allele frequency of $A_1$ and $q_i$ = allele frequency of $A_2$ [1]. Equation (S1.1) is the foundation for pairwise epistasis effects as the interaction between $\alpha_{1o}$ and $\alpha_{2o}$, $\alpha_{1o}$ and $\delta_{2o}$, $\delta_{1o}$ and $\alpha_{1o}$, and $\delta_{1o}$ and $\delta_{2o}$ under the assumption of linkage equilibrium (LE) that allows simplified genomic epistasis relationship matrices based on SNP additive and dominance relationship matrices without creating the epistasis model matrices. Subscript 'o' denotes a genetic effect form the original quantitative genetics model of Equation (S1.1) and later will be removed in the raparameterized and equivalent model resulting from the use of genomic relationship matrices.

**Epistasis effects, model matrices and values**

The epistasis effects defined by Cockerham method can cover high-order epistasis effects. However, the biological significance of high-order epistasis is unknown. This study only considers second- and third-order epistasis. Based on the single-SNP model of two loci defined by Equation (S1.1) and the epistasis effects defined by Cockerham [4], the model matrices of pairwise epistasis effects for n individuals are expressed as the Hadamard products between the model matrices of the additive and dominance effects:

$$\mathbf{E}_2 = \mathbf{w}_{\alpha 1} \# \mathbf{w}_{\alpha 2} (\alpha\alpha)_o + \mathbf{w}_{\alpha 1} \# \mathbf{w}_{\delta 2} (\alpha\delta)_o + \mathbf{w}_{\alpha 2} \# \mathbf{w}_{\delta 1} (\delta\alpha)_o + \mathbf{w}_{\delta 1} \# \mathbf{w}_{\delta 2} (\delta\delta)_o \quad \text{(S1.2)}$$

where $(\alpha\alpha)_o$ = additive × additive (A×A) epistasis effect = allele × allele interaction effect, $(\alpha\delta)_o$ = additive × dominance (A×D) epistasis effects = allele × genotype interaction effect, $(\delta\alpha)_o$ dominance × additive (D×A) epistasis effect = genotype × allele interaction effect, $(\delta\delta)_o$ = dominance × dominance (D×D) epistasis effect, $\mathbf{w}_{\alpha 1} \# \mathbf{w}_{\alpha 2}$ = coefficients of $(\alpha\alpha)_o$, $\mathbf{w}_{\alpha 1} \# \mathbf{w}_{\delta 2}$ = coefficients of $(\alpha\delta)_o$, $\mathbf{w}_{\alpha 2} \# \mathbf{w}_{\delta 1}$ = coefficients of $(\delta\alpha)_o$, $\mathbf{w}_{\delta 1} \# \mathbf{w}_{\delta 2}$ = coefficients of $(\delta\delta)_o$, and '#' indicates the Hadamard product. For m SNPs and n individuals, the total epistasis values of the genome as the summation of all pairwise epistasis values defined by Equation (S1.2) are:

$$\begin{aligned}\mathbf{E}_2 &= \textstyle\sum_{i=1}^{m-1}\sum_{j=i+1}^{m}(\mathbf{w}_{\alpha 1}^{i} \# \mathbf{w}_{\alpha 2}^{j})(\alpha\alpha)_o^{ij} + \sum_{i=1}^{m-1}\sum_{j=i+1}^{m}(\mathbf{w}_{\alpha 1}^{i} \# \mathbf{w}_{\delta 2}^{j})(\alpha\delta)_o^{ij} \\ &\quad + \textstyle\sum_{i=1}^{m-1}\sum_{j=i+1}^{m}(\mathbf{w}_{\delta 1}^{i} \# \mathbf{w}_{\alpha 2}^{j})(\delta\alpha)_o^{ij} + \sum_{i=1}^{m-1}\sum_{j=i+1}^{m}(\mathbf{w}_{\delta 1}^{i} \# \mathbf{w}_{\delta 2}^{j})(\delta\delta)_o^{ij} \\ &= \mathbf{W}_{\alpha\alpha}(\boldsymbol{\alpha\alpha})_o + [\mathbf{W}_{\alpha\delta}(\boldsymbol{\alpha\delta})_o + \mathbf{W}_{\delta\alpha}(\boldsymbol{\delta\alpha})_o] + \mathbf{W}_{\delta\delta}(\boldsymbol{\delta\delta})_o \qquad \text{(S1.3)} \\ &= \mathbf{W}_{\alpha\alpha}(\boldsymbol{\alpha\alpha})_o + \mathbf{W}_{\alpha\delta}^{(2)}(\boldsymbol{\alpha\delta})_o^{(2)} + \mathbf{W}_{\delta\delta}(\boldsymbol{\delta\delta})_o \\ &= \mathbf{aa} + \mathbf{ad} + \mathbf{da} + \mathbf{dd} = \mathbf{aa} + (\mathbf{ad})^{(2)} + \mathbf{dd}\end{aligned}$$

where $(\boldsymbol{\alpha\alpha})_o$ = $\binom{m}{2}\times 1$ column vector of A×A epistasis effects (with $\binom{m}{2} = m(m-1)/2$ = allele × allele interaction effects, $\mathbf{W}_{\alpha\alpha}$ = $n\times\binom{m}{2}$ model matrix of $(\boldsymbol{\alpha\alpha})_o$, $(\boldsymbol{\alpha\delta})_o$ = $\binom{m}{2}\times 1$ column vector of A×D epistasis effects = allele × genotype interaction effects, $\mathbf{W}_{\alpha\delta}$ = $n\times\binom{m}{2}$ model matrix of $(\boldsymbol{\alpha\delta})_o$, $(\boldsymbol{\delta\alpha})_o$ = $\binom{m}{2}\times 1$ column vector of D×A epistasis effects = genotype × allele interaction effects, $\mathbf{W}_{\delta\alpha}$ = $n\times\binom{m}{2}$ model matrix of $(\boldsymbol{\delta\alpha})_o$, $(\boldsymbol{\delta\delta})_o$ = $\binom{m}{2}\times 1$ column vector of D×D epistasis effects = genotype × genotype interaction effects, $\mathbf{W}_{\delta\delta}$ = $n\times\binom{m}{2}$ model matrix of $(\boldsymbol{\delta\delta})_o$, $\mathbf{aa} = \mathbf{W}_{\alpha\alpha}(\boldsymbol{\alpha\alpha})_o$ = $n\times 1$ column vector of genomic A×A epistasis values, $\mathbf{ad} = \mathbf{W}_{\alpha\delta}(\boldsymbol{\alpha\delta})_o$ = $n\times 1$ column vector of genomic A×D epistasis values, $\mathbf{da} = \mathbf{W}_{\delta\alpha}(\boldsymbol{\delta\alpha})_o$ = $n \times 1$ column vector of genomic D×A epistasis values, $\mathbf{dd} = \mathbf{W}_{\delta\delta}(\boldsymbol{\delta\delta})_o$ = $n \times 1$ column vector of genomic D×D epistasis values, $(\mathbf{ad})^{(2)} = n \times 1$ column vector of genomic A×D and D×A epistasis values, $(\boldsymbol{\alpha\delta})_o^{(2)} = [(\boldsymbol{\alpha\delta})_o', (\boldsymbol{\delta\alpha})_o']' = 2\binom{m}{2} \times 1$ column vector of A×D and D×A epistasis effects, $\mathbf{W}_{\alpha\delta}^{(2)} = (\mathbf{W}_{\alpha\delta}, \mathbf{W}_{\delta\alpha}) = n \times [2\binom{m}{2}]$ model matrix of $(\boldsymbol{\alpha\delta})_o^{(2)}$, and superscript '(2)' indicates the vector or matrix involves two alternative expressions.

**Third-order epistasis effects, model matrices and values**

The third-order epistasis values and effects under the QG model are:

$$
\begin{aligned}
\mathbf{E}_3 &= \sum_{i=1}^{m-2}\sum_{j=i+1}^{m-1}\sum_{k=j+1}^{m}(\mathbf{w}_{\alpha1}^{i} \# \mathbf{w}_{\alpha2}^{j} \# \mathbf{w}_{\alpha3}^{j})(\alpha\alpha\alpha)_o^{ijk} + \sum_{i=1}^{m-2}\sum_{j=i+1}^{m-1}\sum_{k=j+1}^{m}(\mathbf{w}_{\alpha1}^{i} \# \mathbf{w}_{\alpha2}^{j} \# \mathbf{w}_{\delta3}^{j})(\alpha\alpha\delta)_o^{ijk} \\
&+ \sum_{i=1}^{m-2}\sum_{j=i+1}^{m-1}\sum_{k=j+1}^{m}(\mathbf{w}_{\alpha1}^{i} \# \mathbf{w}_{\delta2}^{j} \# \mathbf{w}_{\alpha3}^{j})(\alpha\delta\alpha)_o^{ijk} + \sum_{i=1}^{m-2}\sum_{j=i+1}^{m-1}\sum_{k=j+1}^{m}(\mathbf{w}_{\delta1}^{i} \# \mathbf{w}_{\alpha2}^{j} \# \mathbf{w}_{\alpha3}^{j})(\delta\alpha\alpha)_o^{ijk} \\
&+ \sum_{i=1}^{m-2}\sum_{j=i+1}^{m-1}\sum_{k=j+1}^{m}(\mathbf{w}_{\alpha1}^{i} \# \mathbf{w}_{\delta2}^{j} \# \mathbf{w}_{\delta3}^{j})(\alpha\delta\delta)_o^{ijk} + \sum_{i=1}^{m-2}\sum_{j=i+1}^{m-1}\sum_{k=j+1}^{m}(\mathbf{w}_{\delta1}^{i} \# \mathbf{w}_{\delta2}^{j} \# \mathbf{w}_{\alpha3}^{j})(\delta\delta\alpha)_o^{ijk} \\
&+ \sum_{i=1}^{m-2}\sum_{j=i+1}^{m-1}\sum_{k=j+1}^{m}(\mathbf{w}_{\delta1}^{i} \# \mathbf{w}_{\alpha2}^{j} \# \mathbf{w}_{\delta3}^{j})(\delta\alpha\delta)_o^{ijk} + \sum_{i=1}^{m-2}\sum_{j=i+1}^{m-1}\sum_{k=j+1}^{m}(\mathbf{w}_{\delta1}^{i} \# \mathbf{w}_{\delta2}^{j} \# \mathbf{w}_{\delta3}^{j})(\delta\delta\delta)_o^{ijk} \\
&= \mathbf{W}_{\alpha\alpha\alpha}(\boldsymbol{\alpha\alpha\alpha})_o + [\mathbf{W}_{\alpha\alpha\delta}(\boldsymbol{\alpha\alpha\delta})_o + \mathbf{W}_{\alpha\delta\alpha}(\boldsymbol{\alpha\delta\alpha})_o + \mathbf{W}_{\delta\alpha\alpha}(\boldsymbol{\delta\alpha\alpha})_o] \qquad \text{(S1.4)} \\
&+ [\mathbf{W}_{\alpha\delta\delta}(\boldsymbol{\alpha\delta\delta})_o + \mathbf{W}_{\delta\alpha\delta}(\boldsymbol{\delta\alpha\delta})_o + \mathbf{W}_{\delta\delta\alpha}(\boldsymbol{\delta\delta\alpha})_o] + \mathbf{W}_{\delta\delta\delta}(\boldsymbol{\delta\delta\delta})_o \\
&= \mathbf{W}_{\alpha\alpha\alpha}(\boldsymbol{\alpha\alpha\alpha})_o + \mathbf{W}_{\alpha\alpha\delta}^{(3)}(\boldsymbol{\alpha\alpha\delta})_o^{(3)} + \mathbf{W}_{\alpha\delta\delta}^{(3)}(\boldsymbol{\alpha\delta\delta})_o^{(3)} + \mathbf{W}_{\delta\delta\delta}(\boldsymbol{\delta\delta\delta})_o \\
&= \mathbf{aaa} + (\mathbf{aad} + \mathbf{ada} + \mathbf{daa}) + (\mathbf{add} + \mathbf{dad} + \mathbf{dda}) + \mathbf{ddd} \\
&= \mathbf{aaa} + \mathbf{aad}^{(3)} + \mathbf{add}^{(3)} + \mathbf{ddd}
\end{aligned}
$$

where

$(\boldsymbol{\alpha\alpha\alpha})_o = \binom{m}{3} \times 1$ column vector of A×A×A effects with $\binom{m}{3} = m(m-1)(m-2)/6$,

$(\boldsymbol{\alpha\alpha\delta})_o = \binom{m}{3} \times 1$ column vector of A×A×D effects,

$(\boldsymbol{\alpha\delta\alpha})_o = \binom{m}{3} \times 1$ column vector of A×D×A effects,

$(\boldsymbol{\delta\alpha\alpha})_o = \binom{m}{3} \times 1$ column vector of D×A×A effects,

$(\boldsymbol{\alpha\delta\delta})_o = \binom{m}{3} \times 1$ column vector of A×D×D effects,

$(\boldsymbol{\delta\alpha\delta})_o = \binom{m}{3} \times 1$ column vector of D×A×D effects,

$(\boldsymbol{\delta\delta\alpha})_o = \binom{m}{3} \times 1$ column vector of D×D×A effects,

$(\boldsymbol{\delta\delta\delta})_o = \binom{m}{3} \times 1$ column vector of D×D×A effects,

$(\boldsymbol{\alpha\alpha\delta})_o^{(3)} = [(\boldsymbol{\alpha\alpha\delta})_o', (\boldsymbol{\alpha\delta\alpha})_o', (\boldsymbol{\delta\alpha\alpha})_o']' = 3\binom{m}{3} \times 1$ column vector of A×A×D, A×D×A and D×A×A effects,

$(\boldsymbol{\alpha\delta\delta})_o^{(3)} = [\,[(\boldsymbol{\alpha\alpha\delta})_o', (\boldsymbol{\alpha\delta\alpha})_o', (\boldsymbol{\delta\alpha\alpha})_o']' = 3\binom{m}{3} \times 1$ column vector of A×D×D, D×A×D and D×D×A effects,

$\mathbf{W}_{\alpha\alpha\alpha} = n \times \binom{m}{3}$ model matrix of $(\boldsymbol{\alpha\alpha\alpha})_o$,

$\mathbf{W}_{\alpha\alpha\delta}^{(3)} = (\mathbf{W}_{\alpha\alpha\delta}, \mathbf{W}_{\alpha\delta\alpha}, \mathbf{W}_{\delta\alpha\alpha}) = n \times [3\binom{m}{3}]$ model matrix of $(\boldsymbol{\alpha\alpha\delta})_o^{(3)}$,

$\mathbf{W}_{\alpha\delta\delta}^{(3)} = (\mathbf{W}_{\alpha\delta\delta}, \mathbf{W}_{\delta\alpha\delta}, \mathbf{W}_{\delta\delta\alpha}) = n \times [3\binom{m}{3}]$ model matrix of $(\boldsymbol{\alpha\delta\delta})_o^{(3)}$,

$\mathbf{W}_{\delta\delta\delta} = n \times \binom{m}{3}$ model matrix of $(\boldsymbol{\delta\delta\alpha})_o$,

$\mathbf{aaa} = \mathbf{W}_{\alpha\alpha\alpha}(\boldsymbol{\alpha\alpha\alpha})_o = n \times 1$ column vector of A×A×A epistasis values,

$\mathbf{aad} = n \times 1$ column vector of A×A×D epistasis values,

$\mathbf{ada} = n \times 1$ column vector of A×D×A epistasis values,

$\mathbf{daa} = n \times 1$ column vector of D×A×A epistasis values,

$\mathbf{add} = n \times 1$ column vector of A×D×D epistasis values,

$\mathbf{dad} = n \times 1$ column vector of D×A×D epistasis values,

$\mathbf{dda} = n \times 1$ column vector of D×D×A epistasis values,

$\mathbf{aad}^{(3)} = \mathbf{aad} + \mathbf{ada} + \mathbf{daa} = \mathbf{W}_{\alpha\alpha\delta}^{(3)}(\boldsymbol{\alpha\alpha\delta})_o^{(3)} = n \times 1$ column vector of the sum of A×A×D, A×D×A and D×A×A epistasis values,

$\mathbf{add}^{(3)} = \mathbf{add} + \mathbf{dad} + \mathbf{dda} = \mathbf{W}_{\alpha\delta\delta}^{(3)}(\boldsymbol{\alpha\delta\delta})_o^{(3)} = n \times 1$ column vector of the sum of A×D×D, D×A×D and D×D×A epistasis values,

$\mathbf{ddd} = \mathbf{W}_{\delta\delta\delta}(\boldsymbol{\delta\delta\delta})_o = n \times 1$ column vector of D×D×D epistasis values.

An important point of Equations (S1.3) and (S1.4) is that the epistasis model matrices are expressed as the Hadamard products between SNP additive and dominance model matrices. This result is the foundation for calculating genomic epistasis relationship matrices using SNP additive and dominance model matrices without creating the epistasis model matrices.

**Quantitative genetics model with haplotype additive effects and values**

The haplotype additive effects and values are derived by the multiallelic genetic partition of haplotype genotypic values and variances, and the relationship between the haplotype additive effects and values is:

$$\mathbf{a}_h = \mathbf{W}_{\alpha h}\boldsymbol{\alpha}_{oh} \quad \text{(S1.5)}$$

where $\mathbf{a}_h = n \times 1$ haplotype genomic additive values, $\boldsymbol{\alpha}_h = n_{\alpha h} \times 1$ column vector of haplotype additive effects ($n_{\alpha h}$ = number of haplotype additive effects), and $\mathbf{W}_{\alpha h} = n \times n_{\alpha h}$ model matrix of $\boldsymbol{\alpha}_{oh}$. The haplotype coding represented by $w_{\alpha h}^{ij,k}$ in $\mathbf{W}_{\alpha h}$ is: $w_{\alpha h}^{ij,k} = 2p_k$ for $i, j \neq k$ ($a_{ij}$ and $\alpha_{o1k}$ do not share allele k), $w_{\alpha h}^{ij,k} = -(1-2p_k)$ for $i \neq j$ but $i \neq k$ or $j \neq k$ ($a_{ij}$ and $\alpha_{o1k}$ share allele k, $i \neq j$), and $w_{\alpha h}^{ij,k} = -2(1-p_k)$ for $i = j = k$, where $a_{ij}$ = additive value of haplotype genotype with the $i^{th}$ and $j^{th}$ haplotypes, and $\alpha_{o1k}$ = additive effect or the average effect of gene substitution as the difference between the allelic (haplotype) effects of the first and the $k^{th}$ haplotypes [3].

**Integrated QG model and multifactorial notations (Model-I)**

This integrated model is advantageous over the separate SNP, haplotype, and epistasis models, because the predicted effects or values of each effect type is based on the phenotypic values after removing the predicted effects or values of the remaining effect types, whereas such predicted effects or values are unavailable from the separate models, as shown in the main text.

Combining the additive and dominance values [1], haplotype additive values (Equation S1.5), and the epistasis values (Equations S1.3 and S1.4), The integrated QG model with SNP additive and dominance, haplotype additive and epistasis effects and values is:

$$\begin{aligned}\mathbf{g} &= \mathbf{1}\mu + \mathbf{a} + \mathbf{d} + \mathbf{a}_h + \mathbf{aa} + \mathbf{ad}^{(2)} + \mathbf{dd} + \mathbf{aaa} + \mathbf{aad}^{(3)} + \mathbf{add}^{(3)} + \mathbf{ddd} \\ &= \mathbf{1}\mu + \mathbf{W}_\alpha \boldsymbol{\alpha}_o + \mathbf{W}_\delta \boldsymbol{\delta}_o + \mathbf{W}_{\alpha h}\boldsymbol{\alpha}_{ho} + \mathbf{W}_{\alpha\alpha}(\boldsymbol{\alpha\alpha})_o + \mathbf{W}_{\alpha\delta}^{(2)}(\boldsymbol{\alpha\delta})_o^{(2)} + \mathbf{W}_{\delta\delta}(\boldsymbol{\delta\delta})_o \\ &\quad + \mathbf{W}_{\alpha\alpha\alpha}(\boldsymbol{\alpha\alpha\alpha})_o + \mathbf{W}_{\alpha\alpha\delta}^{(3)}(\boldsymbol{\alpha\alpha\delta})_o^{(3)} + \mathbf{W}_{\alpha\delta\delta}^{(3)}(\boldsymbol{\alpha\alpha\delta})_o^{(3)} + \mathbf{W}_{\delta\delta\delta}(\boldsymbol{\delta\delta\delta})_o\end{aligned} \tag{S1.6}$$

where $\boldsymbol{\alpha}_o$ = $m\times1$ column vector of additive effects, $\mathbf{W}_\alpha$ = $N\times m$ model matrix of $\boldsymbol{\alpha}_o$ with $2q_i$, $q_i - p_i$, $-2p_i$ values for the $i^{th}$ column; $\boldsymbol{\delta}_o$ = $m\times1$ column vector of dominance effects, $\mathbf{W}_\delta$ = $N\times m$ model matrix of $\boldsymbol{\delta}_o$ with $-2q_i^2$, $2p_iq_i$, and $-2p_i^2$ values for the $i^{th}$ column, $\mathbf{a}$ = $n\times1$ column vector of additive values, $\mathbf{d}$ = $n\times1$ column vector of dominance values.

With the precise definitions of genetic effects, values and model matrices by Equations S1.3-S1.6, the integrated QG model of Equation S1.6 can be expressed succinctly using multifactorial notations as:

$$\mathbf{g} = \mu\mathbf{I} + \sum_{i=1}^{10}\mathbf{W}_i\boldsymbol{\tau}_{io} = \mu\mathbf{I} + \sum_{i=1}^{10}\mathbf{u}_i \tag{S1.7}$$

where $\boldsymbol{\tau}_{io}$ = genetic effects of the $i^{th}$ effect type from the original QG model, $\mathbf{W}_i$ = model matrix of $\boldsymbol{\tau}_{io}$, $\mathbf{u}_i = \mathbf{W}_i\boldsymbol{\tau}_{io}$ = genetic values of the $i^{th}$ effect type.

**Intra- and inter-chromosome pairwise epistasis effects, model matrices and values**

An epistasis GWAS in Holstein cattle showed that intra- and inter-chromosome epistasis effects affected different traits differently, e.g., daughter pregnancy rate was mostly affected by inter-chromosome epistasis effects whereas milk production traits were mostly affected by intra-chromosome epistasis effects [5]. Genomic heritability estimates of intra- and inter-chromosome heritabilities using methods in this article showed that intra-chromosome A×A heritability was 0.031, and inter-chromosome A×A heritability 0.178 for daughter pregnancy rate [6], consistent with the GWAS results of 21% intra-chromosome and 79% inter-chromosome A×A effects among the top 33,552 pairs of A×A effects. Therefore, dividing pairwise epistasis effects into intra- and inter-chromosome epistasis effects for genomic prediction and estimation provides a tool to help understand epistasis effects.

Pairwise epistasis values of Equation (S1.3) are partitioned into intra- and inter-chromosome pairwise epistasis values:

$$\begin{aligned}\mathbf{E}_2 &= \mathbf{W}_{\alpha\alpha}(\boldsymbol{\alpha\alpha})_o + \mathbf{W}_{\alpha\delta}^{(2)}(\boldsymbol{\alpha\delta})_o^{(2)} + \mathbf{W}_{\delta\delta}(\boldsymbol{\delta\delta})_o = \mathbf{aa} + (\mathbf{ad})^{(2)} + \mathbf{dd} \\ &= [\mathbf{W}_{\alpha\alpha}^{intra}(\boldsymbol{\alpha\alpha})_o^{intra} + \mathbf{W}_{\alpha\alpha}^{inter}(\boldsymbol{\alpha\alpha})^{inter}] + [\mathbf{W}_{\alpha\delta}^{(2)intra}(\boldsymbol{\alpha\delta})_o^{(2)intra} + \mathbf{W}_{\alpha\delta}^{(2)inter}(\boldsymbol{\alpha\delta})_o^{(2)inter}] \\ &\quad + [\mathbf{W}_{\delta\delta}^{intra}(\boldsymbol{\delta\delta})_o^{intra} + \mathbf{W}_{\delta\delta}^{inter}(\boldsymbol{\delta\delta})_o^{inter}] \\ &= [(\mathbf{aa})^{intra} + (\mathbf{aa})^{inter}] + [\mathbf{ad}^{(2)intra} + \mathbf{ad}^{(2)inter}] + [(\mathbf{dd})^{intra} + (\mathbf{dd})^{inter}]\end{aligned} \tag{S1.8}$$

where superscript 'intra' represents intra-chromosome and 'inter' represents inter-chromosome, $(\mathbf{aa})^{\text{intra}} = \mathbf{W}_{\alpha\alpha}^{\text{intra}}(\boldsymbol{\alpha\alpha})_{\text{o}}^{\text{intra}}$ = $n\times1$ column vector of intra-chromosome A×A values, $(\mathbf{aa})^{\text{inter}} = \mathbf{W}_{\alpha\alpha}^{\text{inter}}(\boldsymbol{\alpha\alpha})_{\text{o}}^{\text{inter}}$ = $n\times1$ column vector of inter-chromosome A×A values, $\mathbf{ad}^{(2)\text{intra}} = \mathbf{W}_{\alpha\delta}^{(2)\text{intra}}(\boldsymbol{\alpha\delta})_{\text{o}}^{(2)\text{intra}}$ = $n\times1$ column vector of intra-chromosome A×D and D×A values, $\mathbf{ad}^{(2)\text{inter}} = \mathbf{W}_{\alpha\delta}^{(2)\text{inter}}(\boldsymbol{\alpha\delta})_{\text{o}}^{(2)\text{inter}}$ = $n\times1$ column vector of inter-chromosome A×D and D×A values, $(\mathbf{dd})^{\text{intra}} = \mathbf{W}_{\delta\delta}^{\text{intra}}(\boldsymbol{\delta\delta})_{\text{o}}^{\text{intra}}$ = $n\times1$ column vector of intra-chromosome D×D values, $(\mathbf{dd})^{\text{inter}} = \mathbf{W}_{\delta\delta}^{\text{inter}}(\boldsymbol{\delta\delta})_{\text{o}}^{\text{inter}}$ = $n\times1$ column vector of inter-chromosome D×D values, $(\boldsymbol{\alpha\alpha})_{\text{o}}^{\text{intra}} = c_1\times1$ column vector of intra-chromosome A×A effects with $c_1$ being the number of intra-chromosome SNP pairs, $(\boldsymbol{\alpha\alpha})_{\text{o}}^{\text{inter}} = c_2\times1$ column vector of inter-chromosome A×A effects with $c_2$ being the number of inter-chromosome SNP pairs, $(\boldsymbol{\alpha\delta})_{\text{o}}^{(2)\text{intra}} = 2c_1\times1$ column vector of intra-chromosome A×D and D×A effects, $(\boldsymbol{\alpha\delta})_{\text{o}}^{(2)\text{inter}} = 2c_2\times1$ column vector of inter-chromosome A×D and D×A effects, $(\boldsymbol{\delta\delta})_{\text{o}}^{\text{intra}} = c_1\times1$ column vector of intra-chromosome D×D effects, $(\boldsymbol{\delta\delta})_{\text{o}}^{\text{inter}} = c_2\times1$ column vector of inter-chromosome D×D effects, $\mathbf{W}_{\alpha\alpha}^{\text{intra}} = n\times c_1$ model matrix of $(\boldsymbol{\alpha\alpha})_{\text{o}}^{\text{intra}}$, $\mathbf{W}_{\alpha\alpha}^{\text{inter}} = n\times c_2$ model matrix of $(\boldsymbol{\alpha\alpha})_{\text{o}}^{\text{inter}}$, $\mathbf{W}_{\alpha\delta}^{(2)\text{intra}} = n\times 2c_1$ model matrix of $(\boldsymbol{\alpha\delta})_{\text{o}}^{(2)\text{intra}}$, $\mathbf{W}_{\alpha\delta}^{(2)\text{inter}} = n\times 2c_1$ model matrix of $(\boldsymbol{\alpha\delta})_{\text{o}}^{(2)\text{inter}}$, $\mathbf{W}_{\delta\delta}^{\text{intra}} = n\times c_1$ model matrix of $(\boldsymbol{\delta\delta})_{\text{o}}^{\text{intra}}$, $\mathbf{W}_{\delta\delta}^{\text{inter}} = n\times c_2$ model matrix of $(\boldsymbol{\delta\delta})_{\text{o}}^{\text{inter}}$.

**Integrated QG model with intra- and inter-chromosome epistasis effects and multifactorial notations (Model-II)**

Replacing the pairwise epistasis effects, values and model matrices in Equation S1.6 with the intra- and inter-chromosome epistasis effects, values and model matrices of Equation S1.8 the integrated QG model with intra- and inter-chromosome epistasis effects is:

$$\begin{aligned}\mathbf{g} &= \mu\mathbf{I} + \mathbf{a} + \mathbf{d} + \mathbf{a}_h + [(\mathbf{aa})^{\text{intra}} + (\mathbf{aa})^{\text{inter}}] + [\mathbf{ad}^{(2)\text{intra}} + \mathbf{ad}^{(2)\text{inter}}] \\ &\quad + [(\mathbf{dd})^{\text{intra}} + (\mathbf{dd})^{\text{inter}}] + \mathbf{aaa} + \mathbf{aad}^{(3)} + \mathbf{add}^{(3)} + \mathbf{ddd} \\ &= \mathbf{W}_\alpha\boldsymbol{\alpha}_o + \mathbf{W}_\delta\boldsymbol{\delta}_o + \mathbf{W}_{\alpha h}\boldsymbol{\alpha}_{ho} + [\mathbf{W}_{\alpha\alpha}^{\text{intra}}(\boldsymbol{\alpha\alpha})_o^{\text{intra}} + \mathbf{W}_{\alpha\alpha}^{\text{inter}}(\boldsymbol{\alpha\alpha})^{\text{inter}}] \\ &\quad + [\mathbf{W}_{\alpha\delta}^{(2)\text{intra}}(\boldsymbol{\alpha\delta})_o^{(2)\text{intra}} + \mathbf{W}_{\alpha\delta}^{(2)\text{inter}}(\boldsymbol{\alpha\delta})_o^{(2)\text{inter}}] + [\mathbf{W}_{\delta\delta}^{\text{intra}}(\boldsymbol{\delta\delta})_o^{\text{intra}} + \mathbf{W}_{\delta\delta}^{\text{inter}}(\boldsymbol{\delta\delta})_o^{\text{inter}}] \\ &\quad + \mathbf{W}_{\alpha\alpha\alpha}(\boldsymbol{\alpha\alpha\alpha})_o + \mathbf{W}_{\alpha\alpha\delta}^{(3)}(\boldsymbol{\alpha\alpha\delta})_o^{(3)} + \mathbf{W}_{\alpha\delta\delta}^{(3)}(\boldsymbol{\alpha\alpha\delta})_o^{(3)} + \mathbf{W}_{\delta\delta\delta}(\boldsymbol{\delta\delta\delta})_o\end{aligned} \tag{S1.9}$$

With the precise definitions of genetic effects, values and model matrices by Equations S1.6 and S1.8, the integrated QG model of Equation S1.9 can be expressed succinctly using multifactorial notations as:

$$\mathbf{g} = \mu\mathbf{I} + \sum_{i=1}^{13}\mathbf{W}_i\boldsymbol{\tau}_{io} = \mu\mathbf{I} + \sum_{i=1}^{13}\mathbf{u}_i \tag{S1.10}$$

where $\boldsymbol{\tau}_{io}$ = genetic effects of the $i^{th}$ effect type from the original QG model, $\mathbf{W}_i$ = model matrix of $\boldsymbol{\tau}_{io}$, $\mathbf{u}_i = \mathbf{W}_i\boldsymbol{\tau}_{io}$ = genetic values of the $i^{th}$ effect type.

**General multifactorial model that covers both Model-I and Model-II**

The multifactorial model of Equations S1.7 and S.19 can be summarized by one general multifactorial model as:

$$\mathbf{g} = \mu\mathbf{I} + \sum_{i=1}^{f}\mathbf{W}_i\boldsymbol{\tau}_{io} = \mu\mathbf{I} + \sum_{i=1}^{f}\mathbf{u}_i \quad (S1.11)$$

where $\boldsymbol{\tau}_{io}$ = genetic effects of the $i^{th}$ effect type from the original QG model, $\mathbf{W}_i$ = model matrix of $\boldsymbol{\tau}_{io}$, $\mathbf{u}_i = \mathbf{W}_i\boldsymbol{\tau}_{io}$ = genetic values of the $i^{th}$ effect type, and f = number effect types, with f =10 for Model-I of Equations S1.6 and S1.7, and f = 13 for Model-II of Equations S1.9 and S1.10.

**Genomic epistasis relationship matrices using model matrices**

Based on the general multifactorial model of Equation S.11 that covers both Model-I and Model-II, the genomic relationship matrices are:

$$\mathbf{S}_i = (\mathbf{W}_i\mathbf{W}_i') / k_i \quad (S1.12)$$

$$k_i = tr(\mathbf{W}_i\mathbf{W}_i')/n \quad (S1.13)$$

where $\mathbf{S}_i$ = genomic epistasis relationship matrix of the $i^{th}$ effect type, and $k_i$ = the average of the diagonal elements of $\mathbf{W}_i\mathbf{W}_i'$, originally proposed for genomic additive relationship matrix [7].

This article uses the $k_i$ value of Equation S1.13 because the heritability estimate of each effect type has the interpretation of being the average variance of the genetic values for the effect type over all individuals, as shown in the main text. The original genomic relationship matrix uses the coefficient of total additive variance of all SNPs as the $k_i$ value [8]. We referred to the VanRaden method as Definition-I and the Hayes-Goddard method Definition-II [1, 2]. Definition-I is the genomic version of the pedigree additive relationship because Definition-I maintains the properties of the pedigree relationships and would be conceptually appropriate as the 'genomic version' of pedigree relationships. However, Definition-II will be used in this study because Definition-II yields the interpretation that the variance component is the average of the genetic variances of all individuals for the $i^{th}$ type of genetic effects, although Definition-I and Definition-II have the same prediction accuracy, as shown in the main text.

Equations S1.12 and S1.13 use the original model matrices. For epistasis effects, the model matrices are difficult or impossible to compute. Two methods are available for calculating Equations S1.12 and S1.13 without creating the epistasis model matrices.

**Genomic epistasis relationship matrices without creating epistasis model matrices**

The computing difficulty due to creating epistasis model matrices can be removed by computing the approximate genomic epistasis relationship matrices (AGERM) as the genomic version of Henderson's Hadamard products between additive and dominance relationship matrices [9-11], or by the exact genomic epistasis relationship matrices (EGERM). AGERM contains intra-locus epistasis that should not exist [12] and EGERM removes intra-locus epistasis from AGERM based on products between SNP genomic additive and dominance relationship matrices [12, 13]. Although EGERM is theoretically more appealing than AGERM for being exact, the difference between these two methods in prediction accuracy and heritability estimates was nonexistent for a Holstein dataset with 78,964 SNPs [6] and was negligible for a swine dataset with 52,842 SNPs in this article, but EGERM required 21 times as much computing time as required by AGERM for the Holstein dataset, and required about 9 times as much computing time as required by AGERM for the swine dataset in this article. The lack of difference between these two methods and the computing inefficiency of EGERM should favor AGERM for its mathematical simplicity and computing efficiency at least for datasets with 50,000 or more SNPs. In this article, we use general notations of genomic epistasis relationship matrices defined by Equation S1.12 that can be either

AGERM or EGERM. The EPIHAP computing package [6, 14] implements both AGERM or EGERM. The use of AGERM or EGERM implies the assumption of linkage equilibrium (LE). Both AGERM and EGERM have general formulations to cover any order of epistasis effects, but no evidence is available in livestock populations showing the epistasis beyond the third-order have contributions to the phenotypic variance and prediction accuracy. Therefore, this article limits to second- and third-order epistasis effects.

**Exact genomic epistasis relationship matrices (EGERM)**
For EGERM, we use the general formula of Jiang and Reif [13]. For the example of pairwise epistasis effects in Equation S1.6, the products of the epistasis model matrices for defining pairwise EGERM are calculated as:

$$\mathbf{W}_{\alpha\alpha}\mathbf{W}_{\alpha\alpha}' = \frac{1}{2}[(\mathbf{W}_{\alpha}\mathbf{W}_{\alpha}')\#(\mathbf{W}_{\alpha}\mathbf{W}_{\alpha}') - (\mathbf{W}_{\alpha}\#\mathbf{W}_{\alpha})(\mathbf{W}_{\alpha}\#\mathbf{W}_{\alpha})'] \quad \text{(S1.14)}$$

$$\begin{aligned} \mathbf{W}_{\alpha\delta}^{(2)}\mathbf{W}_{\alpha\delta}^{(2)}{}' &= \mathbf{W}_{\alpha\delta}\mathbf{W}_{\alpha\delta}' + \mathbf{W}_{\delta\alpha}\mathbf{W}_{\delta\alpha}' \\ &= \{[(\mathbf{W}_{\alpha}\mathbf{W}_{\alpha}')\#(\mathbf{W}_{\delta}\mathbf{W}_{\delta}') - (\mathbf{W}_{\alpha}\#\mathbf{W}_{\delta})(\mathbf{W}_{\alpha}\#\mathbf{W}_{\delta})'] \\ &\quad + [(\mathbf{W}_{\alpha}\mathbf{W}_{\alpha}')\#(\mathbf{W}_{\delta}\mathbf{W}_{\delta}') - (\mathbf{W}_{\delta}\#\mathbf{W}_{\alpha})(\mathbf{W}_{\delta}\#\mathbf{W}_{\alpha}\#)']\}/2 \\ &= [(\mathbf{W}_{\alpha}\mathbf{W}_{\alpha}')\#(\mathbf{W}_{\delta}\mathbf{W}_{\delta}') - (\mathbf{W}_{\alpha}\#\mathbf{W}_{\delta})(\mathbf{W}_{\alpha}\#\mathbf{W}_{\delta})'] \end{aligned} \quad \text{(S1.15)}$$

$$\mathbf{W}_{\delta\delta}\mathbf{W}_{\delta\delta}' = \frac{1}{2}[(\mathbf{W}_{\delta}\mathbf{W}_{\delta}')\#(\mathbf{W}_{\delta}\mathbf{W}_{\delta}') - (\mathbf{W}_{\delta}\#\mathbf{W}_{\delta})(\mathbf{W}_{\delta}\#\mathbf{W}_{\delta})'] \quad \text{(S1.16)}$$

In Equations S1.14-S1.16, SNP additive and dominance model matrices are used, and no epistasis model matrix needs to be created. In Equation 3.1.34, $\mathbf{W}_{\alpha}\#\mathbf{W}_{\delta} = \mathbf{W}_{\delta}\#\mathbf{W}_{\alpha}$.

**Pairwise intra- and inter-chromosome EGERM**

Based on the products of the epistasis model matrices, the numerators of intra- and inter-chromosome pairwise EGERM are calculated as:

$$\begin{aligned} \mathbf{W}_{\alpha\alpha}^{\text{intra}}(\mathbf{W}_{\alpha\alpha}^{\text{intra}})' &= \textstyle\sum_{i=1}^{ch}(\mathbf{W}_{\alpha\alpha}^{i}\mathbf{W}_{\alpha\alpha}^{i}{}') \\ &= \frac{1}{2}\textstyle\sum_{i=1}^{ch}[(\mathbf{W}_{\alpha}^{i}\mathbf{W}_{\alpha}^{i}{}')\#(\mathbf{W}_{\alpha}^{i}\mathbf{W}_{\alpha}^{i}{}') - (\mathbf{W}_{\alpha}^{i}\#\mathbf{W}_{\alpha}^{i})(\mathbf{W}_{\alpha}^{i}\#\mathbf{W}_{\alpha}^{i})'] \end{aligned} \quad \text{(S1.17)}$$

$$\mathbf{W}_{\alpha\alpha}^{\text{inter}}(\mathbf{W}_{\alpha\alpha}^{\text{inter}})' = \mathbf{W}_{\alpha\alpha}\mathbf{W}_{\alpha\alpha}' - \mathbf{W}_{\alpha\alpha}^{\text{intra}}(\mathbf{W}_{\alpha\alpha}^{\text{intra}})' \quad \text{(S1.18)}$$

$$\begin{aligned} \mathbf{W}_{\alpha\delta}^{(2)\text{intra}}[\mathbf{W}_{\alpha\delta}^{(2)\text{intra}}]' &= \textstyle\sum_{i=1}^{ch}(\mathbf{W}_{\alpha\delta}^{i}\mathbf{W}_{\alpha\delta}^{i}{}' + \mathbf{W}_{\delta\alpha}^{i}\mathbf{W}_{\delta\alpha}^{i}{}') \\ &= \textstyle\sum_{i=1}^{ch}[(\mathbf{W}_{\alpha}^{i}\mathbf{W}_{\alpha}^{i}{}')\#(\mathbf{W}_{\delta}^{i}\mathbf{W}_{\delta}^{i}{}') - (\mathbf{W}_{\alpha}^{i}\#\mathbf{W}_{\delta}^{i})(\mathbf{W}_{\alpha}^{i}\#\mathbf{W}_{\delta}^{i})'] \end{aligned} \quad \text{(S1.19)}$$

$$\mathbf{W}_{\alpha\delta}^{(2)\text{inter}}[\mathbf{W}_{\alpha\delta}^{(2)\text{inter}}]' = \mathbf{W}_{\alpha\delta}^{(2)}\mathbf{W}_{\alpha\delta}^{(2)}{}' - \mathbf{W}_{\alpha\delta}^{(2)\text{intra}}[\mathbf{W}_{\alpha\delta}^{(2)\text{intra}}]' \quad \text{(S1.20)}$$

$$\begin{aligned} \mathbf{W}_{\delta\delta}^{\text{intra}}(\mathbf{W}_{\delta\delta}^{\text{intra}})' &= \textstyle\sum_{i=1}^{ch}(\mathbf{W}_{\delta}^{i}\mathbf{W}_{\delta}^{i}{}') \\ &= \frac{1}{2}\textstyle\sum_{i=1}^{ch}[(\mathbf{W}_{\delta}^{i}\mathbf{W}_{\delta}^{i}{}')\#(\mathbf{W}_{\delta}^{i}\mathbf{W}_{\delta}^{i}{}') - (\mathbf{W}_{\delta}^{i}\#\mathbf{W}_{\delta}^{i})(\mathbf{W}_{\delta}^{i}\#\mathbf{W}_{\delta}^{i})'] \end{aligned} \quad \text{(S1.21)}$$

$$\mathbf{W}_{\delta\delta}^{\text{inter}}(\mathbf{W}_{\delta\delta}^{\text{inter}})' = \mathbf{W}_{\delta\delta}\mathbf{W}_{\delta\delta}' - \mathbf{W}_{\delta\delta}^{\text{intra}}(\mathbf{W}_{\delta\delta}^{\text{intra}})' \quad \text{(S1.22)}$$

where ch = number of chromosomes. The intra- and inter-chromosome pairwise EGERM are:

$$\mathbf{S}_4 = \mathbf{S}_4^{intra} = \mathbf{W}_{\alpha\alpha}^{intra}(\mathbf{W}_{\alpha\alpha}^{intra})'/k_{\alpha\alpha}^{intra} \quad (S1.23)$$
$$\mathbf{S}_5 = \mathbf{S}_4^{inter} = \mathbf{W}_{\alpha\alpha}^{inter}(\mathbf{W}_{\alpha\alpha}^{inter})'/k_{\alpha\alpha}^{inter} \quad (S1.24)$$
$$\mathbf{S}_6 = \mathbf{S}_5^{intra} = \mathbf{W}_{\alpha\delta}^{(2)intra}[\mathbf{W}_{\alpha\delta}^{(2)intra}]'/k_{\alpha\delta}^{intra} \quad (S1.25)$$
$$\mathbf{S}_7 = \mathbf{S}_5^{inter} = \mathbf{W}_{\alpha\delta}^{(2)inter}[\mathbf{W}_{\alpha\delta}^{(2)inter}]'/k_{\alpha\delta}^{inter} \quad (S1.26)$$
$$\mathbf{S}_8 = \mathbf{S}_6^{intra} = \mathbf{W}_{\delta\delta}^{intra}(\mathbf{W}_{\delta\delta}^{intra})'/k_{\delta\delta}^{intra} \quad (S1.27)$$
$$\mathbf{S}_9 = \mathbf{S}_6^{inter} = \mathbf{W}_{\delta\delta}^{inter}(\mathbf{W}_{\delta\delta}^{inter})'/k_{\alpha\alpha}^{inter} \quad (S1.28)$$

where

$$k_{\alpha\alpha}^{intra} = tr[\mathbf{W}_{\alpha\alpha}^{intra}(\mathbf{W}_{\alpha\alpha}^{intra})']/n \quad (S1.29)$$
$$\text{with } k_{\alpha\alpha}^{inter} = tr[\mathbf{W}_{\alpha\alpha}^{inter}(\mathbf{W}_{\alpha\alpha}^{inter})']/n \quad (S1.30)$$
$$\text{with } k_{\alpha\delta}^{intra} = tr\{\mathbf{W}_{\alpha\delta}^{(2)intra}[\mathbf{W}_{\alpha\delta}^{(2)intra}]'\}/n \quad (S1.31)$$
$$\text{with } k_{\alpha\delta}^{inter} = tr\{\mathbf{W}_{\alpha\delta}^{(2)inter}[\mathbf{W}_{\alpha\delta}^{(2)inter}]'\}/n \quad (S1.32)$$
$$\text{with } k_{\delta\delta}^{intra} = tr[\mathbf{W}_{\delta\delta}^{intra}(\mathbf{W}_{\delta\delta}^{intra})']/n \quad (S1.33)$$
$$\text{with } k_{\alpha\alpha}^{inter} = tr[\mathbf{W}_{\delta\delta}^{inter}(\mathbf{W}_{\delta\delta}^{inter})']/n \quad (S1.34)$$

As shown by Equations S1.14-S1.22, the calculations of Equations S1.23-S1.34 are based on the model matrices of SNP additive and dominance effects and do not involve the epistasis model matrices.

**Approximate genomic epistasis relationship matrices (AGERM)**

Let A = pedigree additive relationship matrix, and D = pedigree dominance relationship matrix. Then, Henderson defined each epistasis relationship matrix as a Hadamard product of A and D matrices [15]. For Model-I:

$$\mathbf{S}_4 = \mathbf{S}_{AA} = \mathbf{A}\#\mathbf{A} = \text{A×A relationship matrix} \quad (S1.35)$$
$$\mathbf{S}_5 = \mathbf{S}_{AD} = \mathbf{A}\#\mathbf{D} = \text{A×D relationship matrix} \quad (S1.36)$$
$$\mathbf{S}_6 = \mathbf{S}_{DD} = \mathbf{D}\#\mathbf{D} = \text{D×D relationship matrix} \quad (S1.37)$$
$$\mathbf{S}_7 = \mathbf{S}_{AAA} = \mathbf{A}\#\mathbf{A}\#\mathbf{A} = \text{A×A×A relationship matrix} \quad (S1.38)$$
$$\mathbf{S}_8 = \mathbf{S}_{AAD} = \mathbf{A}\#\mathbf{A}\#\mathbf{D} = \text{A×A×D relationship matrix} \quad (S1.39)$$
$$\mathbf{S}_9 = \mathbf{S}_{ADD} = \mathbf{A}\#\mathbf{D}\#\mathbf{D} = \text{A×D×D relationship matrix} \quad (S1.40)$$
$$\mathbf{S}_{10} = \mathbf{S}_{DDD} = \mathbf{D}\#\mathbf{D}\#\mathbf{D} = \text{D×D×D relationship matrix} \quad (S1.41)$$

Approximate genomic epistasis relationship matrices are the genomic version of Henderson's epistasis relationship matrices with the pedigree **A** and **D** matrices replaced by the genomic additive and dominance relationship matrices [9]. Using notations in this article, genomic additive and dominance relationship matrices that are the genomic versions of pedigree additive and dominance relationship matrices are:

$$\mathbf{A} = \mathbf{S}_1 = \mathbf{S}_\alpha = \mathbf{W}_\alpha\mathbf{W}_\alpha'/(2\textstyle\sum_{i=1}^m p_iq_i) = \mathbf{W}_1\mathbf{W}_1'/(2\textstyle\sum_{i=1}^m p_iq_i) \quad (S1.42)$$

$$\mathbf{D} = \mathbf{S}_2 = \mathbf{S}_\delta = \mathbf{W}_\delta\mathbf{W}_\delta'/(4\textstyle\sum_{i=1}^m p_i^2q_i^2) = \mathbf{W}_2\mathbf{W}_2'/(4\textstyle\sum_{i=1}^m p_i^2q_i^2) \quad (S1.43)$$

Replacing the **A** and **D** in Equations S1.35-S1.41 with those of Equations S1.42 and S1.43 yields the AGERM.

**Pairwise intra- and inter-chromosome AGERM**

For Model-II, the numerators of the intra- and inter-chromosome AGERM can be derived from those of AGERM. In Equations S1.14-S1.16, the minus term is for removing intra-locus epistasis that is contained in AGERM but should exist. Therefore, removing the minus terms of Equations S1.14-S1.16 yields the numerators of AGERM, i.e.:

$$\mathbf{W}_{\alpha\alpha}\mathbf{W}_{\alpha\alpha}' = \frac{1}{2}(\mathbf{W}_{\alpha}\mathbf{W}_{\alpha}')\#(\mathbf{W}_{\alpha}\mathbf{W}_{\alpha}') \quad \text{(S1.44)}$$

$$\mathbf{W}_{\alpha\delta}^{(2)}\mathbf{W}_{\alpha\delta}^{(2)}{}' = \mathbf{W}_{\alpha\delta}\mathbf{W}_{\alpha\delta}' + \mathbf{W}_{\delta\alpha}\mathbf{W}_{\delta\alpha}' = (\mathbf{W}_{\alpha}\mathbf{W}_{\alpha}')\#(\mathbf{W}_{\delta}\mathbf{W}_{\delta}') \quad \text{(S1.45)}$$

$$\mathbf{W}_{\delta\delta}\mathbf{W}_{\delta\delta}' = \frac{1}{2}(\mathbf{W}_{\delta}\mathbf{W}_{\delta}')\#(\mathbf{W}_{\delta}\mathbf{W}_{\delta}') \quad \text{(S1.46)}$$

noting that the left-hand-side of Equations S1.14-S1.16 and S1.23-S1.25 are the same, but the right-hand-side are different. Similarly, the numerators of the intra-chromosome epistasis relationships based on the additive and dominance model matrix of each chromosome, i.e.:

$$\mathbf{W}_{\alpha\alpha}^{\text{intra}}(\mathbf{W}_{\alpha\alpha}^{\text{intra}})' = \sum_{i=1}^{ch}(\mathbf{W}_{\alpha\alpha}^{i}\mathbf{W}_{\alpha\alpha}^{i}{}') = \frac{1}{2}\sum_{i=1}^{ch}(\mathbf{W}_{\alpha}^{i}\mathbf{W}_{\alpha}^{i}{}')\#(\mathbf{W}_{\alpha}^{i}\mathbf{W}_{\alpha}^{i}{}') \quad \text{(S1.47)}$$

$$\mathbf{W}_{\alpha\delta}^{(2)\text{intra}}[\mathbf{W}_{\alpha\delta}^{(2)\text{intra}}]' = \sum_{i=1}^{ch}(\mathbf{W}_{\alpha\delta}^{i}\mathbf{W}_{\alpha\delta}^{i}{}' + \mathbf{W}_{\delta\alpha}^{i}\mathbf{W}_{\delta\alpha}^{i}{}') = \sum_{i=1}^{ch}(\mathbf{W}_{\alpha}^{i}\mathbf{W}_{\alpha}^{i}{}')\#(\mathbf{W}_{\delta}^{i}\mathbf{W}_{\delta}^{i}{}') \quad \text{(S1.48)}$$

$$\mathbf{W}_{\delta\delta}^{\text{intra}}(\mathbf{W}_{\delta\delta}^{\text{intra}})' = \sum_{i=1}^{ch}(\mathbf{W}_{\delta}^{i}\mathbf{W}_{\delta}^{i}{}') = \frac{1}{2}\sum_{i=1}^{ch}(\mathbf{W}_{\delta}^{i}\mathbf{W}_{\delta}^{i}{}')\#(\mathbf{W}_{\delta}^{i}\mathbf{W}_{\delta}^{i}{}') \quad \text{(S1.49)}$$

where ch = number of chromosomes. Then, the numerators of the inter-chromosome epistasis relationship matrices are calculated as the difference between the numerators of the $\mathbf{A}\#\mathbf{A}$, $\mathbf{A}\#\mathbf{D}$ and $\mathbf{D}\#\mathbf{D}$ genomic relationships matrices and the numerators of the intra-chromosome relationship matrices, i.e.:

$$\begin{aligned}\mathbf{W}_{\alpha\alpha}^{\text{inter}}(\mathbf{W}_{\alpha\alpha}^{\text{inter}})' &= \mathbf{W}_{\alpha\alpha}\mathbf{W}_{\alpha\alpha}' - \mathbf{W}_{\alpha\alpha}^{\text{intra}}(\mathbf{W}_{\alpha\alpha}^{\text{intra}})' \\ &= \frac{1}{2}[(\mathbf{W}_{\alpha}\mathbf{W}_{\alpha}')\#(\mathbf{W}_{\alpha}\mathbf{W}_{\alpha}') - \sum_{i=1}^{ch}(\mathbf{W}_{\alpha}^{i}\mathbf{W}_{\alpha}^{i}{}')\#(\mathbf{W}_{\alpha}^{i}\mathbf{W}_{\alpha}^{i}{}')]\end{aligned} \quad \text{(S1.50)}$$

$$\mathbf{W}_{\alpha\delta}^{(2)\text{inter}}[\mathbf{W}_{\alpha\delta}^{(2)\text{inter}}]' = \mathbf{W}_{\alpha\delta}^{(2)}\mathbf{W}_{\alpha\delta}^{(2)}{}' - \mathbf{W}_{\alpha\delta}^{(2)\text{intra}}[\mathbf{W}_{\alpha\delta}^{(2)\text{intra}}]' \quad \text{(S1.51)}$$

$$\mathbf{W}_{\delta\delta}^{\text{inter}}(\mathbf{W}_{\delta\delta}^{\text{inter}})' = \mathbf{W}_{\delta\delta}\mathbf{W}_{\delta\delta}' - \mathbf{W}_{\delta\delta}^{\text{intra}}(\mathbf{W}_{\delta\delta}^{\text{intra}})' \quad \text{(S1.52)}$$

Then, the pairwise intra- and inter-chromosome AGERM are:

$$\mathbf{S}_4 = \mathbf{S}_4^{\text{intra}} = \mathbf{W}_{\alpha\alpha}^{\text{intra}}(\mathbf{W}_{\alpha\alpha}^{\text{intra}})'/k_{\alpha\alpha}^{\text{intra}} = [\sum_{i=1}^{ch}(\mathbf{W}_{\alpha}^{i}\mathbf{W}_{\alpha}^{i}{}')\#(\mathbf{W}_{\alpha}^{i}\mathbf{W}_{\alpha}^{i}{}')]/k_{\alpha\alpha}^{\text{intra}} \quad \text{(S1.53)}$$

= intra-chromosome A×A AGERM

$$\mathbf{S}_5 = \mathbf{S}_4^{\text{inter}} = \mathbf{W}_{\alpha\alpha}^{\text{inter}}(\mathbf{W}_{\alpha\alpha}^{\text{inter}})'/k_{\alpha\alpha}^{\text{inter}} = [\sum_{i=1}^{ch}(\mathbf{W}_{\alpha}^{i}\mathbf{W}_{\alpha}^{i}{}')\#(\mathbf{W}_{\delta}^{i}\mathbf{W}_{\delta}^{i}{}')]/k_{\alpha\alpha}^{\text{inter}} \quad \text{(S1.54)}$$

= inter-chromosome A×A AGERM

$$\mathbf{S}_6 = \mathbf{S}_5^{\text{intra}} = \mathbf{W}_{\alpha\delta}^{(2)\text{intra}}[\mathbf{W}_{\alpha\delta}^{(2)\text{intra}}]'/k_{\alpha\delta}^{\text{intra}} = [\sum_{i=1}^{ch}(\mathbf{W}_{\delta}^{i}\mathbf{W}_{\delta}^{i}{}')\#(\mathbf{W}_{\delta}^{i}\mathbf{W}_{\delta}^{i}{}')]/k_{\alpha\delta}^{\text{intra}} \quad \text{(S1.55)}$$

= intra-chromosome A×D AGERM

$$\mathbf{S}_7 = \mathbf{S}_5^{\text{inter}} = \mathbf{W}_{\alpha\delta}^{(2)\text{inter}}[\mathbf{W}_{\alpha\delta}^{(2)\text{inter}}]'/k_{\alpha\delta}^{\text{inter}} = \{\mathbf{W}_{\alpha\delta}^{(2)}\mathbf{W}_{\alpha\delta}^{(2)}{}' - \mathbf{W}_{\alpha\delta}^{(2)\text{intra}}[\mathbf{W}_{\alpha\delta}^{(2)\text{intra}}]'\}/k_{\alpha\delta}^{\text{inter}} \quad \text{(S1.56)}$$

= inter-chromosome A×D AGERM

$$\mathbf{S}_8 = \mathbf{S}_6^{intra} = \mathbf{W}_{\delta\delta}^{intra}(\mathbf{W}_{\delta\delta}^{intra})'/k_{\delta\delta}^{intra} = \frac{1}{2}[\sum_{i=1}^{ch}(\mathbf{W}_\delta^i\mathbf{W}_\delta^{i}{}')\#(\mathbf{W}_\delta^i\mathbf{W}_\delta^{i}{}')]/k_{\delta\delta}^{intra} \quad (S1.57)$$

= intra-chromosome D×D AGERM

$$\mathbf{S}_9 = \mathbf{S}_6^{inter} = \mathbf{W}_{\delta\delta}^{inter}(\mathbf{W}_{\delta\delta}^{inter})'/k_{\alpha\alpha}^{inter} = [\mathbf{W}_{\delta\delta}\mathbf{W}_{\delta\delta}' - \mathbf{W}_{\delta\delta}^{intra}(\mathbf{W}_{\delta\delta}^{intra})'] \quad (S1.58)$$

= inter-chromosome D×D AGERM

where

$$k_{\alpha\alpha}^{intra} = tr[\mathbf{W}_{\alpha\alpha}^{intra}(\mathbf{W}_{\alpha\alpha}^{intra})']/n = tr[\sum_{i=1}^{ch}(\mathbf{W}_\alpha^i\mathbf{W}_\alpha^{i}{}')\#(\mathbf{W}_\alpha^i\mathbf{W}_\alpha^{i}{}')]/n \quad (S1.59)$$

$$k_{\alpha\alpha}^{inter} = tr[\mathbf{W}_{\alpha\alpha}^{inter}(\mathbf{W}_{\alpha\alpha}^{inter})']/n = \frac{1}{2}tr[(\mathbf{W}_\alpha\mathbf{W}_\alpha')\#(\mathbf{W}_\alpha\mathbf{W}_\alpha') - \sum_{i=1}^{ch}(\mathbf{W}_\alpha^i\mathbf{W}_\alpha^{i}{}')\#(\mathbf{W}_\alpha^i\mathbf{W}_\alpha^{i}{}')]/n \quad (S1.60)$$

$$k_{\alpha\delta}^{intra} = tr\{\mathbf{W}_{\alpha\delta}^{(2)intra}[\mathbf{W}_{\alpha\delta}^{(2)intra}]'\}/n = tr[\sum_{i=1}^{ch}(\mathbf{W}_\delta^i\mathbf{W}_\delta^{i}{}')\#(\mathbf{W}_\delta^i\mathbf{W}_\delta^{i}{}')]/n \quad (S1.61)$$

$$k_{\alpha\delta}^{inter} = tr\{\mathbf{W}_{\alpha\delta}^{(2)inter}[\mathbf{W}_{\alpha\delta}^{(2)inter}]'\}/n = tr\{\mathbf{W}_{\alpha\delta}^{(2)}\mathbf{W}_{\alpha\delta}^{(2)}{}' - \mathbf{W}_{\alpha\delta}^{(2)intra}[\mathbf{W}_{\alpha\delta}^{(2)intra}]'\}/n \quad (S1.62)$$

$$k_{\delta\delta}^{intra} = tr[\mathbf{W}_{\delta\delta}^{intra}(\mathbf{W}_{\delta\delta}^{intra})']/n = \frac{1}{2}tr[\sum_{i=1}^{ch}(\mathbf{W}_\delta^i\mathbf{W}_\delta^{i}{}')\#(\mathbf{W}_\delta^i\mathbf{W}_\delta^{i}{}')]/n \quad (S1.63)$$

$$k_{\alpha\alpha}^{inter} = tr[\mathbf{W}_{\delta\delta}^{inter}(\mathbf{W}_{\delta\delta}^{inter})']/n = tr[\mathbf{W}_{\delta\delta}\mathbf{W}_{\delta\delta}' - \mathbf{W}_{\delta\delta}^{intra}(\mathbf{W}_{\delta\delta}^{intra})']/n \quad (S1.64)$$

Note that AGERM of Equations S1.53-S1.58 and EGERM of Equations S1.23-S1.28 have the same general expression defined by Equations S1.12-S1.13 but the right-hand-side of the EGERM and AGERM are different.

### Text S2. Numerical Demonstration

This is a summary of the results from DEMO.R program for numerical demonstration of the methods in this article. The small dataset is for the convenience of reading the numerical results. The haplotype data is adopted from the numerical example in Da (2015) and is not derived from the small SNP data set, which does not have any accuracy for imputing haplotypes. Detailed procedure to produce haplotype data from the SNP data is described in the GVHHAP computing pipeline (Prakapenka et al., 2020). The SNP and haplotype data are completely artificial for showing calculations only without any genetic or methodology implications.

**Phenotypic values, SNP and haplotype genotypes, fixed effects**

The dataset in in this demo consists of 5 individuals with one missing phenotypic value for individual #5.
The column vector of the phenotypic values for the four individuals with phenotypic observations is:

$$\mathbf{y} = (1.753222, 1.617655, 1.713119, 1.738835)'.$$

The fixed non-genetic effects are male and female, with individuals 1 and 2 having a sex code of '2', and individuals 3 and 4 having a sex code of '1'. The **X** matrix for these fixed effects is:

| | | |
|---|---|---|
| 1 | 0 | 1 |
| 2 | 0 | 1 |
| 3 | 1 | 0 |
| 4 | 1 | 0. |

Each individual has one phenotypic observation, so the **Z** matrix is an identity matrix.

Each individual has 6 SNP markers:

```
#6 MARKERS, 5 INDIVIDUALS
# LOC1:  0-0-1-0-1
# LOC2:  2-0-1-1-1
# LOC3:  0-0-1-0-1
# LOC4:  1-1-2-0-1
# LOC5:  0-2-0-1-1
# LOC6:  2-0-1-1-0
```

where '0' indicate a homozygous genotype (AA), '2' indicates the other homozygous genotype (aa), and '1' indicates the heterozygous genotype (Aa) of the SNP.

The 5 individuals have one haplotype block that is treated as a 'locus' with four haplotypes that each was treated as an 'allele'. The haplotype frequencies of the four haplotypes are 0.4, 0.3, 0.2 and 0.1 for haplotypes 1, 2 3 and 4 respectively.

**Calculation of allele frequencies**

Lines 42-75 in DEMO.R calculate the allele frequencies of the 6 SNPs from the SNP genotypes of the 5 individuals.

**SNP additive and dominance codings**

Lines 78-100 in DEMO.R calculate the additive coding for each SNP genotype, and lines 110-132 calculate the dominance coding for each SNP genotype using formulae described in the main text.

**Model matrices of SNP additive and dominance effects**

Lines 102-107 and line 144 in DEMO.R calculate the SNP additive model matrix, and lines 110-139 and line 145 calculate the SNP dominance model matrix.

$\mathbf{W}_{\alpha}$ ($\mathbf{W}_1$):

| | SNP1 | SNP2 | SNP3 | SNP4 | SNP5 | SNP6 |
|---|---|---|---|---|---|---|
| ID1 | 0.4 | -1 | 0.4 | 0 | 0.8 | -1.2 |
| ID2 | 0.4 | 1 | 0.4 | 0 | -1.2 | 0.8 |
| ID3 | -0.6 | 0 | -0.6 | -1 | 0.8 | -0.2 |
| ID4 | 0.4 | 0 | 0.4 | 1 | -0.2 | -0.2 |
| ID5 | -0.6 | 0 | -0.6 | 0 | -0.2 | 0.8 |

$\mathbf{W}_{\delta}$ ($\mathbf{W}_2$):

| | SNP1 | SNP2 | SNP3 | SNP4 | SNP5 | SNP6 |
|---|---|---|---|---|---|---|
| ID1 | -0.08 | -0.5 | -0.08 | 0.5 | -0.32 | -0.72 |
| ID2 | -0.08 | -0.5 | -0.08 | 0.5 | -0.72 | -0.32 |
| ID3 | 0.32 | 0.5 | 0.32 | -0.5 | -0.32 | 0.48 |
| ID4 | -0.08 | 0.5 | -0.08 | -0.5 | 0.48 | 0.48 |
| ID5 | 0.32 | 0.5 | 0.32 | 0.5 | 0.48 | -0.32 |

**Haplotype additive codings**

Lines 519-541 in DEMO.R calculate the haplotype additive coding for each genotype of two pairing haplotypes. The haplotype data is artificial for showing calculations only.

**Model matrix of haplotype additive effects**

Lines 546 in DEMO.R calculates the haplotype additive model matrix.

$\mathbf{W}_{\alpha h}$ ($\mathbf{W}_3$):

| | HAP1 | HAP2 | HAP3 |
|---|---|---|---|
| ID1 | 0.6 | 0.4 | 0.2 |
| ID2 | -1.4 | 0.4 | 0.2 |
| ID3 | 0.6 | -1.6 | 0.2 |
| ID4 | 0.6 | 0.4 | -1.8 |
| ID5 | -0.4 | 0.4 | 0.2 |

**Pairwise epistasis**
Lines 152-501 in DEMO.R calculates the pairwise epistasis model matrices.

$\mathbf{W}_{\alpha\alpha}$ ($\mathbf{W}_4$ under Model-I):

| | SNP1_2 | SNP1_3 | SNP1_4 | SNP1_5 | SNP1_6 |
|---|---|---|---|---|---|
| ID1 | -0.4 | 0.16 | 0 | 0.32 | -0.48 |
| ID2 | 0.4 | 0.16 | 0 | -0.48 | 0.32 |
| ID3 | 0 | 0.36 | 0.6 | -0.48 | 0.12 |
| ID4 | 0 | 0.16 | 0.4 | -0.08 | -0.08 |
| ID5 | 0 | 0.36 | 0 | 0.12 | -0.48 |

| | SNP2_3 | SNP2_4 | SNP2_5 | SNP2_6 | SNP3_4 |
|---|---|---|---|---|---|
| ID1 | -0.4 | 0 | -0.8 | 1.2 | 0 |
| ID2 | 0.4 | 0 | -1.2 | 0.8 | 0 |
| ID3 | 0 | 0 | 0 | 0 | 0.6 |
| ID4 | 0 | 0 | 0 | 0 | 0.4 |
| ID5 | 0 | 0 | 0 | 0 | 0 |

| | SNP3_5 | SNP3_6 | SNP4_5 | SNP4_6 | SNP5_6 |
|---|---|---|---|---|---|
| ID1 | 0.32 | -0.48 | 0 | 0 | -0.96 |
| ID2 | -0.48 | 0.32 | 0 | 0 | -0.96 |
| ID3 | -0.48 | 0.12 | -0.8 | 0.2 | -0.16 |
| ID4 | -0.08 | -0.08 | -0.2 | -0.2 | 0.04 |
| ID5 | 0.12 | -0.48 | 0 | 0 | -0.16 |

$\mathbf{W}_{\alpha\delta}$ (part of $\mathbf{W}_5$ under Model-I):

| | SNP1_2 | SNP1_3 | SNP1_4 | SNP1_5 | SNP1_6 |
|---|---|---|---|---|---|
| ID1 | -0.2 | -0.032 | 0.2 | -0.128 | -0.288 |
| ID2 | -0.2 | -0.032 | 0.2 | -0.288 | -0.128 |
| ID3 | -0.3 | -0.192 | 0.3 | 0.192 | -0.288 |
| ID4 | 0.2 | -0.032 | -0.2 | 0.192 | 0.192 |
| ID5 | -0.3 | -0.192 | -0.3 | -0.288 | 0.192 |

| | SNP2_3 | SNP2_4 | SNP2_5 | SNP2_6 | SNP3_4 |
|---|---|---|---|---|---|
| ID1 | 0.08 | -0.5 | 0.32 | 0.72 | 0.2 |
| ID2 | -0.08 | 0.5 | -0.72 | -0.32 | 0.2 |
| ID3 | 0 | 0 | 0 | 0 | 0.3 |
| ID4 | 0 | 0 | 0 | 0 | -0.2 |
| ID5 | 0 | 0 | 0 | 0 | -0.3 |

| | SNP3_5 | SNP3_6 | SNP4_5 | SNP4_6 | SNP5_6 |
|---|---|---|---|---|---|
| ID1 | -0.128 | -0.288 | 0 | 0 | -0.576 |
| ID2 | -0.288 | -0.128 | 0 | 0 | 0.384 |
| ID3 | 0.192 | -0.288 | 0.32 | -0.48 | 0.384 |

| | | | | | |
|---|---|---|---|---|---|
| ID4 | 0.192 | 0.192 | 0.48 | 0.48 | -0.096 |
| ID5 | -0.288 | 0.192 | 0 | 0 | 0.064 |

$\mathbf{W}_{\delta\alpha}$ (part of $\mathbf{W}_5$ under Model-I):

| | SNP1_2 | SNP1_3 | SNP1_4 | SNP1_5 | SNP1_6 |
|---|---|---|---|---|---|
| ID1 | 0.08 | -0.032 | 0 | -0.064 | 0.096 |
| ID2 | -0.08 | -0.032 | 0 | 0.096 | -0.064 |
| ID3 | 0 | -0.192 | -0.32 | 0.256 | -0.064 |
| ID4 | 0 | -0.032 | -0.08 | 0.016 | 0.016 |
| ID5 | 0 | -0.192 | 0 | -0.064 | 0.256 |

| | SNP2_3 | SNP2_4 | SNP2_5 | SNP2_6 | SNP3_4 |
|---|---|---|---|---|---|
| ID1 | -0.2 | 0 | -0.4 | 0.6 | 0 |
| ID2 | -0.2 | 0 | 0.6 | -0.4 | 0 |
| ID3 | -0.3 | -0.5 | 0.4 | -0.1 | -0.32 |
| ID4 | 0.2 | 0.5 | -0.1 | -0.1 | -0.08 |
| ID5 | -0.3 | 0 | -0.1 | 0.4 | 0 |

| | SNP3_5 | SNP3_6 | SNP4_5 | SNP4_6 | SNP5_6 |
|---|---|---|---|---|---|
| ID1 | -0.064 | 0.096 | 0.4 | -0.6 | 0.384 |
| ID2 | 0.096 | -0.064 | -0.6 | 0.4 | -0.576 |
| ID3 | 0.256 | -0.064 | -0.4 | 0.1 | 0.064 |
| ID4 | 0.016 | 0.016 | 0.1 | 0.1 | -0.096 |
| ID5 | -0.064 | 0.256 | -0.1 | 0.4 | 0.384 |

$\mathbf{W}_{\delta\delta}$ ($\mathbf{W}_6$ under Model-I):

| | SNP1_2 | SNP1_3 | SNP1_4 | SNP1_5 | SNP1_6 |
|---|---|---|---|---|---|
| ID1 | 0.04 | 0.0064 | -0.04 | 0.0256 | 0.0576 |
| ID2 | 0.04 | 0.0064 | -0.04 | 0.0576 | 0.0256 |
| ID3 | 0.16 | 0.1024 | -0.16 | -0.1024 | 0.1536 |
| ID4 | -0.04 | 0.0064 | 0.04 | -0.0384 | -0.0384 |
| ID5 | 0.16 | 0.1024 | 0.16 | 0.1536 | -0.1024 |

| | SNP2_3 | SNP2_4 | SNP2_5 | SNP2_6 | SNP3_4 |
|---|---|---|---|---|---|
| ID1 | 0.04 | -0.25 | 0.16 | 0.36 | -0.04 |
| ID2 | 0.04 | -0.25 | 0.36 | 0.16 | -0.04 |
| ID3 | 0.16 | -0.25 | -0.16 | 0.24 | -0.16 |
| ID4 | -0.04 | -0.25 | 0.24 | 0.24 | 0.04 |
| ID5 | 0.16 | 0.25 | 0.24 | -0.16 | 0.16 |

| | SNP3_5 | SNP3_6 | SNP4_5 | SNP4_6 | SNP5_6 |
|---|---|---|---|---|---|
| ID1 | 0.0256 | 0.0576 | -0.16 | -0.36 | 0.2304 |
| ID2 | 0.0576 | 0.0256 | -0.36 | -0.16 | 0.2304 |
| ID3 | -0.1024 | 0.1536 | 0.16 | -0.24 | -0.1536 |
| ID4 | -0.0384 | -0.0384 | -0.24 | -0.24 | 0.2304 |

| ID5 | 0.1536 | -0.1024 | 0.24 | -0.16 | -0.1536 |
|---|---|---|---|---|---|

**Additive and dominance genomic relationship matrices**

Lines 552-554 in DEMO.R calculate the additive genomic matrix, and lines 559-561 calculate the dominance genomic matrix.

$\mathbf{S}_{\alpha}$ ($\mathbf{S}_1$):

| | ID1 | ID2 | ID3 | ID4 | ID5 |
|---|---|---|---|---|---|
| ID1 | 1.4166667 | -1.0833333 | 0.1666667 | 0.166667 | -0.6666667 |
| ID2 | -1.0833333 | 1.4166667 | -0.6666667 | 0.166667 | 0.1666667 |
| ID3 | 0.1666667 | -0.6666667 | 1 | -0.66667 | 0.1666667 |
| ID4 | 0.1666667 | 0.1666667 | -0.6666667 | 0.583333 | -0.25 |
| ID5 | -0.6666667 | 0.1666667 | 0.1666667 | -0.25 | 0.5833333 |

$\mathbf{S}_{\delta}$ ($\mathbf{S}_2$):

| | ID1 | ID2 | ID3 | ID4 | ID5 |
|---|---|---|---|---|---|
| ID1 | 1.0662152 | 0.9157261 | -0.74717833 | -0.92777 | 0.02407825 |
| ID2 | 0.91572611 | 1.0662152 | -0.44620015 | -0.92777 | -0.27689992 |
| ID3 | -0.74717833 | -0.4462002 | 0.97592175 | 0.494357 | -0.09631302 |
| ID4 | -0.92776524 | -0.9277652 | 0.49435666 | 0.915726 | 0.02407825 |
| ID5 | 0.02407825 | -0.2768999 | -0.09631302 | 0.024078 | 0.97592175 |

**Haplotype additive genomic relationship matrix**

Lines 593-595 in DEMO.R calculate the haplotype additive genomic relationship matrix.

$\mathbf{S}_{\alpha h}$ ($\mathbf{S}_3$):

| | ID1 | ID2 | ID3 | ID4 | ID5 |
|---|---|---|---|---|---|
| ID1 | 0.28571429 | -0.3265306 | -0.122449 | 0.081633 | -0.02040816 |
| ID2 | -0.32653061 | 1.1020408 | -0.7346939 | -0.53061 | 0.3877551 |
| ID3 | -0.12244898 | -0.7346939 | 1.5102041 | -0.32653 | -0.42857143 |
| ID4 | 0.08163265 | -0.5306122 | -0.3265306 | 1.918367 | -0.2244898 |
| ID5 | -0.02040816 | 0.3877551 | -0.4285714 | -0.22449 | 0.18367347 |

**Pairwise epistasis genomic relationship matrices calculated from epistasis model matrices**
Lines 566-588 in DEMO.R calculates the pairwise epistasis genomic relationship matrices using the epistasis model matrices.

$\mathbf{S}_{\alpha\alpha}$ ($\mathbf{S}_4$ under Model-I):

| | ID1 | ID2 | ID3 | ID4 | ID5 |
|---|---|---|---|---|---|
| ID1 | 1.796561605 | 0.865329513 | -0.09455587 | 0.005731 | 0.33524355 |
| ID2 | 0.865329513 | 1.796561605 | 0.33524355 | 0.005731 | -0.09455587 |
| ID3 | -0.094555874 | 0.335243553 | 0.91547278 | 0.317335 | -0.03366762 |
| ID4 | 0.005730659 | 0.005730659 | 0.31733524 | 0.202722 | 0.0487106 |
| ID5 | 0.335243553 | -0.094555874 | -0.03366762 | 0.048711 | 0.28868195 |

$\mathbf{S}_{\alpha\delta}$ ($\mathbf{S}_5$ under Model-I):

| | ID1 | ID2 | ID3 | ID4 | ID5 |
|---|---|---|---|---|---|
| ID1 | 1.38554743 | -0.9142301 | -0.15525187 | -0.16796 | 0.07119021 |
| ID2 | -0.9142301 | 1.38554743 | 0.46845702 | -0.16796 | -0.05593517 |
| ID3 | -0.15525187 | 0.46845702 | 1.03873351 | -0.31634 | -0.08136024 |
| ID4 | -0.1679644 | -0.1679644 | -0.31634356 | 0.542547 | -0.03051009 |
| ID5 | 0.07119021 | -0.05593517 | -0.08136024 | -0.03051 | 0.64762434 |

$\mathbf{S}_{\delta\delta}$ ($\mathbf{S}_6$ under Model-I):

| | ID1 | ID2 | ID3 | ID4 | ID5 |
|---|---|---|---|---|---|
| ID1 | 1.0454473 | 0.85066432 | 0.4449322 | 0.836686 | -0.24017 |
| ID2 | 0.8506643 | 1.04544735 | 0.02619429 | 0.836686 | -0.2012741 |
| ID3 | 0.4449322 | 0.02619429 | 1.02922049 | 0.086969 | -0.2168325 |
| ID4 | 0.8366861 | 0.83668614 | 0.08696902 | 0.850664 | -0.24017 |
| ID5 | -0.24017 | -0.20127414 | -0.21683247 | -0.24017 | 1.0292205 |

**GREML Estimation of the variance components under Model-I (EM-REML)**
Lines 613-651 in DEMO.R estimate variance components (excluding third-order variance components) using EM-REML for one iteration.

| | |
|---|---|
| Variance_component_A | 5.29E-04 |
| Variance_component_D | 1.04E-04 |
| Variance_component_AA | 1.68E-03 |
| Variance_component_AD2 | 8.79E-04 |
| Variance_component_DD | 6.97E-05 |
| Variance_component_AH | 2.92E-04 |
| Variance_component_E | 1.48E-04 |

**Heritability estimate for each effect type under Model-I**

Lines 657-664 in DEMO.R estimate variance components (excluding third-order variance components) using EM-REML for one iteration.

| | |
|---|---|
| Heritability_A | 0.143042 |
| Heritability_D | 0.028118 |
| Heritability_AA | 0.453261 |
| Heritability_AD2 | 0.23773 |
| Heritability_DD | 0.018861 |
| Heritability_AH | 0.079069 |
| Heritability_G | 0.96008 |

**GBLUP and reliability**

Lines 672-708 in DEMO.R calculate GBLUP and reliability (excluding third-order epistasis values).

| | Gblup_A | Reliabily_A | Gblup_D | Reliabily_D |
|---|---|---|---|---|
| ID1 | 0.0030404 | -0.007875 | 0.0098429 | 0.0012264 |
| ID2 | 0.0047594 | 0.0046453 | -0.000713 | 0.0069185 |
| ID3 | -0.000412 | 0.0012506 | 0.0134085 | -0.03868 |
| ID4 | -0.001009 | 0.0008768 | 0.0013323 | 0.002898 |
| ID5 | -0.024791 | 0.0226529 | 0.0400866 | 0.0171162 |

| | Gblup_AA | Reliabily_AA | Gblup_AD | Reliabily_AD |
|---|---|---|---|---|
| ID1 | -0.004475 | -3.92E-04 | -0.003651 | -0.001009 |
| ID2 | -0.003026 | 9.19E-03 | 0.002585 | -0.024791 |
| ID3 | 0.0440865 | 6.98E-02 | 0.0291752 | 0.0030404 |
| ID4 | -0.001553 | 4.74E-03 | 0.031094 | 0.0047594 |
| ID5 | 0.000632 | -9.70E-05 | 0.0009413 | -0.000412 |

| | Gblup_DD | Reliabily_DD |
|---|---|---|
| ID1 | 0.0008768 | 0.0013323 |
| ID2 | 0.0226529 | 0.0400866 |
| ID3 | -0.007875 | 0.0098429 |
| ID4 | 0.0046453 | -0.000713 |
| ID5 | 0.0012506 | 0.0134085 |

| | Gblup_AH | Reliabily_AH | Gblup_G | Reliabily_G |
|---|---|---|---|---|
| ID1 | 0.002898 | -0.001553 | 4.74E-03 | 0.031094 |
| ID2 | 0.0171162 | 0.000632 | -9.70E-05 | 0.0009413 |
| ID3 | 0.0012264 | -0.004475 | -3.92E-04 | -0.003651 |
| ID4 | 0.0069185 | -0.003026 | 9.19E-03 | 0.002585 |
| ID5 | -0.03868 | 0.0440865 | 6.98E-02 | 0.0291752 |

**Effect and heritability for each level of an effect type**
Lines 717-750 in DEMO.R calculate the effect and heritability of each level of an effect type (excluding third-order variance components).

| | mrk_eff_A | mrk_eff_D |
|---|---|---|
| SNP1 | -8.14E-03 | 0.002033998 |
| SNP2 | 9.74E-04 | -0.006038177 |
| SNP3 | -2.13E-05 | -0.010138082 |
| SNP4 | 1.68E-03 | -0.001188418 |
| SNP5 | -2.03E-03 | -0.002473631 |
| SNP6 | 7.02E-04 | 0.000274859 |

| | mrk_eff_AH |
|---|---|
| HAP1 | 0.00518 |
| HAP2 | 0.001074 |
| HAP3 | -0.00107 |

| | mrk_eff_AA | mrk_eff_AD | mrk_eff_DA | mrk_eff_DD |
|---|---|---|---|---|
| SNP1_2 | -0.006141951 | 0.009212927 | 0.0149407 | 0.005354 |
| SNP1_3 | -0.002303232 | 0.004549962 | -0.022301 | -0.006885 |
| SNP1_4 | 0.00077178 | 0.010358489 | 0.0146719 | 0.0030952 |
| SNP1_5 | 0.001145562 | -0.000763708 | -0.004582 | -0.000863 |
| SNP1_6 | 0.023119953 | -0.001548663 | -0.010549 | 0.0050232 |
| SNP2_3 | 0.013940898 | -0.013649829 | 0.01388 | -0.003493 |
| SNP2_4 | -0.012133598 | 0.005018198 | -0.00523 | -0.005184 |
| SNP2_5 | 0.003456305 | 0.003456305 | 0.0023885 | -0.003583 |
| SNP2_6 | -0.02487345 | 0.00729717 | 0.0023885 | -0.012274 |
| SNP3_4 | 0.015425695 | 0.014643962 | -0.002176 | -0.012618 |
| SNP3_5 | 0.020501353 | -0.001227232 | 0.0081324 | -0.009043 |
| SNP3_6 | 0.02345172 | -0.006625493 | -0.007864 | -0.003981 |
| SNP4_5 | -0.002607671 | 0.001998911 | 0.0013763 | 0.0071227 |
| SNP4_6 | -0.011030802 | 0.005416921 | 0.0115912 | 0.0065652 |
| SNP5_6 | 0.002057625 | 0.001433362 | 0.0054789 | -0.011279 |

| | h2_A | h2_D |
|---|---|---|
| SNP1 | 0.000536 | 3.91E-04 |
| SNP2 | 0.049876 | 1.43E-30 |
| SNP3 | 0.000536 | 3.91E-04 |
| SNP4 | 0.002142 | 1.43E-30 |
| SNP5 | 0.040075 | 1.82E-02 |
| SNP6 | 0.049876 | 9.11E-03 |

| | h2_AH |
|---|---|
| HAP1 | 0.072813 |

| | |
|---|---|
| HAP2 | 0.003128 |
| HAP3 | 0.003128 |

| | h2_AA | h2_AD | h2_DA | h2_DD |
|---|---|---|---|---|
| SNP1_2 | 0.063003 | 0.000574 | 1.10E-03 | 2.66E-04 |
| SNP1_3 | 0.000169 | 5.88E-05 | 4.73E-05 | 6.13E-05 |
| SNP1_4 | 0.000169 | 0.000574 | 1.06E-04 | 2.66E-04 |
| SNP1_5 | 0.076738 | 0.00137 | 1.89E-03 | 5.43E-05 |
| SNP1_6 | 0.069701 | 0.000196 | 1.34E-03 | 9.40E-06 |
| SNP2_3 | 0.063003 | 0.00137 | 4.62E-04 | 2.66E-04 |
| SNP2_4 | 0 | 0.053497 | 1.85E-03 | 9.21E-33 |
| SNP2_5 | 0.015751 | 0.057862 | 5.24E-02 | 2.12E-03 |
| SNP2_6 | 0.015751 | 0.057862 | 4.30E-02 | 6.19E-03 |
| SNP3_4 | 0.000169 | 0.000574 | 1.06E-04 | 2.66E-04 |
| SNP3_5 | 0.076738 | 0.00137 | 1.89E-03 | 5.43E-05 |
| SNP3_6 | 0.069701 | 0.000196 | 1.34E-03 | 9.40E-06 |
| SNP4_5 | 0.001522 | 5.88E-05 | 5.24E-02 | 2.12E-03 |
| SNP4_6 | 0.000677 | 0.002118 | 4.30E-02 | 6.19E-03 |
| SNP5_6 | 0.000169 | 0.06005 | 3.69E-02 | 9.80E-04 |

**Pairwise epistasis genomic relationship matrices calculated from additive and dominance model matrices**

Lines 761-780 in DEMO.R calculate the pairwise epistasis genomic relationship matrices using the SNP additive and dominance model matrices. The resulting epistasis model matrices are identical to those calculated from the epistasis model matrices.

**Third-order epistasis effects, Lines 784-1245**

Line 784-930 calculate the epistasis model matrices of the third-order epistasis effects.

Lines 951-980 calculate third-order epistasis genomic relationship matrices.

$\mathbf{S}_{\alpha\alpha\alpha}$ ($\mathbf{S}_7$):

| | ID1 | ID2 | ID3 | ID4 | ID5 |
|---|---|---|---|---|---|
| ID1 | 2.02779616 | -0.09000662 | -0.02382528 | -0.01058901 | -0.14295169 |
| ID2 | -0.09000662 | 2.02779616 | -0.14295169 | -0.01058901 | -0.02382528 |
| ID3 | -0.02382528 | -0.14295169 | 0.77696889 | -0.11647915 | -0.02382528 |
| ID4 | -0.01058901 | -0.01058901 | -0.11647915 | 0.05724686 | -0.00397088 |
| ID5 | -0.14295169 | -0.02382528 | -0.02382528 | -0.00397088 | 0.11019193 |

$\mathbf{S}_{\alpha\alpha\delta}$ ($\mathbf{S}_8$):

| | ID1 | ID2 | ID3 | ID4 | ID5 |
|---|---|---|---|---|---|
| ID1 | 1.659640724 | 0.550220613 | 0.13641916 | -0.009827415 | -0.05075649 |
| ID2 | 0.550220613 | 1.659640724 | -0.32112132 | -0.009827415 | 0.06016241 |
| ID3 | 0.136419159 | -0.321121316 | 1.07177285 | 0.209244351 | -0.01644558 |
| ID4 | -0.009827415 | -0.009827415 | 0.20924435 | 0.226198768 | 0.01727377 |
| ID5 | -0.050756490 | 0.060162413 | -0.01644558 | 0.017273770 | 0.38274693 |

**$S_{\alpha\delta\delta}$ ($S_9$):**

| | ID1 | ID2 | ID3 | ID4 | ID5 |
|---|---|---|---|---|---|
| ID1 | 1.659640724 | 0.550220613 | 0.13641916 | -0.009827415 | -0.05075649 |
| ID2 | 0.550220613 | 1.659640724 | -0.32112132 | -0.009827415 | 0.06016241 |
| ID3 | 0.136419159 | -0.321121316 | 1.07177285 | 0.209244351 | -0.01644558 |
| ID4 | -0.009827415 | -0.009827415 | 0.20924435 | 0.226198768 | 0.01727377 |
| ID5 | -0.050756490 | 0.060162413 | -0.01644558 | 0.017273770 | 0.38274693 |

**$S_{\delta\delta\delta}$ ($S_{10}$):**

| | ID1 | ID2 | ID3 | ID4 | ID5 |
|---|---|---|---|---|---|
| ID1 | 0.913906125 | 0.7716022 | -0.07232653 | -0.685148497 | 0.003383527 |
| ID2 | 0.771602240 | 0.9139061 | 0.29408068 | -0.685148497 | 0.258695683 |
| ID3 | -0.072326535 | 0.2940807 | 1.20029276 | -0.204747972 | 0.103343980 |
| ID4 | -0.685148497 | -0.6851485 | -0.20474797 | 0.771602240 | 0.003383527 |
| ID5 | 0.003383527 | 0.2586957 | 0.10334398 | 0.003383527 | 1.200292755 |

Lines 1026-1078 calculate variance components and heritabilities using EM-REML for 1 iteration.

| | |
|---|---|
| Variance_component_A | 0.000206 |
| Variance_component_D | 0.000050 |
| Variance_component_AA | 0.000717 |
| Variance_component_AD2 | 0.000366 |
| Variance_component_DD | 0.000027 |
| Variance_component_AAA | 0.000921 |
| Variance_component_AAD3 | 0.000543 |
| Variance_component_ADD3 | 0.000399 |
| Variance_component_DDD | 0.000088 |
| Variance_component_AH | 0.000124 |
| Variance_component_E | 0.000060 |

| | |
|---|---|
| Heritability_A | 0.05894 |
| Heritability_D | 0.01426 |
| Heritability_AA | 0.20477 |
| Heritability_AD2 | 0.10451 |
| Heritability_DD | 0.00782 |
| Heritability_AAA | 0.26307 |
| Heritability_AAD3 | 0.15512 |
| Heritability_ADD3 | 0.11401 |
| Heritability_DDD | 0.02509 |
| Heritability_AH | 0.03531 |
| Heritability_G | 0.98289 |

Lines 1082-1134 calculate GBLUP and reliability.

| | Gblup_A | Rel_A | Gblup_D | Rel_D | Gblup_AA | Rel_AA |
|---|---|---|---|---|---|---|
| ID1 | -0.02246 | -0.30616 | 0.001853 | 0.004462 | 0.014082 | 0.054619 |
| ID2 | 0.023339 | -0.37694 | -0.00102 | 0.011056 | -0.01367 | 0.052242 |
| ID3 | -0.00924 | -0.22103 | -0.00264 | 0.014952 | -0.00551 | 0.038634 |
| ID4 | 0.001313 | -0.12264 | -0.00053 | 0.01067 | 0.000205 | 0.005057 |
| ID5 | 0.00705 | -0.07668 | 0.003098 | 0.007093 | 0.006435 | 0.070652 |

| Gblup_AD2 | Rel_AD2 | Gblup_DD | Rel_DD | Gblup_AAA | Rel_AAA |
|---|---|---|---|---|---|
| 0.076042 | 0.976758 | 0.000476 | 2.43E-03 | -0.01736 | -0.13975 |
| -0.07353 | 0.895727 | -0.00092 | 1.29E-02 | 0.017504 | -0.14467 |
| -0.01537 | 0.294887 | 0.00169 | 2.27E-02 | -0.00183 | -0.04823 |
| -0.00332 | 0.228947 | -0.00029 | 1.12E-02 | 0.000167 | -0.02204 |
| 0.004004 | 0.005834 | -0.00012 | 3.46E-05 | 0.000996 | -0.00896 |

| Gblup_AAD3 | Rel_AAD3 | Gblup_ADD3 | Rel_ADD3 |
|---|---|---|---|

| | | | |
|---|---|---|---|
| -1.48E-02 | -8.08E-02 | -0.01557 | -0.18822 |
| 1.51E-02 | -8.92E-02 | 0.015996 | -0.2158 |
| -7.34E-03 | -8.40E-02 | -0.00466 | -0.07356 |
| 2.61E-05 | -8.51E-05 | 0.000499 | -0.02289 |
| 1.51E-03 | -3.93E-03 | -0.00136 | -0.00275 |

| Gblup_DDD | Rel_DDD | Gblup_AH | Rel_AH | Gblup_G | Rel_G |
|---|---|---|---|---|---|
| 0.002141 | 0.026164 | 0.017765 | 0.28006 | 0.042157 | 0.303021 |
| -0.00028 | 0.049642 | -0.04385 | 0.466862 | -0.06138 | 0.501699 |
| -0.00202 | 0.076826 | 0.024969 | 0.483807 | -0.02197 | 0.462695 |
| -0.00114 | 0.063008 | 0.010561 | 0.370612 | 0.007494 | 0.477525 |
| -0.00244 | 0.003844 | -0.01304 | 0.296484 | 0.006135 | 0.019598 |

Lines 1143-1169 calculate third-order epistasis genomic relationship matrices using additive and dominance model matrices. The results are the same as the direct calculations using the third-order epistasis model matrices.

**Heritability estimates from EPIHAP**

All results of DEMO.R are verified by the EIPHAP program (Liang et al., 2021), and DEMO.R and EPIHAP have identical results, which provide strong mutual validation for the correct implementation of the methods in this article. The following only the heritability estimates from EPIHAP without repeating all identical results.

Additive heritability: 5.893893e-02

Dominance heritability: 1.425995e-02

Additive x Additive heritability: 2.047676e-01

Additive x Dominance heritability: 1.045092e-01

Dominance x Dominance heritability: 7.814957e-03

Additive x Additive x Additive heritability: 2.630716e-01

Additive x Additive x Dominance heritability: 1.551156e-01

Additive x Dominance x Dominance heritability: 1.140088e-01

Dominance x Dominance x Dominance heritability: 2.509076e-02

Additive Haplotype heritability: 3.530722e-02

Total heritability: 9.828847e-01

#### R Program. DEMO.R for Numerical Demonstration

```
# Program name: DEMO.R
# Authors: Zuoxiang Liang, Dzianis Prakapenka, Yang Da. November 11, 2021.
# Purpose: numerial demonstration
# 1. Pairwise epistasis genomic relationship matrices
#    using epistasis model matrices
# 2. GREML estimation of variance components and heritability
# 3. GBLUP and reliability values
# 4. Effect and hertiability estimates for all levels of all effect types
# 5. Pairwise epistasis genomic relationship matrices
#    using additive and dominance model matrices
# 6. GREML and GBLUP with third-order epistasis effects

# Genotype
# 6 markers, 5 individuals

# LOC1:  0-0-1-0-1
# LOC2:  2-0-1-1-1
# LOC3:  0-0-1-0-1
# LOC4:  1-1-2-0-1
# LOC5:  0-2-0-1-1
# LOC6:  2-0-1-1-0

# Phenotype
# 5 individuals

# ID  FIX   y
# ID1 2     1.753222323
# ID2 2     1.617654738
# ID3 1     1.71311938
# ID4 1     1.738835045
# ID5 -9999 -9999
# -9999 indicates missing value, and FIX means fixed effects

library(matrixcalc)
```

```
library(MASS)
library(pedigree)

ID<-c("ID1","ID2","ID3","ID4","ID5")
FIX<-c(2,2,1,1,1)
y<-c(1.753222323,1.617654738,1.71311938,1.738835045,-9999)

# 1. Genomic relationship matrices of AA, AD and DD using epistasis model matrices

# CALCULATION OF ALLELE FREQUENCIES
# THE NEXT LINE IS LINE 42 REFERENCED BY TEXT S4
P1_II = 3/5
P1_IJ = 2/5
P1_JJ = 0/5
P1_I = P1_II +0.5*P1_IJ
P1_J = P1_JJ + 0.5*P1_IJ

P2_II = 1/5
P2_IJ = 3/5
P2_JJ = 1/5
P2_I = P2_II +0.5*P2_IJ
P2_J = P2_JJ + 0.5*P2_IJ
P3_II = 3/5
P3_IJ = 2/5
P3_JJ = 0/5
P3_I = P3_II +0.5*P3_IJ
P3_J = P3_JJ + 0.5*P3_IJ

P4_II = 1/5
P4_IJ = 3/5
P4_JJ = 1/5
P4_I = P4_II +0.5*P4_IJ
P4_J = P4_JJ + 0.5*P4_IJ

P5_II = 2/5
```

```
P5_IJ = 2/5
P5_JJ = 1/5
P5_I = P5_II +0.5*P5_IJ
P5_J = P5_JJ + 0.5*P5_IJ

P6_II = 2/5
P6_IJ = 2/5
P6_JJ = 1/5
P6_I = P6_II +0.5*P6_IJ
P6_J = P6_JJ + 0.5*P6_IJ

#ADDITIVE SNP CODING
WA1_0 = 2*P1_J
WA1_1 = P1_J - P1_I
WA1_2 = -2*P1_I

WA2_0 = 2*P2_J
WA2_1 = P2_J - P2_I
WA2_2 = -2*P2_I

WA3_0 = 2*P3_J
WA3_1 = P3_J - P3_I
WA3_2 = -2*P3_I

WA4_0 = 2*P4_J
WA4_1 = P4_J - P4_I
WA4_2 = -2*P4_I

WA5_0 = 2*P5_J
WA5_1 = P5_J - P5_I
WA5_2 = -2*P5_I

WA6_0 = 2*P6_J
WA6_1 = P6_J - P6_I
WA6_2 = -2*P6_I
```

```
WA1 =  rbind(WA1_0,WA1_0,WA1_1,WA1_0,WA1_1)
WA2 =  rbind(WA2_2,WA2_0,WA2_1,WA2_1,WA2_1)
WA3 =  rbind(WA3_0,WA3_0,WA3_1,WA3_0,WA3_1)
WA4 =  rbind(WA4_1,WA4_1,WA4_2,WA4_0,WA4_1)
WA5 =  rbind(WA5_0,WA5_2,WA5_0,WA5_1,WA5_1)
WA6 =  rbind(WA6_2,WA6_0,WA6_1,WA6_1,WA6_0)

# DOMINANCE SNP CODING
WD1_0 = -2*P1_J**2
WD1_1 = 2*P1_I*P1_J
WD1_2 = -2*P1_I**2

WD2_0 = -2*P2_J**2
WD2_1 = 2*P2_I*P2_J
WD2_2 = -2*P2_I**2

WD3_0 = -2*P3_J**2
WD3_1 = 2*P3_I*P3_J
WD3_2 = -2*P3_I**2

WD4_0 = -2*P4_J**2
WD4_1 = 2*P4_I*P4_J
WD4_2 = -2*P4_I**2

WD5_0 = -2*P5_J**2
WD5_1 = 2*P5_I*P5_J
WD5_2 = -2*P5_I**2

WD6_0 = -2*P6_J**2
WD6_1 = 2*P6_I*P6_J
WD6_2 = -2*P6_I**2

WD1 = rbind(WD1_0,WD1_0,WD1_1,WD1_0,WD1_1)
WD2 = rbind(WD2_2,WD2_0,WD2_1,WD2_1,WD2_1)
```

```
WD3 = rbind(WD3_0,WD3_0,WD3_1,WD3_0,WD3_1)

WD4 = rbind(WD4_1,WD4_1,WD4_2,WD4_0,WD4_1)

WD5 = rbind(WD5_0,WD5_2,WD5_0,WD5_1,WD5_1)

WD6 = rbind(WD6_2,WD6_0,WD6_1,WD6_1,WD6_0)

rownames<-ID

SNPs_name<-c("SNP1","SNP2","SNP3","SNP4","SNP5","SNP6")

colnames<-SNPs_name

WA<-
matrix(c(WA1,WA2,WA3,WA4,WA5,WA6),nrow=length(rownames),ncol=length(colnames),byrow=
FALSE,dimnames=list(rownames, colnames))

WD<-
matrix(c(WD1,WD2,WD3,WD4,WD5,WD6),nrow=length(rownames),ncol=length(colnames),byrow=
FALSE,dimnames=list(rownames, colnames))

print("WA matrix")

print(WA)

print("WD matrix")

print(WD)

# PAIRWISE

WAA1_12=WA1_0*WA2_2

WAA1_13=WA1_0*WA3_0

WAA1_14=WA1_0*WA4_1

WAA1_15=WA1_0*WA5_0

WAA1_16=WA1_0*WA6_2

WAA1_23=WA2_2*WA3_0

WAA1_24=WA2_2*WA4_1

WAA1_25=WA2_2*WA5_0

WAA1_26=WA2_2*WA6_2

WAA1_34=WA3_0*WA4_1

WAA1_35=WA3_0*WA5_0

WAA1_36=WA3_0*WA6_2

WAA1_45=WA4_1*WA5_0

WAA1_46=WA4_1*WA6_2

WAA1_56=WA5_0*WA6_2

WAD1_12=WA1_0*WD2_2
```

```
WAD1_13=WA1_0*WD3_0

WAD1_14=WA1_0*WD4_1

WAD1_15=WA1_0*WD5_0

WAD1_16=WA1_0*WD6_2

WAD1_23=WA2_2*WD3_0

WAD1_24=WA2_2*WD4_1

WAD1_25=WA2_2*WD5_0

WAD1_26=WA2_2*WD6_2

WAD1_34=WA3_0*WD4_1

WAD1_35=WA3_0*WD5_0

WAD1_36=WA3_0*WD6_2

WAD1_45=WA4_1*WD5_0

WAD1_46=WA4_1*WD6_2

WAD1_56=WA5_0*WD6_2

WDA1_12=WD1_0*WA2_2

WDA1_13=WD1_0*WA3_0

WDA1_14=WD1_0*WA4_1

WDA1_15=WD1_0*WA5_0

WDA1_16=WD1_0*WA6_2

WDA1_23=WD2_2*WA3_0

WDA1_24=WD2_2*WA4_1

WDA1_25=WD2_2*WA5_0

WDA1_26=WD2_2*WA6_2

WDA1_34=WD3_0*WA4_1

WDA1_35=WD3_0*WA5_0

WDA1_36=WD3_0*WA6_2

WDA1_45=WD4_1*WA5_0

WDA1_46=WD4_1*WA6_2

WDA1_56=WD5_0*WA6_2

WDD1_12=WD1_0*WD2_2

WDD1_13=WD1_0*WD3_0

WDD1_14=WD1_0*WD4_1

WDD1_15=WD1_0*WD5_0
```

```
WDD1_16=WD1_0*WD6_2
WDD1_23=WD2_2*WD3_0
WDD1_24=WD2_2*WD4_1
WDD1_25=WD2_2*WD5_0
WDD1_26=WD2_2*WD6_2
WDD1_34=WD3_0*WD4_1
WDD1_35=WD3_0*WD5_0
WDD1_36=WD3_0*WD6_2
WDD1_45=WD4_1*WD5_0
WDD1_46=WD4_1*WD6_2
WDD1_56=WD5_0*WD6_2

WAA2_12=WA1_0*WA2_0
WAA2_13=WA1_0*WA3_0
WAA2_14=WA1_0*WA4_1
WAA2_15=WA1_0*WA5_2
WAA2_16=WA1_0*WA6_0
WAA2_23=WA2_0*WA3_0
WAA2_24=WA2_0*WA4_1
WAA2_25=WA2_0*WA5_2
WAA2_26=WA2_0*WA6_0
WAA2_34=WA3_0*WA4_1
WAA2_35=WA3_0*WA5_2
WAA2_36=WA3_0*WA6_0
WAA2_45=WA4_1*WA5_2
WAA2_46=WA4_1*WA6_0
WAA2_56=WA5_2*WA6_0

WAD2_12=WA1_0*WD2_0
WAD2_13=WA1_0*WD3_0
WAD2_14=WA1_0*WD4_1
WAD2_15=WA1_0*WD5_2
WAD2_16=WA1_0*WD6_0
WAD2_23=WA2_0*WD3_0
WAD2_24=WA2_0*WD4_1
```

```
WAD2_25=WA2_0*WD5_2
WAD2_26=WA2_0*WD6_0
WAD2_34=WA3_0*WD4_1
WAD2_35=WA3_0*WD5_2
WAD2_36=WA3_0*WD6_0
WAD2_45=WA4_1*WD5_2
WAD2_46=WA4_1*WD6_0
WAD2_56=WA5_2*WD6_0

WDA2_12=WD1_0*WA2_0
WDA2_13=WD1_0*WA3_0
WDA2_14=WD1_0*WA4_1
WDA2_15=WD1_0*WA5_2
WDA2_16=WD1_0*WA6_0
WDA2_23=WD2_0*WA3_0
WDA2_24=WD2_0*WA4_1
WDA2_25=WD2_0*WA5_2
WDA2_26=WD2_0*WA6_0
WDA2_34=WD3_0*WA4_1
WDA2_35=WD3_0*WA5_2
WDA2_36=WD3_0*WA6_0
WDA2_45=WD4_1*WA5_2
WDA2_46=WD4_1*WA6_0
WDA2_56=WD5_2*WA6_0

WDD2_12=WD1_0*WD2_0
WDD2_13=WD1_0*WD3_0
WDD2_14=WD1_0*WD4_1
WDD2_15=WD1_0*WD5_2
WDD2_16=WD1_0*WD6_0
WDD2_23=WD2_0*WD3_0
WDD2_24=WD2_0*WD4_1
WDD2_25=WD2_0*WD5_2
WDD2_26=WD2_0*WD6_0
WDD2_34=WD3_0*WD4_1
```

```
WDD2_35=WD3_0*WD5_2
WDD2_36=WD3_0*WD6_0
WDD2_45=WD4_1*WD5_2
WDD2_46=WD4_1*WD6_0
WDD2_56=WD5_2*WD6_0

WAA3_12=WA1_1*WA2_1
WAA3_13=WA1_1*WA3_1
WAA3_14=WA1_1*WA4_2
WAA3_15=WA1_1*WA5_0
WAA3_16=WA1_1*WA6_1
WAA3_23=WA2_1*WA3_1
WAA3_24=WA2_1*WA4_2
WAA3_25=WA2_1*WA5_0
WAA3_26=WA2_1*WA6_1
WAA3_34=WA3_1*WA4_2
WAA3_35=WA3_1*WA5_0
WAA3_36=WA3_1*WA6_1
WAA3_45=WA4_2*WA5_0
WAA3_46=WA4_2*WA6_1
WAA3_56=WA5_0*WA6_1

WAD3_12=WA1_1*WD2_1
WAD3_13=WA1_1*WD3_1
WAD3_14=WA1_1*WD4_2
WAD3_15=WA1_1*WD5_0
WAD3_16=WA1_1*WD6_1
WAD3_23=WA2_1*WD3_1
WAD3_24=WA2_1*WD4_2
WAD3_25=WA2_1*WD5_0
WAD3_26=WA2_1*WD6_1
WAD3_34=WA3_1*WD4_2
WAD3_35=WA3_1*WD5_0
WAD3_36=WA3_1*WD6_1
WAD3_45=WA4_2*WD5_0
```

```
WAD3_46=WA4_2*WD6_1
WAD3_56=WA5_0*WD6_1

WDA3_12=WD1_1*WA2_1
WDA3_13=WD1_1*WA3_1
WDA3_14=WD1_1*WA4_2
WDA3_15=WD1_1*WA5_0
WDA3_16=WD1_1*WA6_1
WDA3_23=WD2_1*WA3_1
WDA3_24=WD2_1*WA4_2
WDA3_25=WD2_1*WA5_0
WDA3_26=WD2_1*WA6_1
WDA3_34=WD3_1*WA4_2
WDA3_35=WD3_1*WA5_0
WDA3_36=WD3_1*WA6_1
WDA3_45=WD4_2*WA5_0
WDA3_46=WD4_2*WA6_1
WDA3_56=WD5_0*WA6_1

WDD3_12=WD1_1*WD2_1
WDD3_13=WD1_1*WD3_1
WDD3_14=WD1_1*WD4_2
WDD3_15=WD1_1*WD5_0
WDD3_16=WD1_1*WD6_1
WDD3_23=WD2_1*WD3_1
WDD3_24=WD2_1*WD4_2
WDD3_25=WD2_1*WD5_0
WDD3_26=WD2_1*WD6_1
WDD3_34=WD3_1*WD4_2
WDD3_35=WD3_1*WD5_0
WDD3_36=WD3_1*WD6_1
WDD3_45=WD4_2*WD5_0
WDD3_46=WD4_2*WD6_1
WDD3_56=WD5_0*WD6_1
```

```
WAA4_12=WA1_0*WA2_1
WAA4_13=WA1_0*WA3_0
WAA4_14=WA1_0*WA4_0
WAA4_15=WA1_0*WA5_1
WAA4_16=WA1_0*WA6_1
WAA4_23=WA2_1*WA3_0
WAA4_24=WA2_1*WA4_0
WAA4_25=WA2_1*WA5_1
WAA4_26=WA2_1*WA6_1
WAA4_34=WA3_0*WA4_0
WAA4_35=WA3_0*WA5_1
WAA4_36=WA3_0*WA6_1
WAA4_45=WA4_0*WA5_1
WAA4_46=WA4_0*WA6_1
WAA4_56=WA5_1*WA6_1

WAD4_12=WA1_0*WD2_1
WAD4_13=WA1_0*WD3_0
WAD4_14=WA1_0*WD4_0
WAD4_15=WA1_0*WD5_1
WAD4_16=WA1_0*WD6_1
WAD4_23=WA2_1*WD3_0
WAD4_24=WA2_1*WD4_0
WAD4_25=WA2_1*WD5_1
WAD4_26=WA2_1*WD6_1
WAD4_34=WA3_0*WD4_0
WAD4_35=WA3_0*WD5_1
WAD4_36=WA3_0*WD6_1
WAD4_45=WA4_0*WD5_1
WAD4_46=WA4_0*WD6_1
WAD4_56=WA5_1*WD6_1

WDA4_12=WD1_0*WA2_1
WDA4_13=WD1_0*WA3_0
WDA4_14=WD1_0*WA4_0
```

```
WDA4_15=WD1_0*WA5_1
WDA4_16=WD1_0*WA6_1
WDA4_23=WD2_1*WA3_0
WDA4_24=WD2_1*WA4_0
WDA4_25=WD2_1*WA5_1
WDA4_26=WD2_1*WA6_1
WDA4_34=WD3_0*WA4_0
WDA4_35=WD3_0*WA5_1
WDA4_36=WD3_0*WA6_1
WDA4_45=WD4_0*WA5_1
WDA4_46=WD4_0*WA6_1
WDA4_56=WD5_1*WA6_1

WDD4_12=WD1_0*WD2_1
WDD4_13=WD1_0*WD3_0
WDD4_14=WD1_0*WD4_0
WDD4_15=WD1_0*WD5_1
WDD4_16=WD1_0*WD6_1
WDD4_23=WD2_1*WD3_0
WDD4_24=WD2_1*WD4_0
WDD4_25=WD2_1*WD5_1
WDD4_26=WD2_1*WD6_1
WDD4_34=WD3_0*WD4_0
WDD4_35=WD3_0*WD5_1
WDD4_36=WD3_0*WD6_1
WDD4_45=WD4_0*WD5_1
WDD4_46=WD4_0*WD6_1
WDD4_56=WD5_1*WD6_1

WAA5_12=WA1_1*WA2_1
WAA5_13=WA1_1*WA3_1
WAA5_14=WA1_1*WA4_1
WAA5_15=WA1_1*WA5_1
WAA5_16=WA1_1*WA6_0
WAA5_23=WA2_1*WA3_1
```

```
WAA5_24=WA2_1*WA4_1
WAA5_25=WA2_1*WA5_1
WAA5_26=WA2_1*WA6_0
WAA5_34=WA3_1*WA4_1
WAA5_35=WA3_1*WA5_1
WAA5_36=WA3_1*WA6_0
WAA5_45=WA4_1*WA5_1
WAA5_46=WA4_1*WA6_0
WAA5_56=WA5_1*WA6_0

WAD5_12=WA1_1*WD2_1
WAD5_13=WA1_1*WD3_1
WAD5_14=WA1_1*WD4_1
WAD5_15=WA1_1*WD5_1
WAD5_16=WA1_1*WD6_0
WAD5_23=WA2_1*WD3_1
WAD5_24=WA2_1*WD4_1
WAD5_25=WA2_1*WD5_1
WAD5_26=WA2_1*WD6_0
WAD5_34=WA3_1*WD4_1
WAD5_35=WA3_1*WD5_1
WAD5_36=WA3_1*WD6_0
WAD5_45=WA4_1*WD5_1
WAD5_46=WA4_1*WD6_0
WAD5_56=WA5_1*WD6_0

WDA5_12=WD1_1*WA2_1
WDA5_13=WD1_1*WA3_1
WDA5_14=WD1_1*WA4_1
WDA5_15=WD1_1*WA5_1
WDA5_16=WD1_1*WA6_0
WDA5_23=WD2_1*WA3_1
WDA5_24=WD2_1*WA4_1
WDA5_25=WD2_1*WA5_1
WDA5_26=WD2_1*WA6_0
```

```
WDA5_34=WD3_1*WA4_1

WDA5_35=WD3_1*WA5_1

WDA5_36=WD3_1*WA6_0

WDA5_45=WD4_1*WA5_1

WDA5_46=WD4_1*WA6_0

WDA5_56=WD5_1*WA6_0

WDD5_12=WD1_1*WD2_1

WDD5_13=WD1_1*WD3_1

WDD5_14=WD1_1*WD4_1

WDD5_15=WD1_1*WD5_1

WDD5_16=WD1_1*WD6_0

WDD5_23=WD2_1*WD3_1

WDD5_24=WD2_1*WD4_1

WDD5_25=WD2_1*WD5_1

WDD5_26=WD2_1*WD6_0

WDD5_34=WD3_1*WD4_1

WDD5_35=WD3_1*WD5_1

WDD5_36=WD3_1*WD6_0

WDD5_45=WD4_1*WD5_1

WDD5_46=WD4_1*WD6_0

WDD5_56=WD5_1*WD6_0

WAA1=cbind(WAA1_12,WAA1_13,WAA1_14,WAA1_15,WAA1_16,WAA1_23,WAA1_24,WAA1_25,WAA1_26,W
AA1_34,WAA1_35,WAA1_36,WAA1_45,WAA1_46,WAA1_56)

WAA2=cbind(WAA2_12,WAA2_13,WAA2_14,WAA2_15,WAA2_16,WAA2_23,WAA2_24,WAA2_25,WAA2_26,W
AA2_34,WAA2_35,WAA2_36,WAA2_45,WAA2_46,WAA2_56)

WAA3=cbind(WAA3_12,WAA3_13,WAA3_14,WAA3_15,WAA3_16,WAA3_23,WAA3_24,WAA3_25,WAA3_26,W
AA3_34,WAA3_35,WAA3_36,WAA3_45,WAA3_46,WAA3_56)

WAA4=cbind(WAA4_12,WAA4_13,WAA4_14,WAA4_15,WAA4_16,WAA4_23,WAA4_24,WAA4_25,WAA4_26,W
AA4_34,WAA4_35,WAA4_36,WAA4_45,WAA4_46,WAA4_56)

WAA5=cbind(WAA5_12,WAA5_13,WAA5_14,WAA5_15,WAA5_16,WAA5_23,WAA5_24,WAA5_25,WAA5_26,W
AA5_34,WAA5_35,WAA5_36,WAA5_45,WAA5_46,WAA5_56)

WAA=rbind(WAA1,WAA2,WAA3,WAA4,WAA5)

SNPs_pair<-
c("SNP1_2","SNP1_3","SNP1_4","SNP1_5","SNP1_6","SNP2_3","SNP2_4","SNP2_5","SNP2_6","
SNP3_4","SNP3_5","SNP3_6","SNP4_5","SNP4_6","SNP5_6")

colnames<-SNPs_pair
```

```
WAA<-
matrix(WAA,nrow=length(rownames),ncol=length(colnames),byrow=FALSE,dimnames=list(row
names, colnames))

WAD1=cbind(WAD1_12,WAD1_13,WAD1_14,WAD1_15,WAD1_16,WAD1_23,WAD1_24,WAD1_25,WAD1_26,W
AD1_34,WAD1_35,WAD1_36,WAD1_45,WAD1_46,WAD1_56)

WAD2=cbind(WAD2_12,WAD2_13,WAD2_14,WAD2_15,WAD2_16,WAD2_23,WAD2_24,WAD2_25,WAD2_26,W
AD2_34,WAD2_35,WAD2_36,WAD2_45,WAD2_46,WAD2_56)

WAD3=cbind(WAD3_12,WAD3_13,WAD3_14,WAD3_15,WAD3_16,WAD3_23,WAD3_24,WAD3_25,WAD3_26,W
AD3_34,WAD3_35,WAD3_36,WAD3_45,WAD3_46,WAD3_56)

WAD4=cbind(WAD4_12,WAD4_13,WAD4_14,WAD4_15,WAD4_16,WAD4_23,WAD4_24,WAD4_25,WAD4_26,W
AD4_34,WAD4_35,WAD4_36,WAD4_45,WAD4_46,WAD4_56)

WAD5=cbind(WAD5_12,WAD5_13,WAD5_14,WAD5_15,WAD5_16,WAD5_23,WAD5_24,WAD5_25,WAD5_26,W
AD5_34,WAD5_35,WAD5_36,WAD5_45,WAD5_46,WAD5_56)

WAD=rbind(WAD1,WAD2,WAD3,WAD4,WAD5)

WAD<-
matrix(WAD,nrow=length(rownames),ncol=length(colnames),byrow=FALSE,dimnames=list(row
names, colnames))

WDA1=cbind(WDA1_12,WDA1_13,WDA1_14,WDA1_15,WDA1_16,WDA1_23,WDA1_24,WDA1_25,WDA1_26,W
DA1_34,WDA1_35,WDA1_36,WDA1_45,WDA1_46,WDA1_56)

WDA2=cbind(WDA2_12,WDA2_13,WDA2_14,WDA2_15,WDA2_16,WDA2_23,WDA2_24,WDA2_25,WDA2_26,W
DA2_34,WDA2_35,WDA2_36,WDA2_45,WDA2_46,WDA2_56)

WDA3=cbind(WDA3_12,WDA3_13,WDA3_14,WDA3_15,WDA3_16,WDA3_23,WDA3_24,WDA3_25,WDA3_26,W
DA3_34,WDA3_35,WDA3_36,WDA3_45,WDA3_46,WDA3_56)

WDA4=cbind(WDA4_12,WDA4_13,WDA4_14,WDA4_15,WDA4_16,WDA4_23,WDA4_24,WDA4_25,WDA4_26,W
DA4_34,WDA4_35,WDA4_36,WDA4_45,WDA4_46,WDA4_56)

WDA5=cbind(WDA5_12,WDA5_13,WDA5_14,WDA5_15,WDA5_16,WDA5_23,WDA5_24,WDA5_25,WDA5_26,W
DA5_34,WDA5_35,WDA5_36,WDA5_45,WDA5_46,WDA5_56)

WDA=rbind(WDA1,WDA2,WDA3,WDA4,WDA5)

WDA<-
matrix(WDA,nrow=length(rownames),ncol=length(colnames),byrow=FALSE,dimnames=list(row
names, colnames))

WDD1=cbind(WDD1_12,WDD1_13,WDD1_14,WDD1_15,WDD1_16,WDD1_23,WDD1_24,WDD1_25,WDD1_26,W
DD1_34,WDD1_35,WDD1_36,WDD1_45,WDD1_46,WDD1_56)

WDD2=cbind(WDD2_12,WDD2_13,WDD2_14,WDD2_15,WDD2_16,WDD2_23,WDD2_24,WDD2_25,WDD2_26,W
DD2_34,WDD2_35,WDD2_36,WDD2_45,WDD2_46,WDD2_56)

WDD3=cbind(WDD3_12,WDD3_13,WDD3_14,WDD3_15,WDD3_16,WDD3_23,WDD3_24,WDD3_25,WDD3_26,W
DD3_34,WDD3_35,WDD3_36,WDD3_45,WDD3_46,WDD3_56)

WDD4=cbind(WDD4_12,WDD4_13,WDD4_14,WDD4_15,WDD4_16,WDD4_23,WDD4_24,WDD4_25,WDD4_26,W
DD4_34,WDD4_35,WDD4_36,WDD4_45,WDD4_46,WDD4_56)

WDD5=cbind(WDD5_12,WDD5_13,WDD5_14,WDD5_15,WDD5_16,WDD5_23,WDD5_24,WDD5_25,WDD5_26,W
DD5_34,WDD5_35,WDD5_36,WDD5_45,WDD5_46,WDD5_56)

WDD=rbind(WDD1,WDD2,WDD3,WDD4,WDD5)

WDD<-
matrix(WDD,nrow=length(rownames),ncol=length(colnames),byrow=FALSE,dimnames=list(row
names, colnames))
```

```
# Print WAA, WAD, WDA and WDD matrices
print("WAA")
print(WAA)
print("WAD")
print(WAD)
print("WDA")
print(WDA)
print("WDD")
print(WDD)

# Haplotype frequencies
P1=0.4
P2=0.3
P3=0.2
P4=0.1

# Haplotype additive coding
WAH11=2*P2
WAH12=2*P3
WAH13=2*P4
WAH1_1=cbind(WAH11,WAH12,WAH13)

WAH21=-2*(1-P2)
WAH22=2*P3
WAH23=2*P4
WAH2_2=cbind(WAH21,WAH22,WAH23)

WAH31=2*P2
WAH32=-2*(1-P3)
WAH33=2*P4
WAH3_3=cbind(WAH31,WAH32,WAH33)

WAH41=2*P2
WAH42=2*P3
WAH43=-2*(1-P4)
```

```
WAH4_4=cbind(WAH41,WAH42,WAH43)

WAH51=-(1-2*P2)

WAH52=2*P3

WAH53=2*P4

WAH1_2=cbind(WAH51,WAH52,WAH53)

WAH=rbind(WAH1_1,WAH2_2,WAH3_3,WAH4_4,WAH1_2)

Haps_name<-c("HAP1","HAP2","HAP3")

colnames<-Haps_name

WAH<-
matrix(WAH,nrow=length(rownames),ncol=length(colnames),byrow=FALSE,dimnames=list(row
names, colnames))

print("WAH matrix")

print(WAH)

# Making S matrices

colnames<-ID

WWA=WA%*%t(WA)

KA=mean(diag(WWA))

SA<-WWA/KA

SA<-
matrix(SA,nrow=length(rownames),ncol=length(colnames),byrow=FALSE,dimnames=list(rown
ames, colnames))

print("SA matrix")

print(SA)

WWD=WD%*%t(WD)

KD=mean(diag(WWD))

SD<-WWD/KD

SD<-
matrix(SD,nrow=length(rownames),ncol=length(colnames),byrow=FALSE,dimnames=list(rown
ames, colnames))

print("SD matrix")

print(SD)

WWAA=WAA%*%t(WAA)

KAA=mean(diag(WWAA))

SAA<-WWAA/KAA
```

```
SAA<-
matrix(SAA,nrow=length(rownames),ncol=length(colnames),byrow=FALSE,dimnames=list(row
names, colnames))

print("SAA matrix")

print(SAA)

WWAD=WAD%*%t(WAD)

KAD=mean(diag(WWAD))

WWDA=WDA%*%t(WDA)

KDA=mean(diag(WWDA))

WWAD2=WWAD+WWDA

KAD2=mean(diag(WWAD2))

SAD2<-WWAD2/KAD2

SAD2<-
matrix(SAD2,nrow=length(rownames),ncol=length(colnames),byrow=FALSE,dimnames=list(ro
wnames, colnames))

print("SAD2 matrix")

print(SAD2)

WWDD=WDD%*%t(WDD)

KDD=mean(diag(WWDD))

SDD<-WWDD/KDD

SDD<-
matrix(SDD,nrow=length(rownames),ncol=length(colnames),byrow=FALSE,dimnames=list(row
names, colnames))

print("SDD matrix")

print(SDD)

WAHWAHt=WAH%*%t(WAH)

KAH=mean(diag(WAHWAHt))

SAH=WAHWAHt/KAH

SAH<-
matrix(SAH,nrow=length(rownames),ncol=length(colnames),byrow=FALSE,dimnames=list(row
names, colnames))

print("SAH matrix")

print(SAH)
```

```
# 2. GREML estimation of variance components and heritability

ID<-factor(ID)
FIX<-factor(FIX)
data<-data.frame(ID,y)
Z<-model.matrix(y ~ID-1)
Z<-Z[-5,]
Zt<-t(Z)
X<-model.matrix(y~FIX-1)
X<-X[-5,]
y<-y[-5]

#Iteration
ZAZ=Z%*%SA%*%(Zt)
ZDZ=Z%*%SD%*%(Zt)
ZAAZ=Z%*%SAA%*%(Zt)
ZAD2Z=Z%*%SAD2%*%(Zt)
ZDDZ=Z%*%SDD%*%(Zt)
ZAHZ=Z%*%SAH%*%(Zt)

var_a=3
var_d=1
var_aa=6
var_ad2=4
var_dd=2
var_ah=3
var_e=1
zrows<-nrow(Z)
k<-1
for(i in (1:k)){
      vge<-diag(var_e,zrows,zrows)
      V<-var_a*ZAZ+var_d*ZDZ+var_aa*ZAAZ+var_ad2*ZAD2Z+var_dd*ZDDZ+var_ah*ZAHZ+vge
      IV<-ginv(V)
      XIV<-t(X)%*%IV
      XIVX<-XIV%*%X
```

```
      XIVY<-XIV%*%y
      P<-IV-(t(XIV)%*%ginv(XIVX)%*%XIV)
# Get new var_a,var_d,var_aa,var_ad2,var_dd,var_ah,var_e
      PZAZ=P%*%ZAZ
      PZDZ=P%*%ZDZ
      PZAAZ=P%*%ZAAZ
      PZAD2Z=P%*%ZAD2Z
      PZDDZ=P%*%ZDDZ
      PZAHZ=P%*%ZAHZ
      var_a=(t(y)%*%PZAZ%*%P%*%y*(var_a/sum(diag(PZAZ))))[1]
      var_d=(t(y)%*%PZDZ%*%P%*%y*(var_d/sum(diag(PZDZ))))[1]
      var_aa=(t(y)%*%PZAAZ%*%P%*%y*(var_aa/sum(diag(PZAAZ))))[1]
      var_ad2=(t(y)%*%PZAD2Z%*%P%*%y*(var_ad2/sum(diag(PZAD2Z))))[1]
      var_dd=(t(y)%*%PZDDZ%*%P%*%y*(var_dd/sum(diag(PZDDZ))))[1]
      var_ah=(t(y)%*%PZAHZ%*%P%*%y*(var_ah/sum(diag(PZAHZ))))[1]
      var_e=(t(y)%*%P%*%P%*%y*(var_e/sum(diag(P))))[1]
}
rownames<-
c("Variance_component_A","Variance_component_D","Variance_component_AA","Variance_co
mponent_AD2","Variance_component_DD","Variance_component_AH","Variance_component_E")
colnames<-c("Value")
outv<-
matrix(c(var_a,var_d,var_aa,var_ad2,var_dd,var_ah,var_e),nrow=length(rownames),ncol=
length(colnames),byrow=FALSE,dimnames=list(rownames, colnames))
print(outv)
# Get Heritability
var_p<-var_a+var_d+var_aa+var_ad2+var_dd+var_ah+var_e
Heritability_A<-var_a/var_p
Heritability_D<-var_d/var_p
Heritability_AA<-var_aa/var_p
Heritability_AD2<-var_ad2/var_p
Heritability_DD<-var_dd/var_p
Heritability_AH<-var_ah/var_p
Heritability_G<-
Heritability_A+Heritability_D+Heritability_AA+Heritability_AD2+Heritability_DD+Herit
ability_AH
rownames<-
c("Heritability_A","Heritability_D","Heritability_AA","Heritability_AD2","Heritabili
ty_DD","Heritability_AH","Heritability_G")
```

```
colnames<-c("Value")

outhr2<-
matrix(c(Heritability_A,Heritability_D,Heritability_AA,Heritability_AD2,Heritability
_DD,Heritability_AH,Heritability_G),nrow=length(rownames),ncol=length(colnames),byro
w=FALSE,dimnames=list(rownames, colnames))

print(outhr2)

# 3. GBLUP and reliability values

vge<-diag(var_e,zrows,zrows)

GA<-var_a*SA

GD<-var_d*SD

GAA<-var_aa*SAA

GAD2<-var_ad2*SAD2

GDD<-var_dd*SDD

GAH<-var_ah*SAH

ZGAZ=Z%*%GA%*%(Zt)

ZGDZ=Z%*%GD%*%(Zt)

ZGAAZ=Z%*%GAA%*%(Zt)

ZGAD2Z=Z%*%GAD2%*%(Zt)

ZGDDZ=Z%*%GDD%*%(Zt)

ZGAHZ=Z%*%GAH%*%(Zt)

V<-ZGAZ+ZGDZ+ZGAAZ+ZGAD2Z+ZGDDZ+ZGAHZ+vge

IV<-ginv(V)

XIV<-t(X)%*%IV

XIVX<-XIV%*%X

XIVY<-XIV%*%y

P<-IV-(t(XIV)%*%ginv(XIVX)%*%XIV)

PY<-P%*%y

ZPY<-Zt%*%PY

ZtPZ<-Zt%*%P%*%Z

Gblup_A<-GA%*%ZPY

Rel_A<-diag(GA%*%ZtPZ%*%GA)/diag(GA) #Reliability_A

Gblup_D<-GD%*%ZPY

Rel_D<-diag(GD%*%ZtPZ%*%GD)/diag(GD)

Gblup_AA<-GAA%*%ZPY

Rel_AA<-diag(GAA%*%ZtPZ%*%GAA)/diag(GAA)
```

```
Gblup_AD2<-GAD2%*%ZPY

Rel_AD2<-diag(GAD2%*%ZtPZ%*%GAD2)/diag(GAD2)

Gblup_DD<-GDD%*%ZPY

Rel_DD<-diag(GDD%*%ZtPZ%*%GDD)/diag(GDD)

Gblup_AH<-GAH%*%ZPY

Rel_AH<-diag(GAH%*%ZtPZ%*%GAH)/diag(GAH)

Gblup_G<-Gblup_A+Gblup_D+Gblup_AA+Gblup_AD2+Gblup_DD+Gblup_AH

G<-GA+GD+GAA+GAD2+GDD+GAH

Rel_G<-diag(G%*%ZtPZ%*%G)/diag(G)

rownames<-ID

colnames<-
c("Gblup_A","Rel_A","Gblup_D","Rel_D","Gblup_AA","Rel_AA","Gblup_AD2","Rel_AD2","Gbl
up_DD","Rel_DD","Gblup_AH","Rel_AH","Gblup_G","Rel_G")

outebv<-
matrix(c(Gblup_A,Rel_A,Gblup_D,Rel_D,Gblup_AA,Rel_AA,Gblup_AD2,Rel_AD2,Gblup_DD,Rel_
DD,Gblup_AH,Rel_AH,Gblup_G,Rel_G),nrow=length(rownames),ncol=length(colnames),byrow=
FALSE,dimnames=list(rownames, colnames))

print(outebv)

# 4. Effect and hertiability estimates for all levels of all effect types

# A and D effects

TA=t(WA)/sqrt(KA)

TD=t(WD)/sqrt(KD)

TAH=t(WAH)/sqrt(KAH)

rownames<-SNPs_name

colnames<-c("Effect_A","Heritability_A","Effect_D","Heritability_D")

mrk_eff_A=var_a*TA%*%ZPY

Her_eff_A=(mrk_eff_A^2/sum(mrk_eff_A^2))*Heritability_A

mrk_eff_D=var_d*TD%*%ZPY

Her_eff_D=(mrk_eff_D^2/sum(mrk_eff_D^2))*Heritability_D

SNP_A_D<-
matrix(c(mrk_eff_A,Her_eff_A,mrk_eff_D,Her_eff_D),nrow=length(rownames),ncol=length(
colnames),byrow=FALSE,dimnames=list(rownames, colnames))

print(SNP_A_D)

# AH effect

rownames<-Haps_name
```

```
colnames<-c("Effect_AH","Heritability_AH")

mrk_eff_AH=var_ah*TAH%*%ZPY

Her_eff_AH=(mrk_eff_AH^2/sum(mrk_eff_AH^2))*Heritability_AH

SNP_eff_AH<-
matrix(c(mrk_eff_AH,Her_eff_AH),nrow=length(rownames),ncol=length(colnames),byrow=FALSE,dimnames=list(rownames, colnames))

print(SNP_eff_AH)

# AA,AD,DA and DD effects

TAA=t(WAA)/sqrt(KAA)

TAD=t(WAD)/sqrt(KAD)

TDA=t(WDA)/sqrt(KDA)

TDD=t(WDD)/sqrt(KDD)

mrk_eff_AA=var_aa*TAA%*%ZPY

Her_eff_AA=(mrk_eff_AA^2/sum(mrk_eff_AA^2))*Heritability_AA

mrk_eff_AD=var_ad2*TAD%*%ZPY

Her_eff_AD=(mrk_eff_AD^2/sum(mrk_eff_AD^2))*Heritability_AD2

mrk_eff_DA=var_ad2*TDA%*%ZPY

Her_eff_DA=(mrk_eff_DA^2/sum(mrk_eff_DA^2))*Heritability_AD2

mrk_eff_DD=var_dd*TDD%*%ZPY

Her_eff_DD=(mrk_eff_DD^2/sum(mrk_eff_DD^2))*Heritability_DD

Her_eff_epi<-
cbind(mrk_eff_AA,Her_eff_AA,mrk_eff_AD,Her_eff_AD,mrk_eff_DA,Her_eff_DA,mrk_eff_DD,Her_eff_DD)

rownames<-SNPs_pair

colnames<-
c("Effect_AA","Heritability_AA","Effect_AD","Heritability_AD","Effect_DA","Heritability_DA","Effect_DD","Heritability_DD")

SNP_epi_eff<-
matrix(Her_eff_epi,nrow=length(rownames),ncol=length(colnames),byrow=FALSE,dimnames=list(rownames, colnames))

print(SNP_epi_eff)

# 5. Genomic relationship matrices of AxA, AxD and DxD using additive and dominance model matrices

#    Method of Jiang and Reif, 2020

rownames<-ID

colnames<-ID
```

```
WWAA=0.5*((WA%*%t(WA))^2-(WA^2)%*%(t((WA^2))))

KAA=mean(diag(WWAA))

SAA<-WWAA/KAA

SAA<-
matrix(SAA,nrow=length(rownames),ncol=length(colnames),byrow=FALSE,dimnames=list(row
names, colnames))

print("SAA matrix")

print(SAA)

WWAD2=(WA%*%t(WA))*(WD%*%t(WD))-(WA*WD)%*%t(WA*WD)

KAD2=mean(diag(WWAD2))

SAD2<-WWAD2/KAD2

SAD2<-
matrix(SAD2,nrow=length(rownames),ncol=length(colnames),byrow=FALSE,dimnames=list(ro
wnames, colnames))

print("SAD2 matrix")

print(SAD2)

WWDD=0.5*((WD%*%t(WD))^2-(WD^2)%*%(t((WD^2))))

KDD=mean(diag(WWDD))

SDD<-WWDD/KDD

SDD<-
matrix(SDD,nrow=length(rownames),ncol=length(colnames),byrow=FALSE,dimnames=list(row
names, colnames))

print("SDD matrix")

print(SDD)

#***********

# Third order epistasis

# Genomic relationship matrices using SNP model matrices

# Method of Jiang and Reif, 2020.

genostat_update_epi <- function(M=NULL,mtype){

  colnames<-
c("genotype_000","genotype_001","genotype_002","genotype_010","genotype_011","genoty
pe_012","genotype_020","genotype_021","genotype_022","genotype_100","genotype_101","
genotype_102","genotype_110","genotype_111","genotype_112","genotype_120","genotype_
121","genotype_122","genotype_200","genotype_201","genotype_202","genotype_210","gen
otype_211","genotype_212","genotype_220","genotype_221","genotype_222")

  mrkn<-ncol(M)

  Mepi<-matrix(0,choose(mrkn,3),27)

  a<-1
```

```
  for(i in 1:mrkn){
      for(j in 2:mrkn){
            for(k in 3:mrkn){
                  if((k>j)&&(j>i)){
                        if(mtype=="AAA"){
                              first_s<-c(M[1,i],M[2,i],M[3,i])
                              second_s<-c(M[1,j],M[2,j],M[3,j])
                              third_s<-c(M[1,k],M[2,k],M[3,k])
                              rownames<-
c("WAAA_123","WAAA_124","WAAA_125","WAAA_126","WAAA_134","WAAA_135","WAAA_136","WAAA
_145","WAAA_146","WAAA_156","WAAA_234","WAAA_235","WAAA_236","WAAA_245","WAAA_246","
WAAA_256","WAAA_345","WAAA_346","WAAA_356","WAAA_456")
                        }
                        else if(mtype=="DDD"){
                              first_s<-c(M[4,i],M[5,i],M[6,i])
                              second_s<-c(M[4,j],M[5,j],M[6,j])
                              third_s<-c(M[4,k],M[5,k],M[6,k])
                              rownames<-
c("WDDD_123","WDDD_124","WDDD_125","WDDD_126","WDDD_134","WDDD_135","WDDD_136","WDDD
_145","WDDD_146","WDDD_156","WDDD_234","WDDD_235","WDDD_236","WDDD_245","WDDD_246","
WDDD_256","WDDD_345","WDDD_346","WDDD_356","WDDD_456")
                        }
                        else if(mtype=="AAD"){
                              first_s<-c(M[1,i],M[2,i],M[3,i])
                              second_s<-c(M[1,j],M[2,j],M[3,j])
                              third_s<-c(M[4,k],M[5,k],M[6,k])
                              rownames<-
c("WAAD_123","WAAD_124","WAAD_125","WAAD_126","WAAD_134","WAAD_135","WAAD_136","WAAD
_145","WAAD_146","WAAD_156","WAAD_234","WAAD_235","WAAD_236","WAAD_245","WAAD_246","
WAAD_256","WAAD_345","WAAD_346","WAAD_356","WAAD_456")
                        }
                        else if(mtype=="ADA"){
                              first_s<-c(M[1,i],M[2,i],M[3,i])
                              second_s<-c(M[4,j],M[5,j],M[6,j])
                              third_s<-c(M[1,k],M[2,k],M[3,k])
                              rownames<-
c("WADA_123","WADA_124","WADA_125","WADA_126","WADA_134","WADA_135","WADA_136","WADA
_145","WADA_146","WADA_156","WADA_234","WADA_235","WADA_236","WADA_245","WADA_246","
WADA_256","WADA_345","WADA_346","WADA_356","WADA_456")
                        }
```

```
else if(mtype=="DAA"){
	first_s<-c(M[4,i],M[5,i],M[6,i])
	second_s<-c(M[1,j],M[2,j],M[3,j])
	third_s<-c(M[1,k],M[2,k],M[3,k])
	rownames<-
c("WDAA_123","WDAA_124","WDAA_125","WDAA_126","WDAA_134","WDAA_135","WDAA_136","WDAA
_145","WDAA_146","WDAA_156","WDAA_234","WDAA_235","WDAA_236","WDAA_245","WDAA_246","
WDAA_256","WDAA_345","WDAA_346","WDAA_356","WDAA_456")
}
else if(mtype=="DDA"){
	first_s<-c(M[4,i],M[5,i],M[6,i])
	second_s<-c(M[4,j],M[5,j],M[6,j])
	third_s<-c(M[1,k],M[2,k],M[3,k])
	rownames<-
c("WDDA_123","WDDA_124","WDDA_125","WDDA_126","WDDA_134","WDDA_135","WDDA_136","WDDA
_145","WDDA_146","WDDA_156","WDDA_234","WDDA_235","WDDA_236","WDDA_245","WDDA_246","
WDDA_256","WDDA_345","WDDA_346","WDDA_356","WDDA_456")
}
else if(mtype=="DAD"){
	first_s<-c(M[4,i],M[5,i],M[6,i])
	second_s<-c(M[1,j],M[2,j],M[3,j])
	third_s<-c(M[4,k],M[5,k],M[6,k])
	rownames<-
c("WDAD_123","WDAD_124","WDAD_125","WDAD_126","WDAD_134","WDAD_135","WDAD_136","WDAD
_145","WDAD_146","WDAD_156","WDAD_234","WDAD_235","WDAD_236","WDAD_245","WDAD_246","
WDAD_256","WDAD_345","WDAD_346","WDAD_356","WDAD_456")
}
else if(mtype=="ADD"){
	first_s<-c(M[1,i],M[2,i],M[3,i])
	second_s<-c(M[4,j],M[5,j],M[6,j])
	third_s<-c(M[4,k],M[5,k],M[6,k])
	rownames<-
c("WADD_123","WADD_124","WADD_125","WADD_126","WADD_134","WADD_135","WADD_136","WADD
_145","WADD_146","WADD_156","WADD_234","WADD_235","WADD_236","WADD_245","WADD_246","
WADD_256","WADD_345","WADD_346","WADD_356","WADD_456")
}
snp1<-rep(first_s,each=9)
snp2<-rep(second_s,each=3,time=3)
snp3<-rep(third_s,9)
one_comb<-snp1*snp2*snp3
```

```
                        Mepi[a,]<-one_comb

                        a<-a+1

                  }

            }

      }

  }

  Mepi<-
matrix(Mepi,nrow=length(rownames),ncol=length(colnames),byrow=FALSE,dimnames=list(ro
wnames, colnames))

  return (Mepi)

}

geno_transform_epi<-function(G=NULL,Gepi=NULL){

      my_3_order_dic=array(0,dim=c(3,3,3))

      my_3_order_dic[1,1,1]=1

      my_3_order_dic[1,1,2]=2

      my_3_order_dic[1,1,3]=3

      my_3_order_dic[1,2,1]=4

      my_3_order_dic[1,2,2]=5

      my_3_order_dic[1,2,3]=6

      my_3_order_dic[1,3,1]=7

      my_3_order_dic[1,3,2]=8

      my_3_order_dic[1,3,3]=9

      my_3_order_dic[2,1,1]=10

      my_3_order_dic[2,1,2]=11

      my_3_order_dic[2,1,3]=12

      my_3_order_dic[2,2,1]=13

      my_3_order_dic[2,2,2]=14

      my_3_order_dic[2,2,3]=15

      my_3_order_dic[2,3,1]=16

      my_3_order_dic[2,3,2]=17

      my_3_order_dic[2,3,3]=18

      my_3_order_dic[3,1,1]=19

      my_3_order_dic[3,1,2]=20

      my_3_order_dic[3,1,3]=21

      my_3_order_dic[3,2,1]=22

      my_3_order_dic[3,2,2]=23
```

```
	my_3_order_dic[3,2,3]=24
	my_3_order_dic[3,3,1]=25
	my_3_order_dic[3,3,2]=26
	my_3_order_dic[3,3,3]=27
	indv<-nrow(G)
	mrkn<-ncol(G)
	r=choose(mrkn,3)
	rownames<-ID
	colnames<-
c("SNP_123","SNP_124","SNP_125","SNP_126","SNP_134","SNP_135","SNP_136","SNP_145","SNP_146","SNP_156","SNP_234","SNP_235","SNP_236","SNP_245","SNP_246","SNP_256","SNP_345","SNP_346","SNP_356","SNP_456")
	W=matrix(0,indv,r)
	for(m in 1:indv){
		a=1
		for(i in 1:mrkn){
			for(j in 2:mrkn){
				for(k in 3:mrkn){
					if((k>j)&&(j>i)){
						indx1=G[m,i]+1
						indx2=G[m,j]+1
						indx3=G[m,k]+1
						W[m,a]<-
Gepi[a,my_3_order_dic[indx1,indx2,indx3]]
						a<-a+1
					}
				}
			}
		}
	}
	W<-
matrix(W,nrow=length(rownames),ncol=length(colnames),byrow=FALSE,dimnames=list(rownames, colnames))
	return (W)
}
rownames<-SNPs_name
colnames<-ID
```

```
genotype_file<-
matrix(c(0,2,0,1,0,2,0,0,0,1,2,0,1,1,1,2,0,1,0,1,0,0,1,1,1,1,1,1,1,0),nrow=length(ro
wnames),ncol=length(colnames),byrow=FALSE,dimnames=list(rownames, colnames))

geno<-t(genotype_file)

rownames<-
c("Additive_0","Additive_1","Additive_2","Dominance_0","Dominance_1","Dominance_2")

colnames<-SNPs_name

geno_stat<-
matrix(c(WA1_0,WA2_0,WA3_0,WA4_0,WA5_0,WA6_0,WA1_1,WA2_1,WA3_1,WA4_1,WA5_1,WA6_1,WA1
_2,WA2_2,WA3_2,WA4_2,WA5_2,WA6_2,WD1_0,WD2_0,WD3_0,WD4_0,WD5_0,WD6_0,WD1_1,WD2_1,WD3
_1,WD4_1,WD5_1,WD6_1,WD1_2,WD2_2,WD3_2,WD4_2,WD5_2,WD6_2),nrow=length(rownames),ncol
=length(colnames),byrow=TRUE,dimnames=list(rownames, colnames))

Mepi<-genostat_update_epi(geno_stat,"AAA")

WAAA<-geno_transform_epi(geno,Mepi)

Mepi<-genostat_update_epi(geno_stat,"AAD")

WAAD<-geno_transform_epi(geno,Mepi)

Mepi<-genostat_update_epi(geno_stat,"ADA")

WADA<-geno_transform_epi(geno,Mepi)

Mepi<-genostat_update_epi(geno_stat,"DAA")

WDAA<-geno_transform_epi(geno,Mepi)

Mepi<-genostat_update_epi(geno_stat,"ADD")

WADD<-geno_transform_epi(geno,Mepi)

Mepi<-genostat_update_epi(geno_stat,"DAD")

WDAD<-geno_transform_epi(geno,Mepi)

Mepi<-genostat_update_epi(geno_stat,"DDA")

WDDA<-geno_transform_epi(geno,Mepi)

Mepi<-genostat_update_epi(geno_stat,"DDD")

WDDD<-geno_transform_epi(geno,Mepi)

# Print WAAA, WAAD, WADA ,WDAA, WADD, WDAD, WDDA, and WDDD matrices

print("WAAA")

print(WAAA)

print("WAAD")

print(WAAD)

print("WADA")

print(WADA)

print("WDAA")

print(WDAA)

print("WADD")
```

```
print(WADD)
print("WDAD")
print(WDAD)
print("WDDA")
print(WDDA)
print("WDDD")
print(WDDD)
#
rownames<-ID
colnames<-ID
WWAAA=WAAA%*%t(WAAA)
KAAA=mean(diag(WWAAA))
SAAA<-WWAAA/KAAA
SAAA<-
matrix(SAAA,nrow=length(rownames),ncol=length(colnames),byrow=FALSE,dimnames=list(ro
wnames, colnames))
print("SAAA matrix")
print(SAAA)
#
WWAAD=WAAD%*%t(WAAD)
WWADA=WADA%*%t(WADA)
WWDAA=WDAA%*%t(WDAA)
WWAAD3=WWAAD+WWADA+WWDAA
KAAD3=mean(diag(WWAAD3))
SAAD3<-WWAAD3/KAAD3
SAAD3<-
matrix(SAAD3,nrow=length(rownames),ncol=length(colnames),byrow=FALSE,dimnames=list(r
ownames, colnames))
print("SAAD3 matrix")
print(SAAD3)
#
WWADD=WADD%*%t(WADD)
WWDAD=WDAD%*%t(WDAD)
WWDDA=WDDA%*%t(WDDA)
WWADD3=WWADD+WWDAD+WWDDA
KADD3=mean(diag(WWADD3))
SADD3<-WWADD3/KADD3
```

```
SADD3<-
matrix(SADD3,nrow=length(rownames),ncol=length(colnames),byrow=FALSE,dimnames=list(r
ownames, colnames))

print("SADD3 matrix")

print(SADD3)

#

WWDDD=WDDD%*%t(WDDD)

KDDD=mean(diag(WWDDD))

SDDD<-WWDDD/KDDD

SDDD<-
matrix(SDDD,nrow=length(rownames),ncol=length(colnames),byrow=FALSE,dimnames=list(ro
wnames, colnames))

print("SDDD matrix")

print(SDDD)

FIX<-c(2,2,1,1,1)

y<-c(1.753222323,1.617654738,1.71311938,1.738835045,-9999)

ID<-factor(ID)

FIX<-factor(FIX)

data<-data.frame(ID,y)

Z<-model.matrix(y ~ID-1)

Z<-Z[-5,]

Zt<-t(Z)

X<-model.matrix(y~FIX-1)

X<-X[-5,]

y<-y[-5]

#Iteration

ZAZ=Z%*%SA%*%(Zt)

ZDZ=Z%*%SD%*%(Zt)

ZAAZ=Z%*%SAA%*%(Zt)

ZAD2Z=Z%*%SAD2%*%(Zt)

ZDDZ=Z%*%SDD%*%(Zt)

ZAAAZ=Z%*%SAAA%*%(Zt)

ZAAD3Z=Z%*%SAAD3%*%(Zt)

ZADD3Z=Z%*%SADD3%*%(Zt)

ZDDDZ=Z%*%SDDD%*%(Zt)

ZAHZ=Z%*%SAH%*%(Zt)
```

```
var_a=3
var_d=1
var_aa=6
var_ad2=4
var_dd=2
var_aaa=9
var_aad3=7
var_add3=5
var_ddd=3
var_ah=3
var_e=1
zrows<-nrow(Z)

#EM-REML
k<-1
for(i in (1:k)){
      vge<-diag(var_e,zrows,zrows)
      V<-
var_a*ZAZ+var_d*ZDZ+var_aa*ZAAZ+var_ad2*ZAD2Z+var_dd*ZDDZ+var_aaa*ZAAAZ+var_aad3*ZAA
D3Z+var_add3*ZADD3Z+var_ddd*ZDDDZ+var_ah*ZAHZ+vge
      IV<-ginv(V)
      XIV<-t(X)%*%IV
      XIVX<-XIV%*%X
      XIVY<-XIV%*%y
      P<-IV-(t(XIV)%*%ginv(XIVX)%*%XIV)
# Get new var_a,var_d,var_aa,var_ad2,var_dd,var_ah,var_e
      PZAZ=P%*%ZAZ
      PZDZ=P%*%ZDZ
      PZAAZ=P%*%ZAAZ
      PZAD2Z=P%*%ZAD2Z
      PZDDZ=P%*%ZDDZ
      PZAAAZ=P%*%ZAAAZ
      PZAAD3Z=P%*%ZAAD3Z
      PZADD3Z=P%*%ZADD3Z
      PZDDDZ=P%*%ZDDDZ
```

```
	PZAHZ=P%*%ZAHZ
	var_a=(t(y)%*%PZAZ%*%P%*%y*(var_a/sum(diag(PZAZ))))[1]
	var_d=(t(y)%*%PZDZ%*%P%*%y*(var_d/sum(diag(PZDZ))))[1]
	var_aa=(t(y)%*%PZAAZ%*%P%*%y*(var_aa/sum(diag(PZAAZ))))[1]
	var_ad2=(t(y)%*%PZAD2Z%*%P%*%y*(var_ad2/sum(diag(PZAD2Z))))[1]
	var_dd=(t(y)%*%PZDDZ%*%P%*%y*(var_dd/sum(diag(PZDDZ))))[1]
	var_aaa=(t(y)%*%PZAAAZ%*%P%*%y*(var_aaa/sum(diag(PZAAAZ))))[1]
	var_aad3=(t(y)%*%PZAAD3Z%*%P%*%y*(var_aad3/sum(diag(PZAAD3Z))))[1]
	var_add3=(t(y)%*%PZADD3Z%*%P%*%y*(var_add3/sum(diag(PZADD3Z))))[1]
	var_ddd=(t(y)%*%PZDDDZ%*%P%*%y*(var_ddd/sum(diag(PZDDDZ))))[1]
	var_ah=(t(y)%*%PZAHZ%*%P%*%y*(var_ah/sum(diag(PZAHZ))))[1]
	var_e=(t(y)%*%P%*%P%*%y*(var_e/sum(diag(P))))[1]
}
rownames<-
c("Variance_component_A","Variance_component_D","Variance_component_AA","Variance_co
mponent_AD2","Variance_component_DD","Variance_component_AAA","Variance_component_AA
D3","Variance_component_ADD3","Variance_component_DDD","Variance_component_AH","Vari
ance_component_E")
colnames<-c("Value")
outv3<-
matrix(c(var_a,var_d,var_aa,var_ad2,var_dd,var_aaa,var_aad3,var_add3,var_ddd,var_ah,
var_e),nrow=length(rownames),ncol=length(colnames),byrow=FALSE,dimnames=list(rowname
s, colnames))
print(outv3)
# Get Heritability
var_p<-
var_a+var_d+var_aa+var_ad2+var_dd+var_aaa+var_aad3+var_add3+var_ddd+var_ah+var_e
Heritability_A<-var_a/var_p
Heritability_D<-var_d/var_p
Heritability_AA<-var_aa/var_p
Heritability_AD2<-var_ad2/var_p
Heritability_DD<-var_dd/var_p
Heritability_AAA<-var_aaa/var_p
Heritability_AAD3<-var_aad3/var_p
Heritability_ADD3<-var_add3/var_p
Heritability_DDD<-var_ddd/var_p
Heritability_AH<-var_ah/var_p
```

```
Heritability_G<-
Heritability_A+Heritability_D+Heritability_AA+Heritability_AD2+Heritability_DD+Herit
ability_AAA+Heritability_AAD3+Heritability_ADD3+Heritability_DDD+Heritability_AH

rownames<-
c("Heritability_A","Heritability_D","Heritability_AA","Heritability_AD2","Heritabili
ty_DD","Heritability_AAA","Heritability_AAD3","Heritability_ADD3","Heritability_DDD"
,"Heritability_AH","Heritability_G")

colnames<-c("Value")

outher<-
matrix(c(Heritability_A,Heritability_D,Heritability_AA,Heritability_AD2,Heritability
_DD,Heritability_AAA,Heritability_AAD3,Heritability_ADD3,Heritability_DDD,Heritabili
ty_AH,Heritability_G),nrow=length(rownames),ncol=length(colnames),byrow=FALSE,dimnam
es=list(rownames, colnames))

print(outher)

# 3. GBLUP and reliability values

vge<-diag(var_e,zrows,zrows)

GA<-var_a*SA

GD<-var_d*SD

GAA<-var_aa*SAA

GAD2<-var_ad2*SAD2

GDD<-var_dd*SDD

GAAA<-var_aaa*SAAA

GAAD3<-var_aad3*SAAD3

GADD3<-var_add3*SADD3

GDDD<-var_ddd*SDDD

GAH<-var_ah*SAH

ZGAZ=Z%*%GA%*%(Zt)

ZGDZ=Z%*%GD%*%(Zt)

ZGAAZ=Z%*%GAA%*%(Zt)

ZGAD2Z=Z%*%GAD2%*%(Zt)

ZGDDZ=Z%*%GDD%*%(Zt)

ZGAAAZ=Z%*%GAAA%*%(Zt)

ZGAAD3Z=Z%*%GAAD3%*%(Zt)

ZGADD3Z=Z%*%GADD3%*%(Zt)

ZGDDDZ=Z%*%GDDD%*%(Zt)

ZGAHZ=Z%*%GAH%*%(Zt)

V<-ZGAZ+ZGDZ+ZGAAZ+ZGAD2Z+ZGDDZ+ZGAAAZ+ZGAAD3Z+ZGADD3Z+ZGDDDZ+ZGAHZ+vge
```

```
IV<-ginv(V)

XIV<-t(X)%*%IV

XIVX<-XIV%*%X

XIVY<-XIV%*%y

P<-IV-(t(XIV)%*%ginv(XIVX)%*%XIV)

PY<-P%*%y

ZPY<-Zt%*%PY

ZtPZ<-Zt%*%P%*%Z

Gblup_A<-GA%*%ZPY

Rel_A<-diag(GA%*%ZtPZ%*%GA)/diag(GA) #Reliability_A

Gblup_D<-GD%*%ZPY

Rel_D<-diag(GD%*%ZtPZ%*%GD)/diag(GD)

Gblup_AA<-GAA%*%ZPY

Rel_AA<-diag(GAA%*%ZtPZ%*%GAA)/diag(GAA)

Gblup_AD2<-GAD2%*%ZPY

Rel_AD2<-diag(GAD2%*%ZtPZ%*%GAD2)/diag(GAD2)

Gblup_DD<-GDD%*%ZPY

Rel_DD<-diag(GDD%*%ZtPZ%*%GDD)/diag(GDD)

Gblup_AAA<-GAAA%*%ZPY

Rel_AAA<-diag(GAAA%*%ZtPZ%*%GAAA)/diag(GAAA)

Gblup_AAD3<-GAAD3%*%ZPY

Rel_AAD3<-diag(GAAD3%*%ZtPZ%*%GAAD3)/diag(GAAD3)

Gblup_ADD3<-GADD3%*%ZPY

Rel_ADD3<-diag(GADD3%*%ZtPZ%*%GADD3)/diag(GADD3)

Gblup_DDD<-GDDD%*%ZPY

Rel_DDD<-diag(GDDD%*%ZtPZ%*%GDDD)/diag(GDDD)

Gblup_AH<-GAH%*%ZPY

Rel_AH<-diag(GAH%*%ZtPZ%*%GAH)/diag(GAH)

Gblup_G<-
Gblup_A+Gblup_D+Gblup_AA+Gblup_AD2+Gblup_DD+Gblup_AAA+Gblup_AAD3+Gblup_ADD3+Gblup_DD
D+Gblup_AH

G<-GA+GD+GAA+GAD2+GDD+GAAA+GAAD3+GADD3+GDDD+GAH

Rel_G<-diag(G%*%ZtPZ%*%G)/diag(G)

rownames<-ID

colnames<-
c("Gblup_A","Rel_A","Gblup_D","Rel_D","Gblup_AA","Rel_AA","Gblup_AD2","Rel_AD2","Gbl
```

```
up_DD","Rel_DD","Gblup_AAA","Rel_AAA","Gblup_AAD3","Rel_AAD3","Gblup_ADD3","Rel_ADD3
","Gblup_DDD","Rel_DDD","Gblup_AH","Rel_AH","Gblup_G","Rel_G")

outgblup<-
matrix(c(Gblup_A,Rel_A,Gblup_D,Rel_D,Gblup_AA,Rel_AA,Gblup_AD2,Rel_AD2,Gblup_DD,Rel_
DD,Gblup_AAA,Rel_AAA,Gblup_AAD3,Rel_AAD3,Gblup_ADD3,Rel_ADD3,Gblup_DDD,Rel_DDD,Gblup
_AH,Rel_AH,Gblup_G,Rel_G),nrow=length(rownames),ncol=length(colnames),byrow=FALSE,di
mnames=list(rownames, colnames))

print(outgblup)

# 6. Third order epistasis genomic relationship matrices using additive and
dominance model matrices

#    Method of Jiang and Reif, 2020

rownames<-ID

colnames<-ID

WWAAA=(1/6)*((WA%*%t(WA))^3)-
(1/2)*(((WA^2)%*%(t(WA^2)))*(WA%*%t(WA)))+(1/3)*((WA^3)%*%(t(WA^3)))

KAAA=mean(diag(WWAAA))

SAAA<-WWAAA/KAAA

SAAA<-
matrix(SAAA,nrow=length(rownames),ncol=length(colnames),byrow=FALSE,dimnames=list(ro
wnames, colnames))

print("SAAA matrix")

print(SAAA)

WWAAD3=0.5*(((WA%*%t(WA))^2)*(WD%*%t(WD))-((WA^2)%*%(t(WA^2)))*(WD%*%t(WD)))-
((WA*WD)%*%(t(WA*WD)))*(WA%*%t(WA))+((WA*WA*WD)%*%(t(WA*WA*WD)))

KAAD3=mean(diag(WWAAD3))

SAAD3<-WWAAD3/KAAD3

SAAD3<-
matrix(SAAD3,nrow=length(rownames),ncol=length(colnames),byrow=FALSE,dimnames=list(r
ownames, colnames))

print("SAAD3 matrix")

print(SAAD3)

WWADD3=0.5*(((WD%*%t(WD))^2)*(WA%*%t(WA))-((WD^2)%*%(t(WD^2)))*(WA%*%t(WA)))-
((WA*WD)%*%(t(WA*WD)))*(WD%*%t(WD))+((WA*WD*WD)%*%(t(WA*WD*WD)))

KADD3=mean(diag(WWADD3))

SADD3<-WWADD3/KADD3

SADD3<-
matrix(SADD3,nrow=length(rownames),ncol=length(colnames),byrow=FALSE,dimnames=list(r
ownames, colnames))
```

```
print("SADD3 matrix")

print(SADD3)

WWDDD=(1/6)*((WD%*%t(WD))^3)-
(1/2)*(((WD^2)%*%(t(WD^2)))*(WD%*%t(WD)))+(1/3)*((WD^3)%*%(t(WD^3)))

KDDD=mean(diag(WWDDD))

SDDD<-WWDDD/KDDD

SDDD<-
matrix(SDDD,nrow=length(rownames),ncol=length(colnames),byrow=FALSE,dimnames=list(ro
wnames, colnames))

print("SDDD matrix")

print(SDDD)
```

**TABLE S1** | Notations of the quantitative genetics (QG) model, reparameterized and equivalent QG (RE-QG) model, and multifactorial (MF) model.

| | QG model | | RE-QG model | |
|---|---|---|---|---|
| Description | QG notations | MF notations | RE-QG notations | MF notations |
| | Model-I: pairwise epistasis effects without dividing into intra- and inter-chromosome effects | | | |
| Additive effects | $\boldsymbol{\alpha}_o$ | $\boldsymbol{\tau}_{1o}$ | $\boldsymbol{\alpha} = k_\alpha^{1/2}\boldsymbol{\alpha}_o$ | $\boldsymbol{\tau}_1$, $m \times 1$ |
| Model matrix | $\mathbf{W}_\alpha$ | $\mathbf{W}_1$ | $\mathbf{T}_\alpha = \mathbf{W}_\alpha / k_\alpha^{1/2}$ | $\mathbf{T}_1$, $n \times m$ |
| Additive values | $\mathbf{a} = \mathbf{W}_\alpha \boldsymbol{\alpha}_o$ | $\mathbf{u}_1$ | $\mathbf{a} = \mathbf{T}_\alpha \boldsymbol{\alpha}$ | $\mathbf{u}_1$, $n \times 1$ |
| Variance of additive effects | $\sigma^2_{\alpha o}$ | $\sigma^2_{1o}$ | $\sigma^2_\alpha = k_\alpha \sigma^2_{\alpha o}$ | $\sigma^2_1$ |
| Variance of additive values | $G_a^{jj} = \sigma^2_{aj} = \sigma^2_{\alpha o}(\mathbf{W}_\alpha \mathbf{W}_\alpha')^{jj}$ | $G_1^{jj} = \sigma^2_{1o}(\mathbf{W}_1\mathbf{W}_1')^{jj}$ | $G_a^{jj} = \sigma^2_{aj} = \sigma^2_\alpha(\mathbf{S}_a)^{jj}$ | $G_1^{jj} = \sigma^2_1(\mathbf{S}_1)^{jj}$ |
| k value | – | – | $k_\alpha = tr(\mathbf{W}_\alpha \mathbf{W}_\alpha')/n$ | $k_1$ |
| Genomic relationship matrix | – | – | $\mathbf{S}_\alpha = \mathbf{T}_\alpha \mathbf{T}_\alpha'$ | $\mathbf{S}_1$, $n \times n$ |
| **G** matrix | $\mathbf{G}_a = \sigma^2_{\alpha o} \mathbf{W}_\alpha \mathbf{W}_\alpha'$ | $\mathbf{G}_1$ | $\mathbf{G}_a = \sigma^2_\alpha \mathbf{S}_\alpha$ | $\mathbf{G}_1$, $n \times n$ |
| Dominance effects | $\boldsymbol{\delta}_o$ | $\boldsymbol{\tau}_{2o}$ | $\boldsymbol{\delta} = k_\delta^{1/2}\boldsymbol{\delta}_o$ | $\boldsymbol{\tau}_2$, $m \times 1$ |
| Model matrix | $\mathbf{W}_\delta$ | $\mathbf{W}_2$ | $\mathbf{T}_\delta = \mathbf{W}_\delta / k_\delta^{1/2}$ | $\mathbf{T}_2$, $n \times m$ |
| Dominance values | $\mathbf{d} = \mathbf{W}_\delta \boldsymbol{\delta}_o$ | $\mathbf{u}_2$ | $\mathbf{d} = \mathbf{T}_\delta \boldsymbol{\delta}$ | $\mathbf{u}_2$, $n \times 1$ |
| Variance of dominance effects | $\sigma^2_{\alpha o}$ | $\sigma^2_{2o}$ | $\sigma^2_\delta = k_\delta \sigma^2_{\delta o}$ | $\sigma^2_2$ |
| Variance of dominance values | $G_d^{jj} = \sigma^2_{dj} = \sigma^2_{\delta o}(\mathbf{W}_\delta \mathbf{W}_\delta')^{jj}$ | $G_2^{jj} = \sigma^2_{2o}(\mathbf{W}_2\mathbf{W}_2')^{jj}$ | $G_d^{jj} = \sigma^2_{dj} = \sigma^2_\delta(\mathbf{S}_\delta)^{jj}$ | $G_2^{jj} = \sigma^2_2(\mathbf{S}_2)^{jj}$ |
| k value | – | – | $k_\delta = tr(\mathbf{W}_\delta \mathbf{W}_\delta')/n$ | $k_2$ |
| Genomic relationship matrix | – | – | $\mathbf{S}_\delta = \mathbf{T}_\delta \mathbf{T}_\delta'$ | $\mathbf{S}_2$, $n \times n$ |
| **G** matrix | $\mathbf{G}_d = \sigma^2_{\delta o} \mathbf{W}_\delta \mathbf{W}_\delta'$ | $\mathbf{G}_2$ | $\mathbf{G}_d = \sigma^2_\delta \mathbf{S}_\delta$ | $\mathbf{G}_2$, $n \times n$ |
| Haplotype additive effects | $\boldsymbol{\alpha}_{oh}$ | $\boldsymbol{\tau}_{3o}$ | $\boldsymbol{\alpha}_h = k_{\alpha h}^{1/2}\boldsymbol{\alpha}_{oh}$ | $\boldsymbol{\tau}_3$, $n_{\alpha h} \times 1$ |

| | | | | |
|---|---|---|---|---|
| Model matrix | $\mathbf{W}_{\alpha h}$ | $\mathbf{W}_3$ | $\mathbf{T}_{\alpha h} = \mathbf{W}_{\alpha h} / k_{\alpha h}^{1/2}$ | $\mathbf{T}_3$, $n\times n_{\alpha h}$ |
| Haplotype additive values | $\mathbf{a}_h = \mathbf{W}_{\alpha h}\boldsymbol{\alpha}_{oh}$ | $\mathbf{u}_3$ | $\mathbf{a}_h = \mathbf{T}_{\alpha h}\boldsymbol{\alpha}_h$ | $\mathbf{u}_3$, $n\times 1$ |
| Variance of haplotype additive effects | $\sigma^2_{\alpha ho}$ | $\sigma^2_{3o}$ | $\sigma^2_{\alpha h} = k_{\alpha h}\sigma^2_{ho}$ | $\sigma^2_3$ |
| Variance of haplotype additive values | $G^{ij}_{ah} = \sigma^2_{ahj} = \sigma^2_{\alpha ho}(\mathbf{W}_{\alpha h}\mathbf{W}_{\alpha h}')^{ij}$ | $G^{ij}_3 = \sigma^2_{3o}(\mathbf{W}_3\mathbf{W}_3')^{ij}$ | $G^{ij}_{ah} = \sigma^2_{ahj} = \sigma^2_{\alpha h}(\mathbf{S}_{\alpha h})^{ij}$ | $G^{ij}_3 = \sigma^2_3(\mathbf{S}_3)^{ij}$ |
| k value | – | – | $k_{\alpha h} = tr(\mathbf{W}_{\alpha h}\mathbf{W}_{\alpha h}')/n$ | $k_3$ |
| Genomic relationship matrix | – | – | $\mathbf{S}_{\alpha h} = \mathbf{T}_{\alpha h}\mathbf{T}_{\alpha h}'$ | $\mathbf{S}_3$, $n\times n$ |
| **G** matrix | $\mathbf{G}_{ah} = \sigma^2_{\alpha ho}\mathbf{W}_{\alpha h}\mathbf{W}_{\alpha h}'$ | $\mathbf{G}_3$ | $\mathbf{G}_{ah} = \sigma^2_{\alpha h}\mathbf{S}_{\alpha h}$ | $\mathbf{G}_3$, $n\times n$ |
| A×A effects | $(\boldsymbol{\alpha\alpha})_o$ | $\boldsymbol{\tau}_{4o}$ | $(\boldsymbol{\alpha\alpha}) = k^{1/2}_{\alpha\alpha}(\boldsymbol{\alpha\alpha})_o$ | $\boldsymbol{\tau}_4$, $\binom{m}{2}\times 1$ |
| Model matrix | $\mathbf{W}_{\alpha\alpha}$ | $\mathbf{W}_4$ | $\mathbf{T}_{\alpha\alpha} = \mathbf{W}_{\alpha\alpha} / k^{1/2}_{\alpha\alpha}$ | $\mathbf{T}_4$, $n\times\binom{m}{2}$ |
| A×A values | $\mathbf{aa} = \mathbf{W}_{\alpha\alpha}(\boldsymbol{\alpha\alpha})_o$ | $\mathbf{u}_4$ | $\mathbf{aa} = \mathbf{T}_{\alpha\alpha}(\boldsymbol{\alpha\alpha})$ | $\mathbf{u}_4$, $n\times 1$ |
| Variance of A×A effects | $\sigma^2_{\alpha\alpha o}$ | $\sigma^2_{4o}$ | $\sigma^2_{\alpha\alpha} = k_{\alpha\alpha}\sigma^2_{\alpha\alpha o}$ | $\sigma^2_4$ |
| Variance of A×A values | $G^{ij}_{aa} = \sigma^2_{aaj} = \sigma^2_{\alpha\alpha o}(\mathbf{W}_{\alpha\alpha}\mathbf{W}_{\alpha\alpha}')^{ij}$ | $G^{ij}_4 = \sigma^2_4(\mathbf{W}_4\mathbf{W}_4')^{ij}$ | $G^{ij}_{aa} = \sigma^2_{aaj} = \sigma^2_{\alpha\alpha}(\mathbf{S}_{\alpha\alpha})^{ij}$ | $G^{ij}_4 = \sigma^2_4(\mathbf{S}_4)^{ij}$ |
| k value | – | – | $k_{\alpha\alpha} = tr(\mathbf{W}_{\alpha\alpha}\mathbf{W}_{\alpha\alpha}')/n$ | $k_4$ |
| Genomic relationship matrix | – | – | $\mathbf{S}_{\alpha\alpha} = \mathbf{T}_{\alpha\alpha}\mathbf{T}_{\alpha\alpha}'$, | $\mathbf{S}_4$, $n\times n$ |
| **G** matrix | $\mathbf{G}_{aa} = \sigma^2_{\alpha\alpha o}\mathbf{W}_{\alpha\alpha}\mathbf{W}_{\alpha\alpha}'$ | $\mathbf{G}_4$ | $\mathbf{G}_{aa} = \sigma^2_{\alpha\alpha}\mathbf{S}_{\alpha\alpha}$, | $\mathbf{G}_4$, $n\times n$ |
| A×D and D×A effects | $(\boldsymbol{\alpha\delta})^{(2)}_o$, | $\boldsymbol{\tau}_{5o}$ | $(\boldsymbol{\delta\alpha}) = k^{1/2}_{\delta\alpha}(\boldsymbol{\alpha\delta})^{(2)}_o$ | $\boldsymbol{\tau}_5$ $2\binom{m}{2}\times 1$ |
| Model matrix | $\mathbf{W}^{(2)}_{\alpha\delta}$, | $\mathbf{W}_5$ | $\mathbf{T}^{(2)}_{\alpha\delta} = \mathbf{W}^{(2)}_{\alpha\delta}/k^{1/2}_{\alpha\delta}$ | $\mathbf{T}_5$, $n\times 2\binom{m}{2}$ |
| A×D and D×A values | $\mathbf{ad}^{(2)} = \mathbf{W}^{(2)}_{\alpha\delta}(\boldsymbol{\alpha\delta})^{(2)}_o$ | $\mathbf{u}_5$ | $\mathbf{ad}^{(2)} = \mathbf{T}^{(2)}_{\alpha\delta}(\boldsymbol{\alpha\delta})^{(2)}$ | $\mathbf{u}_5$, $n\times 1$ |
| Variance of A×D and D×A effects | $\sigma^2_{\alpha\delta o}$ | $\sigma^2_{5o}$ | $\sigma^2_{\alpha\delta} = k_{\alpha\delta}\sigma^2_{\alpha\delta o}$ | $\sigma^2_5$ |

| | | | | |
|---|---|---|---|---|
| Variance of A×D and D×A values | $G^{jj}_{ad} = \sigma^2_{adj} = \sigma^2_{\alpha\delta o}[\mathbf{W}^{(2)}_{\alpha\delta}\mathbf{W}^{(2)\prime}_{\alpha\delta}]^{jj}$ | $G^{jj}_5 = \sigma^2_{5o}[\mathbf{W}^{(2)}_5\mathbf{W}^{(2)\prime}_5]^{jj}$ | $G^{jj}_{ad} = \sigma^2_{adj} = \sigma^2_{\alpha\delta}(\mathbf{S}_{\alpha\delta})^{jj}$ | $G^{jj}_5 = \sigma^2_5(\mathbf{S}_5)^{jj}$ |
| k value | – | – | $k_{\alpha\delta} = tr(\mathbf{W}^{(2)}_{\alpha\delta}\mathbf{W}^{(2)\prime}_{\alpha\delta})/n$ | $k_5$ |
| Genomic relationship matrix | – | – | $\mathbf{S}_{\alpha\delta} = \mathbf{T}^{(2)}_{\alpha\delta}\mathbf{T}^{(2)\prime}_{\alpha\delta}$ | $\mathbf{S}_5$, $n \times n$ |
| **G** matrix | $\mathbf{G}_{ad} = \sigma^2_{\alpha\delta o}\mathbf{W}^{(2)}_{\alpha\delta}\mathbf{W}^{(2)\prime}_{\alpha\delta}$ | $\mathbf{G}_5$ | $\mathbf{G}_{ad} = \sigma^2_{\alpha\delta}\mathbf{S}_{\alpha\delta}$ | $\mathbf{G}_5$, $n \times n$ |
| D×D effects | $(\boldsymbol{\delta\delta})_o$ | $\boldsymbol{\tau}_{6o}$ | $(\boldsymbol{\delta\delta}) = k^{1/2}_{\delta\delta}(\boldsymbol{\delta\delta})_o$ | $\boldsymbol{\tau}_6$, $\binom{m}{2} \times 1$ |
| Model matrix | $\mathbf{W}_{\delta\delta}$ | $\mathbf{W}_6$ | $\mathbf{T}_{\delta\delta} = \mathbf{W}_{\delta\delta}/k^{1/2}_{\delta\delta}$ | $\mathbf{T}_6$, $n \times \binom{m}{2}$ |
| D×D values | $\mathbf{dd} = \mathbf{W}_{\delta\delta}(\boldsymbol{\delta\delta})_o$ | $\mathbf{u}_6$ | $\mathbf{dd} = \mathbf{T}_{\delta\delta}(\boldsymbol{\delta\delta})$ | $\mathbf{u}_6$, $n \times 1$ |
| Variance of D×D effects | $\sigma^2_{\delta\delta o}$ | $\sigma^2_{6o}$ | $\sigma^2_{\delta\delta} = k_{\delta\delta}\sigma^2_{\delta\delta o}$ | $\sigma^2_6$ |
| Variance of D×D values | $G^{jj}_{dd} = \sigma^2_{ddj} = \sigma^2_{\delta\delta o}(\mathbf{W}_{\delta\delta}\mathbf{W}_{\delta\delta}')^{jj}$ | $G^{jj}_6 = \sigma^2_{6o}[\mathbf{W}^{(2)}_6\mathbf{W}^{(2)\prime}_6]^{jj}$ | $G^{jj}_{dd} = \sigma^2_{ddj} = \sigma^2_{\delta\delta}(\mathbf{S}_{\delta\delta})^{jj}$ | $G^{jj}_6 = \sigma^2_6(\mathbf{S}_6)^{jj}$ |
| k value | – | – | $k_{\delta\delta} = \mathbf{W}_{\delta\delta}\mathbf{W}_{\delta\delta}'/n$ | $k_6$ |
| Genomic relationship matrix | – | – | $\mathbf{S}_{\delta\delta} = \mathbf{T}_{\delta\delta}\mathbf{T}_{\delta\delta}'$ | $\mathbf{S}_6$, $n \times n$ |
| **G** matrix | $\mathbf{G}_{\delta\delta} = \sigma^2_{\delta\delta o}\mathbf{W}_{\delta\delta}\mathbf{W}_{\delta\delta}'$ | $\mathbf{G}_6$ | $\mathbf{G}_{\delta\delta} = \sigma^2_{\delta\delta}\mathbf{S}_{\delta\delta}$, | $\mathbf{G}_6$, $n \times n$ |
| A×A×A effects | $(\boldsymbol{\alpha\alpha\alpha})_o$ | $\boldsymbol{\tau}_{7o}$ | $(\boldsymbol{\alpha\alpha\alpha}) = k^{1/2}_{\alpha\alpha\alpha}(\boldsymbol{\alpha\alpha\alpha})_o$ | $\boldsymbol{\tau}_7$, $\binom{m}{3} \times 1$ |
| Model matrix | $\mathbf{W}_{\alpha\alpha\alpha}$ | $\mathbf{W}_7$ | $\mathbf{T}_{\alpha\alpha\alpha} = \mathbf{W}_{\alpha\alpha\alpha}/k^{1/2}_{\alpha\alpha\alpha}$ | $\mathbf{T}_7$, $n \times \binom{m}{3}$ |
| A×A×A values | $\mathbf{aaa} = \mathbf{W}_{\alpha\alpha\alpha}(\boldsymbol{\alpha\alpha\alpha})_o$ | $\mathbf{u}_7$ | $\mathbf{aa} = \mathbf{T}_{\alpha\alpha}(\boldsymbol{\alpha\alpha})$ | $\mathbf{u}_7$, $n \times 1$ |
| Variance of A×A×A effects | $\sigma^2_{\alpha\alpha o}$ | $\sigma^2_{7o}$ | $\sigma^2_{\alpha\alpha\alpha} = k_{\alpha\alpha\alpha}\sigma^2_{\alpha\alpha\alpha o}$ | $\sigma^2_7$ |
| Variance of A×A×A values | $G^{jj}_{aaa} = \sigma^2_{aaaj} = \sigma^2_{\alpha\alpha\alpha o}\mathbf{W}_{\alpha\alpha\alpha}\mathbf{W}_{\alpha\alpha\alpha}'$ | $G^{jj}_7 = \sigma^2_{7o}[\mathbf{W}^{(2)}_7\mathbf{W}^{(2)\prime}_7]^{jj}$ | $G^{jj}_{aaa} = \sigma^2_{aaaj} = \sigma^2_{\alpha\alpha\alpha}(\mathbf{S}_{\alpha\alpha\alpha})^{jj}$ | $G^{jj}_7 = \sigma^2_7(\mathbf{S}_7)^{jj}$ |
| k value | – | – | $k_{\alpha\alpha\alpha} = tr(\mathbf{W}_{\alpha\alpha\alpha}\mathbf{W}_{\alpha\alpha\alpha}')/n$ | $k_7$ |
| Genomic relationship matrix | – | – | $\mathbf{S}_{\alpha\alpha\alpha} = \mathbf{T}_{\alpha\alpha\alpha}\mathbf{T}_{\alpha\alpha\alpha}'$ | $\mathbf{S}_7$, $n \times n$ |
| **G** matrix | $\mathbf{G}_{aaa} = \sigma^2_{\alpha\alpha\alpha o}\mathbf{W}_{\alpha\alpha\alpha}\mathbf{W}_{\alpha\alpha\alpha}'$ | $\mathbf{G}_7$ | $\mathbf{G}_{aaa} = \sigma^2_{\alpha\alpha\alpha}\mathbf{S}_{\alpha\alpha\alpha}$ | $\mathbf{G}_7$, $n \times n$ |
| A×A×D effects | $(\boldsymbol{\alpha\alpha\delta})^{(3)}_o$ | $\boldsymbol{\tau}_{8o}$ | $(\boldsymbol{\alpha\alpha\delta})^{(3)}_o = k^{1/2}_{\alpha\alpha\delta}(\boldsymbol{\alpha\alpha\delta})^{(3)}_o$ | $\boldsymbol{\tau}_8$, $3\binom{m}{3} \times 1$ |

| | | | | |
|---|---|---|---|---|
| Model matrix | $\mathbf{W}^{(3)}_{\alpha\alpha\delta}$ | $\mathbf{W}_{8}$ | $\mathbf{T}^{(3)}_{\alpha\alpha\delta} = \mathbf{W}^{(3)}_{\alpha\alpha\delta}/k^{1/2}_{\alpha\alpha\delta}$ | $\mathbf{T}_{8}$, $n\times[3\binom{m}{3}]$ |
| A×A×D values | $\mathbf{aad}^{(3)} = \mathbf{W}^{(3)}_{\alpha\alpha\delta}(\boldsymbol{\alpha\alpha\delta})^{(3)}_{o}$ | $\mathbf{u}_{8}$ | $\mathbf{aad}^{(3)} = \mathbf{T}^{(3)}_{\alpha\alpha\delta}(\boldsymbol{\alpha\alpha\delta})^{(3)}$ | $\mathbf{u}_{8}$, $n\times 1$ |
| Variance of A×A×D, A×D×A and D×A×A effects | $\sigma^{2}_{\alpha\alpha\delta o}$ | $\sigma^{2}_{8o}$ | $\sigma^{2}_{\alpha\alpha\delta} = \sigma^{2}_{\alpha\alpha\delta o}$ | $\sigma^{2}_{8}$ |
| Variance of A×A×D, A×D×A and D×A×A values | $G^{jj}_{aad} = \sigma^{2}_{aadj} = \sigma^{2}_{\alpha\alpha\delta o}[\mathbf{W}^{(3)}_{\alpha\alpha\delta}\mathbf{W}^{(3)\prime}_{\alpha\alpha\delta}]^{jj}$ | $G^{jj}_{8} = \sigma^{2}_{8o}[\mathbf{W}^{(2)}_{8}\mathbf{W}^{(2)\prime}_{8}]^{jj}$ | $G^{jj}_{aad} = \sigma^{2}_{aadj} = \sigma^{2}_{\alpha\alpha\delta}(\mathbf{S}_{\alpha\alpha\delta})^{jj}$ | $G^{jj}_{8} = \sigma^{2}_{8}(\mathbf{S}_{8})^{jj}$ |
| k value | – | – | $k_{\alpha\alpha\delta} = tr(\mathbf{W}^{(3)}_{\alpha\alpha\delta}\mathbf{W}^{(3)\prime}_{\alpha\alpha\delta})/n$ | $k_{8}$ |
| Genomic relationship matrix | – | – | $\mathbf{S}_{\alpha\alpha\delta} = \mathbf{T}^{(3)}_{\alpha\alpha\delta}\mathbf{T}^{(3)\prime}_{\alpha\alpha\delta}$ | $\mathbf{S}_{8}$, $n\times n$ |
| **G** matrix | $\mathbf{G}_{aad} = \sigma^{2}_{\alpha\alpha\delta o}\mathbf{W}^{(3)}_{\alpha\alpha\delta}\mathbf{W}^{(3)\prime}_{\alpha\alpha\delta}$ | $\mathbf{G}_{8}$ | $\mathbf{G}_{aad} = \sigma^{2}_{\alpha\alpha\delta}\mathbf{S}_{\alpha\alpha\delta}$ | $\mathbf{G}_{8}$, $n\times n$ |
| A×D×D effects | $(\boldsymbol{\alpha\delta\delta})^{(3)}_{o}$ | $\boldsymbol{\tau}_{9o}$ | $(\boldsymbol{\alpha\delta\delta})^{(3)} = k^{1/2}_{\alpha\delta\delta}(\boldsymbol{\alpha\delta\delta})^{(3)}_{o}$ | $\boldsymbol{\tau}_{9}$, $3\binom{m}{3}\times 1$ |
| Model matrix | $\mathbf{W}^{(3)}_{\alpha\delta\delta}$ | $\mathbf{W}_{9}$ | $\mathbf{T}^{(3)}_{\alpha\delta\delta} = \mathbf{W}^{(3)}_{\alpha\delta\delta}/k^{1/2}_{\alpha\delta\delta}$ | $\mathbf{T}_{9}$, $n\times[3\binom{m}{3}]$ |
| A×D×D values | $\mathbf{add}^{(3)} = \mathbf{W}^{(3)}_{\alpha\delta\delta}(\boldsymbol{\alpha\delta\delta})^{(3)}_{o}$ | $\mathbf{u}_{9}$ | $\mathbf{add}^{(3)} = \mathbf{T}^{(3)}_{\alpha\delta\delta}(\boldsymbol{\alpha\delta\delta})^{(3)}$ | $\mathbf{u}_{9}$, $n\times 1$ |
| Variance of A×D×D, D×A×D, and D×D×A effects | $\sigma^{2}_{\alpha\alpha\delta o}$ | $\sigma^{2}_{9o}$ | $\sigma^{2}_{\alpha\alpha\delta} = \sigma^{2}_{\alpha\alpha\delta o}$ | $\sigma^{2}_{9}$ |
| Variance of A×D×D, D×A×D, and D×D×A values | $G^{jj}_{add} = \sigma^{2}_{addj} = \sigma^{2}_{\alpha\delta\delta o}[\mathbf{W}^{(3)}_{\alpha\delta\delta}\mathbf{W}^{(3)\prime}_{\alpha\delta\delta}]^{jj}$ | $G^{jj}_{9} = \sigma^{2}_{9o}[\mathbf{W}^{(2)}_{9}\mathbf{W}^{(2)\prime}_{9}]^{jj}$ | $G^{jj}_{add} = \sigma^{2}_{addj} = \sigma^{2}_{\alpha\delta\delta}(\mathbf{S}_{\alpha\delta\delta})^{jj}$ | $G^{jj}_{9} = \sigma^{2}_{9}(\mathbf{S}_{9})^{jj}$ |
| k value | – | – | $k_{\alpha\delta\delta} = tr(\mathbf{W}^{(3)}_{\alpha\delta\delta}\mathbf{W}^{(3)\prime}_{\alpha\delta\delta})/n$ | $k_{9}$ |
| Genomic relationship matrix | – | – | $\mathbf{S}_{\alpha\delta\delta} = \mathbf{T}^{(3)}_{\alpha\delta\delta}\mathbf{T}^{(3)\prime}_{\alpha\delta\delta}$ | $\mathbf{S}_{9}$, $n\times n$ |
| **G** matrix | $\mathbf{G}_{add} = \sigma^{2}_{\alpha\delta\delta o}\mathbf{W}^{(3)}_{\alpha\delta\delta}\mathbf{W}^{(3)\prime}_{\alpha\delta\delta}$ | $\mathbf{G}_{9}$ | $\mathbf{G}_{add} = \sigma^{2}_{\alpha\delta\delta}\mathbf{S}_{\alpha\delta\delta}$ | $\mathbf{G}_{9}$, $n\times n$ |
| D×D×D effects | $(\boldsymbol{\delta\delta\delta})_{o}$ | $\boldsymbol{\tau}_{10o}$ | $(\boldsymbol{\delta\delta\delta}) = k^{1/2}_{\delta\delta\delta}(\boldsymbol{\delta\delta\delta})_{o}$ | $\boldsymbol{\tau}_{10}$, $\binom{m}{3}\times 1$ |
| Model matrix | $\mathbf{W}_{\delta\delta\delta}$ | $\mathbf{W}_{10}$ | $\mathbf{T}_{\delta\delta\delta} = \mathbf{W}_{\delta\delta\delta}/k^{1/2}_{\delta\delta\delta}$ | $\mathbf{T}_{10}$, $n\times\binom{m}{3}$ |

| | | | | |
|---|---|---|---|---|
| D×D×D values | $\mathbf{ddd} = \mathbf{W}_{\delta\delta\delta}(\boldsymbol{\delta\delta\delta})_o$ | $\mathbf{u}_{10}$ | $\mathbf{ddd} = \mathbf{T}_{\delta\delta\delta}(\boldsymbol{\delta\delta\delta})$ | $\mathbf{u}_{10}$, n×1 |
| Variance of D×D×D effects | $\sigma^2_{\delta\delta\delta o}$ | $\sigma^2_{10o}$ | $\sigma^2_{\alpha\alpha\delta} = \sigma^2_{\alpha\alpha\delta o}$ | $\sigma^2_{10}$ |
| Variance of D×D×D values | $G^{jj}_{ddd} = \sigma^2_{dddj} = \sigma^2_{\delta\delta\delta o}[\mathbf{W}_{\delta\delta\delta}\mathbf{W}_{\delta\delta\delta}']^{jj}$ | $G^{jj}_{10} = \sigma^2_{10o}[\mathbf{W}^{(2)}_{10}\mathbf{W}^{(2)\prime}_{10}]^{jj}$ | $G^{jj}_{ddd} = \sigma^2_{dddj} = \sigma^2_{\delta\delta\delta}(\mathbf{S}_{\delta\delta\delta})^{jj}$ | $G^{jj}_{10} = \sigma^2_{10}(\mathbf{S}_{10})^{jj}$ |
| k value | – | – | $k_{\delta\delta\delta} = tr(\mathbf{W}_{\delta\delta\delta}\mathbf{W}_{\delta\delta\delta}')/n$ | $k_{10}$ |
| Genomic relationship matrix | – | – | $\mathbf{S}_{\delta\delta\delta} = \mathbf{T}_{\delta\delta\delta}\mathbf{T}_{\delta\delta\delta}'$ | $\mathbf{S}_{10}$, n×n |
| **G** matrix | $\mathbf{G}_{ddd} = \sigma^2_{\delta\delta\delta o}\mathbf{W}_{\delta\delta\delta}\mathbf{W}_{\delta\delta\delta}'$ | $\mathbf{G}_{10}$ | $\mathbf{G}_{ddd} = \sigma^2_{\delta\delta\delta}\mathbf{S}_{\delta\delta\delta}$ | $\mathbf{G}_{10}$, n×n |
| | Model-II: pairwise epistasis effects dividing into intra- and inter-chromosome effects<br>► Subscripts 1-3 of the MF model are the same for Model-I and Model-II<br>► Subscripts 4-9 of the MF Model-II are intra- and inter-chromosome terms<br>► Subscript J of the QG Model-II indicates intra-chromosome terms<br>► Subscript B of the QG Model-II indicates inter-chromosome terms<br>► Subscripts 7-10 of the MF Model-I for 3rd-order epistasis effects are changed to 10-13 for the MF Model-II | | | |
| Additive effects | $\boldsymbol{\alpha}_o$ | $\boldsymbol{\tau}_{1o}$ | $\boldsymbol{\alpha} = k^{1/2}_{\alpha}\boldsymbol{\alpha}_o$ | $\boldsymbol{\tau}_1$, m×1 |
| Model matrix | $\mathbf{W}_{\alpha}$ | $\mathbf{W}_1$ | $\mathbf{T}_{\alpha} = \mathbf{W}_{\alpha}/k^{1/2}_{\alpha}$ | $\mathbf{T}_1$, n×m |
| Additive values | $\mathbf{a} = \mathbf{W}_{\alpha}\boldsymbol{\alpha}_o$ | $\mathbf{u}_1$ | $\mathbf{a} = \mathbf{T}_{\alpha}\boldsymbol{\alpha}$ | $\mathbf{u}_1$, n×1 |
| Variance of additive effects | $\sigma^2_{\alpha o}$ | $\sigma^2_{1o}$ | $\sigma^2_{\alpha} = k_{\alpha}\sigma^2_{\alpha o}$ | $\sigma^2_1$ |
| Variance of additive values | $G^{jj}_a = \sigma^2_{aj} = \sigma^2_{\alpha o}(\mathbf{W}_{\alpha}\mathbf{W}_{\alpha}')^{jj}$ | $G^{jj}_1 = \sigma^2_{1o}(\mathbf{W}_1\mathbf{W}_1')^{jj}$ | $G^{jj}_a = \sigma^2_{aj} = \sigma^2_{\alpha}(\mathbf{S}_a)^{jj}$ | $G^{jj}_1 = \sigma^2_1(\mathbf{S}_1)^{jj}$ |
| k value | – | – | $k_{\alpha} = tr(\mathbf{W}_{\alpha}\mathbf{W}_{\alpha}')/n$ | $k_1$ |
| Genomic relationship matrix | – | – | $\mathbf{S}_{\alpha} = \mathbf{T}_{\alpha}\mathbf{T}_{\alpha}'$ | $\mathbf{S}_1$, n×n |
| **G** matrix | $\mathbf{G}_a = \sigma^2_{\alpha o}\mathbf{W}_{\alpha}\mathbf{W}_{\alpha}'$ | $\mathbf{G}_1$ | $\mathbf{G}_a = \sigma^2_{\alpha}\mathbf{S}_{\alpha}$ | $\mathbf{G}_1$, n×n |
| Dominance effects | $\boldsymbol{\delta}_o$ | $\boldsymbol{\tau}_{2o}$ | $\boldsymbol{\delta} = k^{1/2}_{\delta}\boldsymbol{\delta}_o$ | $\boldsymbol{\tau}_2$, m×1 |
| Model matrix | $\mathbf{W}_{\delta}$ | $\mathbf{W}_2$ | $\mathbf{T}_{\delta} = \mathbf{W}_{\delta}/k^{1/2}_{\delta}$ | $\mathbf{T}_2$, n×m |
| Dominance values | $\mathbf{d} = \mathbf{W}_{\delta}\boldsymbol{\delta}_o$ | $\mathbf{u}_2$ | $\mathbf{d} = \mathbf{T}_{\delta}\boldsymbol{\delta}$ | $\mathbf{u}_2$, n×1 |

| | | | | |
|---|---|---|---|---|
| Variance of dominance effects | $\sigma^2_{\alpha o}$ | $\sigma^2_{2o}$ | $\sigma^2_{\delta} = k_{\delta}\sigma^2_{\delta o}$ | $\sigma^2_2$ |
| Variance of dominance values | $G^{jj}_d = \sigma^2_{dj} = \sigma^2_{\delta o}(\mathbf{W}_{\delta}\mathbf{W}_{\delta}')^{jj}$ | $G^{jj}_2 = \sigma^2_{2o}(\mathbf{W}_2\mathbf{W}_2')^{jj}$ | $G^{jj}_d = \sigma^2_{dj} = \sigma^2_{\delta}(\mathbf{S}_{\delta})^{jj}$ | $G^{jj}_2 = \sigma^2_2(\mathbf{S}_2)^{jj}$ |
| k value | – | – | $k_{\delta} = tr(\mathbf{W}_{\delta}\mathbf{W}_{\delta}')/n$ | $k_2$ |
| Genomic relationship matrix | – | – | $\mathbf{S}_{\delta} = \mathbf{T}_{\delta}\mathbf{T}_{\delta}'$ | $\mathbf{S}_2$, $n \times n$ |
| **G** matrix | $\mathbf{G}_d = \sigma^2_{\delta o}\mathbf{W}_{\delta}\mathbf{W}_{\delta}'$ | $\mathbf{G}_2$ | $\mathbf{G}_d = \sigma^2_{\delta}\mathbf{S}_{\delta}$ | $\mathbf{G}_2$, $n \times n$ |
| Haplotype additive effects | $\boldsymbol{\alpha}_{oh}$ | $\boldsymbol{\tau}_{3o}$ | $\boldsymbol{\alpha}_h = k^{1/2}_{\alpha h}\boldsymbol{\alpha}_{oh}$ | $\boldsymbol{\tau}_3$, $n_{\alpha h} \times 1$ |
| Model matrix | $\mathbf{W}_{\alpha h}$ | $\mathbf{W}_3$ | $\mathbf{T}_{\alpha h} = \mathbf{W}_{\alpha h} / k^{1/2}_{\alpha h}$ | $\mathbf{T}_3$, $n \times n_{\alpha h}$ |
| Haplotype additive values | $\mathbf{a}_h = \mathbf{W}_{\alpha h}\boldsymbol{\alpha}_{oh}$ | $\mathbf{u}_3$ | $\mathbf{a}_h = \mathbf{T}_{\alpha h}\boldsymbol{\alpha}_h$ | $\mathbf{u}_3$, $n \times 1$ |
| Variance of haplotype additive effects | $\sigma^2_{\alpha ho}$ | $\sigma^2_{3o}$ | $\sigma^2_{\alpha h} = k_{\alpha h}\sigma^2_{ho}$ | $\sigma^2_3$ |
| Variance of haplotype additive values | $G^{jj}_{ah} = \sigma^2_{ahj} = \sigma^2_{\alpha ho}(\mathbf{W}_{\alpha h}\mathbf{W}_{\alpha h}')^{jj}$ | $G^{jj}_3 = \sigma^2_{3o}(\mathbf{W}_3\mathbf{W}_3')^{jj}$ | $G^{jj}_{ah} = \sigma^2_{ahj} = \sigma^2_{\alpha h}(\mathbf{S}_{\alpha h})^{jj}$ | $G^{jj}_3 = \sigma^2_3(\mathbf{S}_3)^{jj}$ |
| k value | – | – | $k_{\alpha h} = tr(\mathbf{W}_{\alpha h}\mathbf{W}_{\alpha h}')/n$ | $k_3$ |
| Genomic relationship matrix | – | – | $\mathbf{S}_{\alpha h} = \mathbf{T}_{\alpha h}\mathbf{T}_{\alpha h}'$ | $\mathbf{S}_3$, $n \times n$ |
| **G** matrix | $\mathbf{G}_{ah} = \sigma^2_{\alpha ho}\mathbf{W}_{\alpha h}\mathbf{W}_{\alpha h}'$ | $\mathbf{G}_3$ | $\mathbf{G}_{ah} = \sigma^2_{\alpha h}\mathbf{S}_{\alpha h}$ | $\mathbf{G}_3$, $n \times n$ |
| A×A effects | $(\boldsymbol{\alpha\alpha})^{intra}_o$ | $\boldsymbol{\tau}_{4o}$ | $(\boldsymbol{\alpha\alpha})^{intra} = (k^{intra}_{\alpha\alpha})^{1/2}(\boldsymbol{\alpha\alpha})^{intra}_o$ | $\boldsymbol{\tau}_4$, $c_1 \times 1$ |
| Model matrix | $\mathbf{W}^{intra}_{\alpha\alpha}$ | $\mathbf{W}_4$ | $\mathbf{T}^{intra}_{\alpha\alpha} = \mathbf{W}^{intra}_{\alpha\alpha} / (k^{intra}_{\alpha\alpha})^{1/2}$ | $\mathbf{T}_4$, $n \times c_1$ |
| A×A values | $(\mathbf{aa})^{intra} = \mathbf{W}^{intra}_{\alpha\alpha}(\boldsymbol{\alpha\alpha})^{intra}_o$ | $\mathbf{u}_4$ | $(\mathbf{aa})^{intra} = \mathbf{T}^{intra}_{\alpha\alpha}(\boldsymbol{\alpha\alpha})^{intra}$ | $\mathbf{u}_4$, $n \times 1$ |
| Variance of A×A effects | $(\sigma^2_{\alpha\alpha o})^{intra}$ | $\sigma^2_{4o}$ | $(\sigma^2_{\alpha\alpha})^{intra} = (k^{intra}_{\alpha\alpha})^{1/2}(\sigma^2_{\alpha\alpha o})^{intra}$ | $\sigma^2_4$ |
| Variance of A×A values | $(G^{intra}_{aa})^{jj} = (\sigma^2_{\alpha\alpha o})^{intra}[\mathbf{W}^{intra}_{\alpha\alpha}(\mathbf{W}^{intra}_{\alpha\alpha})']^{jj}$ | $G^{jj}_4 = \sigma^2_4(\mathbf{W}_4\mathbf{W}_4')^{jj}$ | $(G^{intra}_{aa})^{jj} = (\sigma^2_{\alpha\alpha})^{intra}(\mathbf{S}^{intra}_{\alpha\alpha})^{jj}$ | $(G_4)^{jj} = \sigma^2_4(\mathbf{S}_4)^{jj}$ |
| k value | – | – | $k^{intra}_{\alpha\alpha} = tr(\mathbf{W}^{intra}_{\alpha\alpha}\mathbf{W}^{intra\prime}_{\alpha\alpha})/n$ | $k_4$ |
| Genomic relationship matrix | – | – | $\mathbf{S}^{intra}_{\alpha\alpha} = \mathbf{T}^{intra}_{\alpha\alpha}\mathbf{T}^{intra\prime}_{\alpha\alpha}$ | $\mathbf{S}_4$, $n \times n$ |

| | | | | |
|---|---|---|---|---|
| **G** matrix | $\mathbf{G}_{aa}^{intra}=(\sigma^2_{\alpha\alpha o})^{intra}\mathbf{W}_{\alpha\alpha}^{intra}(\mathbf{W}_{\alpha\alpha}^{intra})'$ | $\mathbf{G}_4$ | $\mathbf{G}_{aa}^{intra}=(\sigma^2_{\alpha\alpha})^{intra}\mathbf{S}_{\alpha\alpha}^{intra}$ | $\mathbf{G}_4$, n×n |
| A×D and D×A effects | $(\boldsymbol{\alpha\delta})_o^{(2)intra}$ | $\boldsymbol{\tau}_{5o}$ | $(\boldsymbol{\alpha\delta})^{(2)intra}=(k_{\alpha\delta}^{intra})^{1/2}(\boldsymbol{\alpha\delta})_o^{(2)intra}$ | $\boldsymbol{\tau}_5$ $2c_1\times 1$ |
| Model matrix | $\mathbf{W}_{\alpha\delta}^{(2)intra}$ | $\mathbf{W}_5$ | $\mathbf{T}_{\alpha\delta}^{(2)intra}=\mathbf{W}_{\alpha\delta}^{(2)intra}/(k_{\alpha\delta}^{intra})^{1/2}$ | $\mathbf{T}_5$, $n\times(2c_1)$ |
| A×A and D×A values | $(\mathbf{ad})^{(2)intra}=\mathbf{W}_{\alpha\delta}^{(2)intra}(\boldsymbol{\alpha\delta})_o^{intra}$ | $\mathbf{u}_5$ | $(\mathbf{ad})^{(2)intra}=\mathbf{T}_{\alpha\delta}^{(2)intra}(\boldsymbol{\alpha\delta})^{(2)intra}$ | $\mathbf{u}_5$, n×1 |
| Variance of A×D and D×A effects | $(\sigma^2_{\alpha\alpha o})^{intra}$ | $\sigma^2_{5o}$ | $(\sigma^2_{\alpha\delta})^{intra}=(k_{\alpha\delta}^{intra})^{1/2}(\sigma^2_{\alpha\delta o})^{intra}$ | $\sigma^2_5$ |
| Variance of A×D and D×A values | $(G_{ad}^{intra})^{jj}=(\sigma^2_{\alpha\delta o})^{intra}[\mathbf{W}_{\alpha\delta}^{(2)intra}\mathbf{W}_{\alpha\delta}^{(2)intra}{}']^{jj}$ | $G_5^{jj}=\sigma^2_{5o}[\mathbf{W}_5^{(2)}\mathbf{W}_5^{(2)}{}']^{jj}$ | $(G_{ad}^{intra})^{jj}=(\sigma^2_{\alpha\delta})^{intra}(\mathbf{S}_{\alpha\delta}^{intra})^{jj}$ | $(G_5)^{jj}=\sigma^2_5(\mathbf{S}_5)^{jj}$ |
| k value | – | – | $k_{\alpha\delta}^{intra}=tr[\mathbf{W}_{\alpha\delta}^{(2)intra}\mathbf{W}_{\alpha\delta}^{(2)intra}{}']/n$ | $k_5$ |
| Genomic relationship matrix | – | – | $\mathbf{S}_{\alpha\delta}^{intra}=\mathbf{T}_{\alpha\delta}^{(2)intra}\mathbf{T}_{\alpha\delta}^{(2)intra}{}'$ | $\mathbf{S}_5$, n×n |
| **G** matrix | $\mathbf{G}_{ad}^{intra}=(\sigma^2_{\alpha\delta o})^{intra}\mathbf{W}_{\alpha\delta}^{(2)intra}\mathbf{W}_{\alpha\delta}^{(2)intra}{}'$ | $\mathbf{G}_5$ | $\mathbf{G}_{ad}^{intra}=(\sigma^2_{\alpha\delta})^{intra}\mathbf{S}_{\alpha\delta}^{intra}$ | $\mathbf{G}_5$, n×n |
| D×D effects | $(\boldsymbol{\delta\delta})_o^{intra}$ | $\boldsymbol{\tau}_{6o}$ | $(\boldsymbol{\delta\delta})^{intra}=(k_{\delta\delta}^{intra})^{1/2}(\boldsymbol{\delta\delta})_o^{intra}$ | $\boldsymbol{\tau}_6$, $c_1\times 1$ |
| Model matrix | $\mathbf{W}_{\delta\delta}^{intra}$, | $\mathbf{W}_6$ | $\mathbf{T}_{\delta\delta}^{intra}=\mathbf{W}_{\delta\delta}^{intra}/(k_{\delta\delta}^{intra})^{1/2}$ | $\mathbf{T}_6$, $n\times c_1$ |
| D×D values | $(\mathbf{dd})^{intra}=\mathbf{W}_{\delta\delta}^{intra}(\boldsymbol{\delta\delta})_o^{intra}$ | $\mathbf{u}_6$ | $(\mathbf{dd})^{intra}=\mathbf{T}_{\delta\delta}^{intra}(\boldsymbol{\delta\delta})^{intra}$ | $\mathbf{u}_6$, n×1 |
| Variance of D×D effects | $(\sigma^2_{\delta\delta o})^{intra}$ | $\sigma^2_{6o}$ | $(\sigma^2_{\delta\delta})^{intra}=(k_{\delta\delta}^{1/2})^{intra}(\sigma^2_{\delta\delta o})^{intra}$ | $\sigma^2_6$ |
| Variance of D×D values | $(G_{dd}^{intra})^{jj}=(\sigma^2_{\delta\delta o})^{intra}[\mathbf{W}_{\delta\delta}^{intra}(\mathbf{W}_{\delta\delta}^{intra})']^{jj}$ | $G_6^{jj}=\sigma^2_{6o}[\mathbf{W}_6^{(2)}\mathbf{W}_6^{(2)}{}']^{jj}$ | $(G_{dd}^{intra})^{jj}=(\sigma^2_{\delta\delta})^{intra}(\mathbf{S}_{\delta\delta}^{intra})^{jj}$ | $(G_6)^{jj}=\sigma^2_6(\mathbf{S}_6)^{jj}$ |
| k value | – | – | $k_{\delta\delta}^{intra}=tr(\mathbf{W}_{\delta\delta}^{intra}\mathbf{W}_{\delta\delta}^{intra}{}')/n$ | $k_6$ |
| Genomic relationship matrix | – | – | $\mathbf{S}_{\delta\delta}^{intra}=\mathbf{T}_{\delta\delta}^{intra}\mathbf{T}_{\delta\delta}^{intra}{}'$ | $\mathbf{S}_6$, n×n |
| **G** matrix | $\mathbf{G}_{dd}^{intra}=(\sigma^2_{\delta\delta o})^{intra}\mathbf{W}_{\delta\delta}^{intra}(\mathbf{W}_{\delta\delta}^{intra})'$ | $\mathbf{G}_6$ | $\mathbf{G}_{aa}^{intra}=(\sigma^2_{\delta\delta})^{intra}\mathbf{S}_{\delta\delta}^{intra}$ | $\mathbf{G}_6$, n×n |
| | Inter-chromosome pairwise epistasis | | | |
| A×A effects | $(\boldsymbol{\alpha\alpha})_o^{inter}$ | $\boldsymbol{\tau}_{7o}$ | $(\boldsymbol{\alpha\alpha})^{inter}=(k_{\alpha\alpha}^{inter})^{1/2}(\boldsymbol{\alpha\alpha})_o^{inter}$ | $\boldsymbol{\tau}_7$, $c_2\times 1$ |
| Model matrix | $\mathbf{W}_{\alpha\alpha}^{inter}$ | $\mathbf{W}_7$ | $\mathbf{T}_{\alpha\alpha}^{inter}=\mathbf{W}_{\alpha\alpha}^{inter}/(k_{\alpha\alpha}^{inter})^{1/2}$ | $\mathbf{T}_7$, $n\times c_2$ |
| A×A values | $(\mathbf{aa})^{inter}=\mathbf{W}_{\alpha\alpha}^{inter}(\boldsymbol{\alpha\alpha})_o^{inter}$ | $\mathbf{u}_7$ | $(\mathbf{aa})^{inter}=\mathbf{T}_{\alpha\alpha}^{inter}(\boldsymbol{\alpha\alpha})^{inter}$ | $\mathbf{u}_7$, n×1 |

| | | | | |
|---|---|---|---|---|
| Variance of A×A effects | $(\sigma^2_{\alpha\alpha o})^{inter}$ | $\sigma^2_{7o}$ | $(\sigma^2_{\alpha\alpha})^{inter}=(k^{inter}_{\alpha\alpha})^{1/2}(\sigma^2_{\alpha\alpha o})^{inter}$ | $\sigma^2_7$ |
| Variance of A×A values | $(G^{inter}_{aa})^{jj}=(\sigma^2_{\alpha\alpha o})^{inter}[\mathbf{W}^{inter}_{\alpha\alpha}(\mathbf{W}^{inter}_{\alpha\alpha})']^{jj}$ | $G^{jj}_7=\sigma^2_{7o}[\mathbf{W}^{(2)}_7\mathbf{W}^{(2)\prime}_7]^{jj}$ | $(G^{inter}_{aa})^{jj}=(\sigma^2_{\alpha\alpha})^{inter}(\mathbf{S}^{inter}_{\alpha\alpha})^{jj}$ | $(G_7)^{jj}=\sigma^2_7(\mathbf{S}_7)^{jj}$ |
| k value | – | – | $k^{inter}_{\alpha\alpha}=tr(\mathbf{W}^{inter}_{\alpha\alpha}\mathbf{W}^{inter\prime}_{\alpha\alpha})/n$ | $k_7$ |
| Genomic relationship matrix | – | – | $\mathbf{S}^{inter}_{\alpha\alpha}=\mathbf{T}^{inter}_{\alpha\alpha}\mathbf{T}^{inter\prime}_{\alpha\alpha}$ | $\mathbf{S}_7$, n×n |
| **G** matrix | $\mathbf{G}^{inter}_{aa}=(\sigma^2_{\alpha\alpha o})^{inter}\mathbf{W}^{inter}_{\alpha\alpha}(\mathbf{W}^{inter}_{\alpha\alpha})'$ | $\mathbf{G}_7$ | $\mathbf{G}^{inter}_{aa}=(\sigma^2_{\alpha\alpha})^{inter}\mathbf{S}^{inter}_{\alpha\alpha}$ | $\mathbf{G}_7$, n×n |
| A×D and D×A effects | $(\boldsymbol{\alpha\delta})^{(2)inter}_o$ | $\boldsymbol{\tau}_{8o}$ | $(\boldsymbol{\alpha\delta})^{(2)inter}=(k^{inter}_{\alpha\delta})^{1/2}(\boldsymbol{\alpha\delta})^{(2)inter}_o$ | $\boldsymbol{\tau}_8$ $2c_2\times 1$ |
| Model matrix | $\mathbf{W}^{(2)inter}_{\alpha\delta}$ | $\mathbf{W}_8$ | $\mathbf{T}^{(2)inter}_{\alpha\delta}=\mathbf{W}^{(2)inter}_{\alpha\delta}/(k^{inter}_{\alpha\delta})^{1/2}$ | $\mathbf{T}_8$, $n\times(2c_2)$ |
| A×A and D×A values | $(\mathbf{ad})^{(2)inter}=\mathbf{W}^{(2)inter}_{\alpha\delta}(\boldsymbol{\alpha\delta})_o$ | $\mathbf{u}_8$ | $(\mathbf{ad})^{(2)inter}=\mathbf{T}^{(2)inter}_{\alpha\delta}(\boldsymbol{\alpha\delta})^{(2)inter}$ | $\mathbf{u}_8$, n×1 |
| Variance of A×D and D×A effects | $(\sigma^2_{\alpha\alpha o})^{inter}$ | $\sigma^2_{8o}$ | $(\sigma^2_{\alpha\delta})^{inter}=k^{inter}_{\alpha\delta}(\sigma^2_{\alpha\delta o})^{inter}$ | $\sigma^2_8$ |
| Variance of A×D and D×A values | $(G^{inter}_{ad})^{jj}=(\sigma^2_{\alpha\delta o})^{inter}[\mathbf{W}^{(2)inter}_{\alpha\delta}\mathbf{W}^{(2)inter\prime}_{\alpha\delta}]^{jj}$ | $G^{jj}_8=\sigma^2_{8o}[\mathbf{W}^{(2)}_8\mathbf{W}^{(2)\prime}_8]^{jj}$ | $(G^{inter}_{ad})^{jj}=(\sigma^2_{\alpha\delta})^{inter}(\mathbf{S}^{inter}_{\alpha\delta})^{jj}$ | $(G_8)^{jj}=\sigma^2_8(\mathbf{S}_8)^{jj}$ |
| k value | – | – | $k^{inter}_{\alpha\delta}=tr[\mathbf{W}^{(2)inter}_{\alpha\delta}\mathbf{W}^{(2)inter\prime}_{\alpha\delta}]/n$ | $k_8$ |
| Genomic relationship matrix | – | – | $\mathbf{S}^{inter}_{\alpha\delta}=\mathbf{T}^{(2)inter}_{\alpha\delta}\mathbf{T}^{(2)inter\prime}_{\alpha\delta}$ | $\mathbf{S}_8$, n×n |
| **G** matrix | $\mathbf{G}^{inter}_{ad}=(\sigma^2_{\alpha\delta o})^{inter}\mathbf{W}^{(2)inter}_{\alpha\delta}\mathbf{W}^{(2)inter\prime}_{\alpha\delta}$ | $\mathbf{G}_8$ | $\mathbf{G}^{inter}_{ad}=(\sigma^2_{\alpha\delta})^{inter}\mathbf{S}^{inter}_{\alpha\delta}$ | $\mathbf{G}_8$, n×n |
| D×D effects | $(\boldsymbol{\delta\delta})^{inter}_o$ | $\boldsymbol{\tau}_{9o}$ | $(\boldsymbol{\delta\delta})^{inter}=(k^{inter}_{\delta\delta})^{1/2}(\boldsymbol{\delta\delta})^{inter}_o$ | $\boldsymbol{\tau}_9$, $c_2\times 1$ |
| Model matrix | $\mathbf{W}^{inter}_{\delta\delta}$ | $\mathbf{W}_9$ | $\mathbf{T}^{inter}_{\delta\delta}=\mathbf{W}^{inter}_{\delta\delta}/(k^{inter}_{\delta\delta})^{1/2}$ | $\mathbf{T}_9$, $n\times c_2$ |
| D×D values | $(\mathbf{dd})^{inter}=\mathbf{W}^{inter}_{\delta\delta}(\boldsymbol{\delta\delta})^{inter}_o$, | $\mathbf{u}_9$ | $(\mathbf{dd})^{inter}=\mathbf{T}^{inter}_{\delta\delta}(\boldsymbol{\delta\delta})^{inter}$ | $\mathbf{u}_9$, n×1 |
| Variance of D×D effects | $(\sigma^2_{\delta\delta o})^{inter}$ | $\sigma^2_{9o}$ | $(\sigma^2_{\delta\delta})^{inter}=k^{inter}_{\delta\delta}(\sigma^2_{\delta\delta o})^{inter}$ | $\sigma^2_9$ |
| Variance of D×D values | $(G^{inter}_{dd})^{jj}=(\sigma^2_{\delta\delta o})^{inter}[\mathbf{W}^{inter}_{\delta\delta}\mathbf{W}^{inter\prime}_{\delta\delta}]^{jj}$ | $G^{jj}_9=\sigma^2_{9o}[\mathbf{W}^{(2)}_9\mathbf{W}^{(2)\prime}_9]^{jj}$ | $(G^{inter}_{dd})^{jj}=(\sigma^2_{\delta\delta})^{inter}(\mathbf{S}^{inter}_{\delta\delta})^{jj}$ | $(G_9)^{jj}=\sigma^2_9(\mathbf{S}_9)^{jj}$ |
| k value | – | – | $k^{inter}_{\delta\delta}=tr(\mathbf{W}^{inter}_{\delta\delta}\mathbf{W}^{inter\prime}_{\delta\delta})/n$ | $k_9$ |
| Genomic relationship matrix | – | – | $\mathbf{S}^{inter}_{\delta\delta}=\mathbf{T}^{inter}_{\delta\delta}\mathbf{T}^{inter\prime}_{\delta\delta}$ | $\mathbf{S}_9$, n×n |

| | | | | |
|---|---|---|---|---|
| **G** matrix | $\mathbf{G}_{dd}^{inter}=(\sigma^2_{\delta\delta o})^{inter}\mathbf{W}_{\delta\delta}^{inter}\mathbf{W}_{\delta\delta}^{inter}{}'$ | $\mathbf{G}_9$ | $\mathbf{G}_{aa}^{inter}=(\sigma^2_{\delta\delta})^{inter}\mathbf{S}_{\delta\delta}^{inter}$ | $\mathbf{G}_9$, $n\times n$ |
| | Third-order epistasis | | | |
| A×A×A effects | $(\boldsymbol{\alpha\alpha\alpha})_o$ | $\boldsymbol{\tau}_{10o}$ | $(\boldsymbol{\alpha\alpha\alpha})=k^{1/2}_{\alpha\alpha\alpha}(\boldsymbol{\alpha\alpha\alpha})_o$ | $\boldsymbol{\tau}_{10}$, $\binom{m}{3}\times 1$ |
| Model matrix | $\mathbf{W}_{\alpha\alpha\alpha}$ | $\mathbf{W}_{10}$ | $\mathbf{T}_{\alpha\alpha\alpha}=\mathbf{W}_{\alpha\alpha\alpha}/k^{1/2}_{\alpha\alpha\alpha}$ | $\mathbf{T}_{10}$, $n\times\binom{m}{3}$ |
| A×A×A values | $\mathbf{aaa}=\mathbf{W}_{\alpha\alpha\alpha}(\boldsymbol{\alpha\alpha\alpha})_o$ | $\mathbf{u}_{10}$ | $\mathbf{aa}=\mathbf{T}_{\alpha\alpha}(\boldsymbol{\alpha\alpha})$ | $\mathbf{u}_{10}$, $n\times 1$ |
| Variance of A×A×A effects | $\sigma^2_{\alpha\alpha o}$ | $\sigma^2_{10o}$ | $\sigma^2_{\alpha\alpha\alpha}=k_{\alpha\alpha\alpha}\sigma^2_{\alpha\alpha\alpha o}$ | $\sigma^2_{10}$ |
| Variance of A×A×A values | $G^{jj}_{aaa}=\sigma^2_{aaaj}=\sigma^2_{\alpha\alpha\alpha o}\mathbf{W}_{\alpha\alpha\alpha}\mathbf{W}_{\alpha\alpha\alpha}'$ | $G^{jj}_{10}=\sigma^2_{10o}[\mathbf{W}^{(2)}_{10}\mathbf{W}^{(2)}_{10}{}']^{jj}$ | $G^{jj}_{aaa}=\sigma^2_{aaaj}=\sigma^2_{\alpha\alpha\alpha}(\mathbf{S}_{\alpha\alpha\alpha})^{jj}$ | $G^{jj}_{10}=\sigma^2_{10}(\mathbf{S}_{10})^{jj}$ |
| k value | – | – | $k_{\alpha\alpha\alpha}=tr(\mathbf{W}_{\alpha\alpha\alpha}\mathbf{W}_{\alpha\alpha\alpha}')/n$ | $k_{10}$ |
| Genomic relationship matrix | – | – | $\mathbf{S}_{\alpha\alpha\alpha}=\mathbf{T}^{(3)}_{\alpha\alpha\alpha}\mathbf{T}^{(3)}_{\alpha\alpha\alpha}{}'$ | $\mathbf{S}_{10}$, $n\times n$ |
| **G** matrix | $\mathbf{G}_{aaa}=\sigma^2_{\alpha\alpha\alpha o}\mathbf{W}_{\alpha\alpha\alpha}\mathbf{W}_{\alpha\alpha\alpha}'$ | $\mathbf{G}_{10}$ | $\mathbf{G}_{aaa}=\sigma^2_{\alpha\alpha\alpha}\mathbf{S}_{\alpha\alpha\alpha}$ | $\mathbf{G}_{10}$, $n\times n$ |
| A×A×D effects | $(\boldsymbol{\alpha\alpha\delta})^{(3)}_o$ | $\boldsymbol{\tau}_{11o}$ | $(\boldsymbol{\alpha\alpha\delta})^{(3)}_o=k^{1/2}_{\alpha\alpha\delta}(\boldsymbol{\alpha\alpha\delta})^{(3)}_o$ | $\boldsymbol{\tau}_{11}$, $3\binom{m}{3}\times 1$ |
| Model matrix | $\mathbf{W}^{(3)}_{\alpha\alpha\delta}$ | $\mathbf{W}_{11}$ | $\mathbf{T}^{(3)}_{\alpha\alpha\delta}=\mathbf{W}^{(3)}_{\alpha\alpha\delta}/k^{1/2}_{\alpha\alpha\delta}$ | $\mathbf{T}_{11}$, $n\times[3\binom{m}{3}]$ |
| A×A×D values | $\mathbf{aad}^{(3)}=\mathbf{W}^{(3)}_{\alpha\alpha\delta}(\boldsymbol{\alpha\alpha\delta})^{(3)}_o$ | $\mathbf{u}_{11}$ | $\mathbf{aad}^{(3)}=\mathbf{T}^{(3)}_{\alpha\alpha\delta}(\boldsymbol{\alpha\alpha\delta})^{(3)}$ | $\mathbf{u}_{11}$, $n\times 1$ |
| Variance of A×A×D, A×D×A and D×A×A effects | $\sigma^2_{\alpha\alpha\delta o}$ | $\sigma^2_{11o}$ | $\sigma^2_{\alpha\alpha\delta}=\sigma^2_{\alpha\alpha\delta o}$ | $\sigma^2_{11}$ |
| Variance of A×A×D, A×D×A and D×A×A values | $G^{jj}_{aad}=\sigma^2_{aadj}=\sigma^2_{\alpha\alpha\delta o}[\mathbf{W}^{(3)}_{\alpha\alpha\delta}\mathbf{W}^{(3)}_{\alpha\alpha\delta}{}']^{jj}$ | $G^{jj}_{11}=\sigma^2_{11o}[\mathbf{W}^{(2)}_{11}\mathbf{W}^{(2)}_{11}{}']^{jj}$ | $G^{jj}_{aad}=\sigma^2_{aadj}=\sigma^2_{\alpha\alpha\delta}(\mathbf{S}_{\alpha\alpha\delta})^{jj}$ | $G^{jj}_{11}=\sigma^2_{11}(\mathbf{S}_{11})^{jj}$ |
| k value | – | – | $k_{\alpha\alpha\delta}=tr(\mathbf{W}^{(3)}_{\alpha\alpha\delta}\mathbf{W}^{(3)}_{\alpha\alpha\delta}{}')/n$ | $k_{11}$ |
| Genomic relationship matrix | – | – | $\mathbf{S}_{\alpha\alpha\delta}=\mathbf{T}^{(3)}_{\alpha\alpha\delta}\mathbf{T}^{(3)}_{\alpha\alpha\delta}{}'$ | $\mathbf{S}_{11}$, $n\times n$ |
| **G** matrix | $\mathbf{G}_{aad}=\sigma^2_{\alpha\alpha\delta o}\mathbf{W}^{(3)}_{\alpha\alpha\delta}\mathbf{W}^{(3)}_{\alpha\alpha\delta}{}'$ | $\mathbf{G}_{11}$ | $\mathbf{G}_{aad}=\sigma^2_{\alpha\alpha\delta}\mathbf{S}_{\alpha\alpha\delta}$ | $\mathbf{G}_{11}$, $n\times n$ |
| A×D×D effects | $(\boldsymbol{\alpha\delta\delta})^{(3)}_o$ | $\boldsymbol{\tau}_{12o}$ | $(\boldsymbol{\alpha\delta\delta})^{(3)}=k^{1/2}_{\alpha\delta\delta}(\boldsymbol{\alpha\delta\delta})^{(3)}_o$ | $\boldsymbol{\tau}_{12}$, $3\binom{m}{3}\times 1$ |
| Model matrix | $\mathbf{W}^{(3)}_{\alpha\delta\delta}$ | $\mathbf{W}_{12}$ | $\mathbf{T}^{(3)}_{\alpha\delta\delta}=\mathbf{W}^{(3)}_{\alpha\delta\delta}/k^{1/2}_{\alpha\delta\delta}$ | $\mathbf{T}_{12}$, $n\times[3\binom{m}{3}]$ |

| | | | | |
|---|---|---|---|---|
| A×D×D values | $\mathbf{add}^{(3)} = \mathbf{W}^{(3)}_{\alpha\delta\delta}(\boldsymbol{\alpha\delta\delta})^{(3)}_{o}$ | $\mathbf{u}_{12}$ | $\mathbf{add}^{(3)} = \mathbf{T}^{(3)}_{\alpha\delta\delta}(\boldsymbol{\alpha\delta\delta})^{(3)}$ | $\mathbf{u}_{12}$, n×1 |
| Variance of A×D×D, D×A×D, and D×D×A effects | $\sigma^2_{\alpha\alpha\delta o}$ | $\sigma^2_{12o}$ | $\sigma^2_{\alpha\alpha\delta} = \sigma^2_{\alpha\alpha\delta o}$ | $\sigma^2_{12}$ |
| Variance of A×D×D, D×A×D, and D×D×A values | $G^{jj}_{add} = \sigma^2_{addj} = \sigma^2_{\alpha\delta\delta o}[\mathbf{W}^{(3)}_{\alpha\delta\delta}\mathbf{W}^{(3)}_{\alpha\delta\delta}{}']^{jj}$ | $G^{jj}_{12} = \sigma^2_{12o}[\mathbf{W}^{(2)}_{12}\mathbf{W}^{(2)}_{12}{}']^{jj}$ | $G^{jj}_{add} = \sigma^2_{addj} = \sigma^2_{\alpha\delta\delta}(\mathbf{S}_{\alpha\delta\delta})^{jj}$ | $G^{jj}_{12} = \sigma^2_{12}(\mathbf{S}_{12})^{jj}$ |
| k value | – | – | $k_{\alpha\delta\delta} = tr(\mathbf{W}^{(3)}_{\alpha\delta\delta}\mathbf{W}^{(3)}_{\alpha\delta\delta}{}')/n$ | $k_{12}$ |
| Genomic relationship matrix | – | – | $\mathbf{S}_{\alpha\delta\delta} = \mathbf{T}^{(3)}_{\alpha\delta\delta}\mathbf{T}^{(3)}_{\alpha\delta\delta}{}'$ | $\mathbf{S}_{12}$, n×n |
| **G** matrix | $\mathbf{G}_{add} = \sigma^2_{\alpha\delta\delta o}\mathbf{W}^{(3)}_{\alpha\delta\delta}\mathbf{W}^{(3)}_{\alpha\delta\delta}{}'$ | $\mathbf{G}_{12}$ | $\mathbf{G}_{add} = \sigma^2_{\alpha\delta\delta}\mathbf{S}_{\alpha\delta\delta}$ | $\mathbf{G}_{12}$, n×n |
| D×D×D effects | $(\boldsymbol{\delta\delta\delta})_o$ | $\boldsymbol{\tau}_{13o}$ | $(\boldsymbol{\delta\delta\delta}) = k^{1/2}_{\delta\delta\delta}(\boldsymbol{\delta\delta\delta})_o$ | $\boldsymbol{\tau}_{13}$, $\binom{m}{3}\times 1$ |
| Model matrix | $\mathbf{W}_{\delta\delta\delta}$ | $\mathbf{W}_{13}$ | $\mathbf{T}_{\delta\delta\delta} = \mathbf{W}_{\delta\delta\delta}/k^{1/2}_{\delta\delta\delta}$ | $\mathbf{T}_{13}$, $n\times\binom{m}{3}$ |
| D×D×D values | $\mathbf{ddd} = \mathbf{W}_{\delta\delta\delta}(\boldsymbol{\delta\delta\delta})_o$ | $\mathbf{u}_{13}$ | $\mathbf{ddd} = \mathbf{T}_{\delta\delta\delta}(\boldsymbol{\delta\delta\delta})$ | $\mathbf{u}_{13}$, n×1 |
| Variance of D×D×D effects | $\sigma^2_{\delta\delta\delta o}$ | $\sigma^2_{13o}$ | $\sigma^2_{\alpha\alpha\delta} = \sigma^2_{\alpha\alpha\delta o}$ | $\sigma^2_{13}$ |
| Variance of D×D×D values | $G^{jj}_{ddd} = \sigma^2_{dddj} = \sigma^2_{\delta\delta\delta o}[\mathbf{W}_{\delta\delta\delta}\mathbf{W}_{\delta\delta\delta}{}']^{jj}$ | $G^{jj}_{13} = \sigma^2_{13o}[\mathbf{W}^{(2)}_{13}\mathbf{W}^{(2)}_{13}{}']^{jj}$ | $G^{jj}_{ddd} = \sigma^2_{dddj} = \sigma^2_{\delta\delta\delta}(\mathbf{S}_{\delta\delta\delta})^{jj}$ | $G^{jj}_{13} = \sigma^2_{13}(\mathbf{S}_{13})^{jj}$ |
| k value | – | – | $k_{\delta\delta\delta} = tr(\mathbf{W}_{\delta\delta\delta}\mathbf{W}_{\delta\delta\delta}{}')/n$ | $k_{13}$ |
| Genomic relationship matrix | – | – | $\mathbf{S}_{\delta\delta\delta} = \mathbf{T}_{\delta\delta\delta}\mathbf{T}_{\delta\delta\delta}{}'$ | $\mathbf{S}_{13}$, n×n |
| **G** matrix | $\mathbf{G}_{ddd} = \sigma^2_{\delta\delta\delta o}\mathbf{W}_{\delta\delta\delta}\mathbf{W}_{\delta\delta\delta}{}'$ | $\mathbf{G}_{13}$ | $\mathbf{G}_{ddd} = \sigma^2_{\delta\delta\delta}\mathbf{S}_{\delta\delta\delta}$ | $\mathbf{G}_{13}$, n×n |
